# Diurnal metabolic regulation of isoflavones and soyasaponins in soybean roots

**DOI:** 10.1101/2020.04.22.056598

**Authors:** Hinako Matsuda, Masaru Nakayasu, Yuichi Aoki, Shinichi Yamazaki, Atsushi J. Nagano, Kazufumi Yazaki, Akifumi Sugiyama

## Abstract

Isoflavones and soyasaponins are major specialized metabolites accumulated in soybean roots and secreted into the rhizosphere. Unlike the biosynthetic pathway, the transporters involved in metabolite secretion remain unknown. The developmental regulation of isoflavone and soyasaponin secretions has been recently reported, but the diurnal regulation of their biosynthesis and secretion still needs to be further studied. To address these challenges, we conducted transcriptome and metabolite analysis using hydroponically grown soybean plants at 6-hour intervals for 48 hours in a 12-h-light/12-h-dark condition. Isoflavone and soyasaponin biosynthetic genes showed opposite patterns in the root tissues; that is, the former genes are highly expressed in daytime, while the latter ones are strongly induced at nighttime. *GmMYB176* encoding a transcription factor of isoflavone biosynthesis was upregulated from ZT0 (6:00 am) to ZT6 (12:00 am), followed by the induction of isoflavone biosynthetic genes at ZT6. The isoflavone aglycone content in the roots accordingly increased from ZT6 to ZT18 (0:00 am), accompanied by an increase in glucoside levels that peaked at ZT0. The isoflavone aglycone content in root exudates was kept consistent throughout the day, whereas that of glucosides increased at ZT6, which reflected the decreased expression of the gene encoding beta-glucosidase involved in the hydrolysis of apoplast-localized isoflavone conjugates. Co-expression analysis revealed that those isoflavone and soyasaponin biosynthetic genes formed separate clusters, which exhibited a correlation to ABC and MATE transporter genes. As summary, the results in this study indicated the diurnal regulation of isoflavone biosynthesis in soybean roots and the putative transporter genes responsible for isoflavone and soyasaponin transport.

## 1 INTRODUCTION

Plants change their metabolisms and physiological functions during the day to adapt to external biotic and abiotic stresses (Grundy et al., 2015; Seo and Mas, 2015; Lu et al., 2017; Gil and Park, 2019). Specialized metabolites play important roles in adapting to the diurnally changing external environments. The glucosinolate content in the leaves of Arabidopsis (*Arabidopsis thaliana*) and cabbage (*Brassica oleracea*) peaks in the morning and decreases in the evening to midnight to protect their leaves against pests, such as the cabbage looper (*Trichoplusia ni*), which is active during the daytime (Goodspeed et al., 2013). In an iron-deficient condition, barley (*Hordeum vulgare*) roots secrete mugineic acids, which are iron-chelating phytosiderophores, 2–3 hours after dawn to presumably avoid the degradation by microbes (Takagi et al., 1984; Römheld and Marschner, 1990; Nagasaka et al., 2009). The flowers of the rose plant (*Rosa hybrida L.*) secrete geranyl acetate and germacrene D before and at dawn to attract pollinators that are active at dawn (Hendel-Rahmanim et al., 2007).

Transcription factors, such as circadian clock associated 1 (CCA1), late elongated hypocotyl (LHY), timing of CAB expression 1 / pseudo-response regulator 1 (TOC1 / PRR1), pseudo-response regulator 3/5/7/9 (PRR3/5/7/9), REVEILLE 4/6/8 (RVE4/6/8), and night-light-inducible and clock-regulated 1/2 (LNK1/2), coordinately regulate the plant circadian clock (Srivastava et al., 2019). These transcription factors also regulate specialized metabolism (Nguyen and Lee, 2016). For example, RVE8 positively regulates anthocyanin biosynthesis (Pérez-García et al., 2015), and PRR5/7/9 negatively regulates carotenoid and abscisic acid biosynthesis in Arabidopsis (Fukushima et al., 2009). In soybeans, CCA1-like MYB transcription factor GmMYB133 (Glyma.07G066100) stimulates isoflavone biosynthesis by inducing *chalcone synthase 8* (*CHS8*) and *isoflavone synthase 2* (*IFS2*) (Bian et al, 2018), suggesting the diurnal regulation of isoflavone biosynthesis.

Isoflavones are major specialized metabolites in soybeans and function as phytoalexins in defense against pathogens (Subramanian et al., 2005). Isoflavones work not only inside a plant but also in the rhizosphere, a soil region close to the roots. Daidzein, the major isoflavone secreted from soybean roots, acts as a signal for the nodulation (Kosslak et al., 1987) and also modulates the rhizosphere microbiota (Okutani et al., 2020). In addition to isoflavones, soybean roots secrete an equivalent amount of soyasaponins, which was first demonstrated by Tsuno et al. (2018). The secretion of these specialized metabolites changes dramatically in quality and quantity depending on the developmental stage of the soybean plant (Sugiyama et al., 2016; Tsuno et al. 2018); however, the diurnal regulation of these soybean metabolites remains to be described.

Isoflavone and soyasaponin biosynthesis occurs in the cytosol (Figure 1) (Augustin et al., 2011; Nakayama et al., 2019). Conceivably, both isoflavones and soyasaponins accumulate in the vacuoles as glycosides and are secreted *via* transporters (Figure 1) (Yazaki et al., 2008; Mylona, et al., 2008; Sawai and Saito, 2011; Yoo et al., 2013; de Brito Francisco and Martinoia, 2018; Sugiyama 2019). ATP-binding cassette (ABC) and multidrug and toxic compound extrusion (MATE) transporters are involved in the intracellular transport of isoflavones. For example, in *Medicago truncatula*, MtMATE1 and MtMATE2 are responsible for the transport of isoflavone glycosides, such as daidzin and genistin, into the vacuole (Zhao and Dixon, 2009; Zhao et al., 2011). Biochemical analysis has suggested the involvement of ABC-type transporters and apoplast-localized isoflavone conjugate-hydrolyzing beta-glucosidase (ICHG) in isoflavone aglycone secretion to the rhizosphere (Suzuki et al., 2006; Sugiyama et al., 2007), although these processes have not been genetically characterized (Figure 1A). In contrast to isoflavones, the secretory mechanism involving the transporters that are responsible for the vacuolar accumulation and secretion to the rhizosphere are still unknown for soyasaponins, whereas the soyasaponin biosynthetic pathway in soybeans has been intensively studied and characterized (Sundaramoorthy et al., 2019; Krishnamurthy et al., 2019) (Figure 1B).

**Figure 1.**
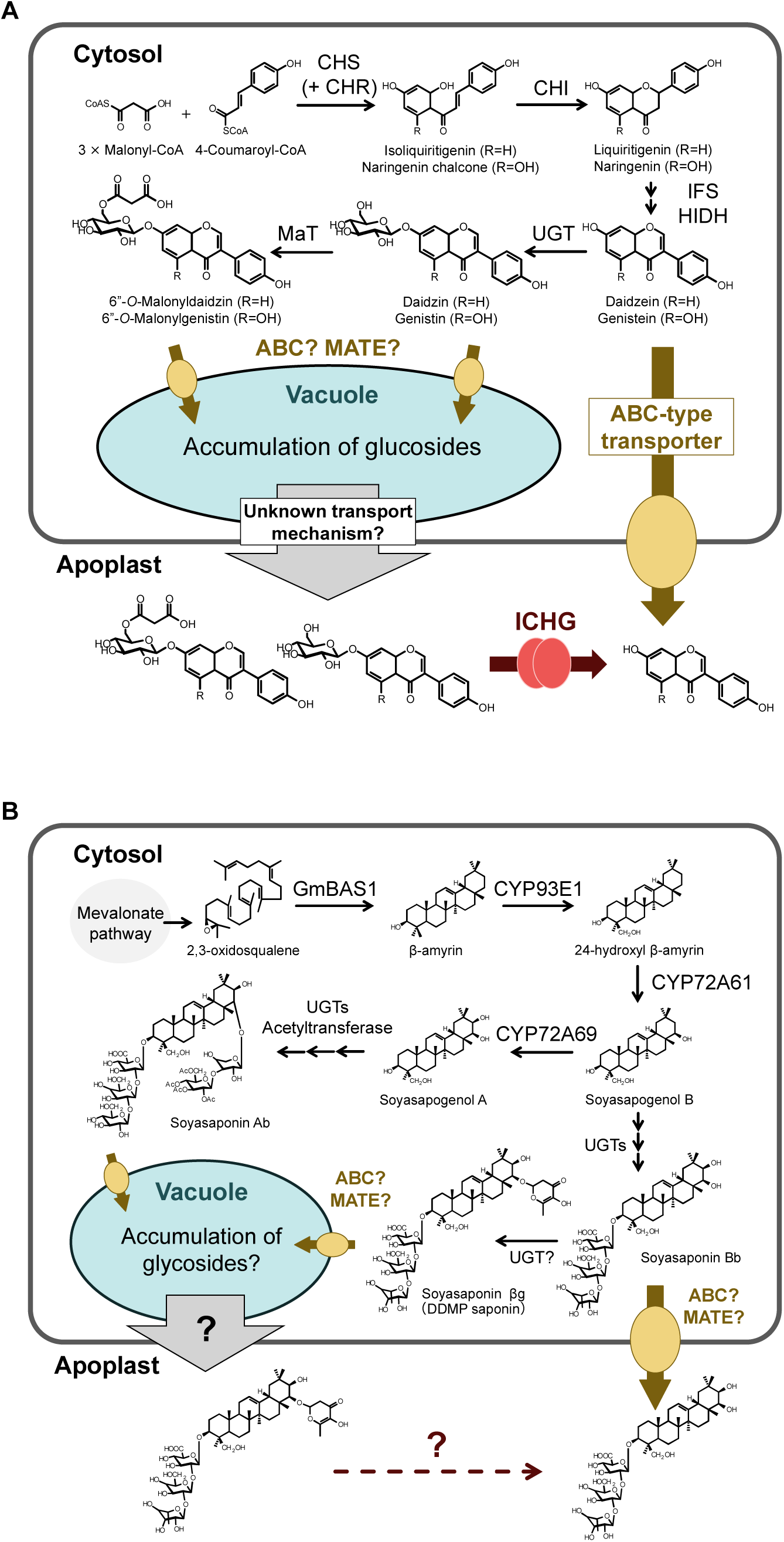
Isoflavone and soyasaponin biosynthesis, and their proposed secretion pathways in soybean root. (A) Isoflavones and (B) soyasaponins. Abbreviations: CHS, Chalcone synthase; CHR, Chalcone reductase; CHI, Chalcone isomerase; IFS, Isoflavone synthase; HIDH, 2-hydroxyisoflavanone dehydratase; UGT, UDP-glucuronosyltransferase; MaT, Malonyl transferase; ICHG, Isoflavone conjugate-hydrolyzing beta-glucosidase; ABC, ATP-binding cassette transporter; MATE, Multidrug and toxic compound extrusion transporter; BAS1, beta-amyrin synthase 1; CYP, Cytochrome P450 (CYP).

In this study, we performed transcriptomic and metabolic analyses using hydroponically grown soybean plants to characterize the diurnal regulation of biosynthesis and secretion of two major classes of specialized metabolites in soybeans. We also narrowed down candidate transporter genes that are responsible for accumulation and secretion of isoflavone and soyasaponin through co-expression network analysis, which highlighted the members of ABC and MATE transporters that reveal a tight correlation with the biosynthetic gene expression pattern of isoflavone and soyasaponin.

## 2 MATERIALS AND METHODS

### 2.1 Chemicals

Malonyldaidzin and malonylgenistin were purchased from Nagara Science (Gifu, Japan). Soyasaponin Ab, soyasapogenol A, soyasapogenol B were purchased from Funakoshi (Tokyo, Japan). Soyasaponin Bb was purchased from ChromaDex (Irvine, CA, US). The other chemicals were obtained from Wako Pure Chemical Industries Ltd. (Osaka, Japan) or Nacalai Tesque Inc. (Kyoto, Japan), unless otherwise stated.

### 2.2 Plant materials and growth conditions

The soybean seeds (cv. Enrei) used in the study were obtained from Tsurushin Shubyo (Matsumoto, Japan). The growth condition of the soybeans in hydroponic cultures was set up according to the description of Sugiyama et al., 2016. After 7 days of the growth in autoclaved vermiculite containing water at 25 °C with 16/8-hour photoperiods, the seedlings were rinsed and transferred to a hydroponic culture system where the soybeans were grown in 450-mL plastic containers filled with a mineral nutrient medium consisting of 3.0 mM MgSO_4_, 6.3 mM KNO_3_, 0.87 mM KCl, 1.4 mM KH_2_PO_4_, 2.5 mM Ca(NO_3_)_2_, 21 μM Fe–EDTA, 4.5□μM KI, 28 μM MnCl_2_, 19 μM H_3_BO_3_, 2.3 μM ZnSO_4_, 0.5 μM CuSO_4_, and 0.003 μM Na_2_MoO_4_, with pH 6.0. The soybeans were kept in a cultivation room set at 25 °C with a 12-hour light/dark cycle. After 2 weeks, the plants were transferred to a new medium 6 hours before sampling for analysis. The leaf and root tissues and root exudates were sampled at ZT0 (6:00 am), ZT6 (0:00 pm), ZT12 (6:00 pm), and ZT18 (0:00 am) for 48 hours with five biological replicates.

### 2.3 RNA extraction and transcriptome sequencing

The total RNA was derived from soybean leaves and roots using RNeasy Plant Mini Kits (Qiagen, CA) according to the manufacturer’s instruction. The DNA in each total RNA sample was digested using DNase I (RNase-free DNase sets, Qiagen, CA). The RNA-seq library was prepared through the Lasy-Seq v1.1 protocol (Kamitani et al., 2019; https://sites.google.com/view/lasy-seq/) using 500 ng total RNA. The library was sequenced by paired-end 150bp + 150bp mode of HiSeqX platform (Illumina, CA).

### 2.4 Transcriptome data analysis

The raw-reads data were quality controlled by removing low-quality bases using Trimmomatic (Bolger et al., 2014) with default parameters. The trimmed reads were aligned to the soybean genome (Glycine_max_v2.1 assembly) (Schmutz et al., 2010) using STAR v2.7.0f (Dobin et al., 2013) based on Ensembl Plants release 43 (Monaco et al., 2014) gene annotations. Gene expression levels were estimated as transcripts per million (TPM) (Wagner et al., 2012) using RSEM v1.3.1 (Li et al., 2011) with default parameters. The transcriptome data set supporting the results of this study was submitted to the DNA Data Bank of Japan (https://www.ddbj.nig.ac.jp) and to be publicly available. Principal component analysis (PCA) was performed using the whole transcriptome data set (30,362 genes; TPM > 1) after removing low-expression genes. The differentially expressed genes (DEGs) between two consecutive sampling points were determined using the DESeq2 package (Love et al., 2014) in the R environment with a false discovery rate (FDR) cutoff of 1%. Gene Ontology (GO) enrichment analysis of gene set was performed using the SoyBase GO Term Enrichment Tool (http://www.soybase.org) in accordance with a diurnal pattern of interests. Gene IDs for isoflavone and soyasaponin biosynthesis, ABC and MATE transporters and ICHG were collected based on the procedures described in previous literatures (Yoo et al., 2013; Liu et al., 2016; Ahmad et al., 2017; Krishnamurthy et al., 2019; Mishra et al., 2019; Sundaramoothy et al., 2019). The networks were constructed using the network visualization software Cytoscape (v. 3.7.2; Shannon et al., 2003).

### 2.5 Preparation of root extracts and exudates

The preparation of root extracts and exudates was performed according to the procedure previously described by Sugiyama et al. (2016). The medium containing root exudates was filtered through Omnipore membrane filters (Millipore, Darmstadt, Germany). The medium was passed through a Sep-pak C18 Plus short cartridge (Waters, MA), which was eluted with 2 mL MeOH. The eluant was dried under nitrogen and reconstituted in 50 µL MeOH for LC-MS/MS analysis.

### 2.6 LC-MS/MS analysis

The samples were separated using an ACQUITY UPLC BEH C18 Column (2.1 × 50 mm, 1.7 µm, Waters) on an LC system (Acquity H-Class System, Wnters). The LC mobile phase consisted of (C) water containing 0.1% (v/v) formic acid and (D) acetonitrile. The gradient program was isocratic at 10% D, Initial; linear 1at 0–85% D, 0–15 min; isocratic at 100 % D, 15–16 min; and isocratic at 100 % D, 16–20.5 min. The injection volume of each sample was 5 µl, and the flow rate was 0.2 ml min. The isolated samples were detected using a tandem quadrupole MS (Xevo TQ-S, Waters) in the Multiple Reaction Monitoring (MRM) mode. The MRM conditions for the respective compounds are listed in Supplementary Table S1.

### 2.7 Statistical analysis

Statistical differences were calculated using the Tukey’s HSD test at p ≤ 0.05 implemented in R (v. 3.6.1; R Core Team, 2019). The outliers (defined as > 1.5*IQR) in the boxplots were excluded from the calculated data.

## Supplementary materials

The following supplemental materials have been provided:

Supplemental Figure S1. Principal component analysis (PCA) for the leaf and root transcriptomes.

Supplemental Figure S2. The 12 different diurnal patterns set for this study (A) and gene expression variations in the leaves (B) and roots (C). (B, C) The shaded areas show nighttime. The data points indicate the average of expression levels of five replicates.

Supplemental Figure S3. The diurnal variations of MYB transcription factors regulating isoflavone biosynthesis in soybean roots. Each boxplot was constructed by ten replicates (n = 5, 2 time points). The individual dots indicate raw data. The outliers were identified with the 1.5*IQR (interquartile range) rule. Tukey’s HSD test was used for statistical analysis (p < 0.05). ZT, hours after dawn. Abbreviations: TPM, Transcripts per million.

Supplemental Figure S4. Co-expression network analysis in soybean roots. Isoflavone and soyasaponin biosynthetic genes, ICHG, ABC, and MATE transporters were analyzed. The exhibited relationships have the Spearman’s rank correlation coefficient > 0.7. Light blue square, isoflavone biosynthetic gene; green square, soyasaponin biosynthetic gene; blue triangle, ICHG gene; red circle, ABC transporter gene; orange circle, MATE transporter gene. Abbreviations: ICHG, Isoflavone conjugate-hydrolyzing beta-glucosidase; ABC, ATP-binding cassette transporter; MATE, Multidrug and toxic compound extrusion transporter.

Supplemental Table S1. MRM conditions for LC-MS/MS analysis.

Supplemental Table S2. Genes in accordance with 12 diurnal patterns.

Supplemental Table S3. Co-expressed biosynthetic and transporter genes in soybean roots (Spearman correlation coefficient > 0.7). The genes are exhibited in Figure S4. Pink, ABC transporter; orange, MATE transporter; light blue, isoflavone biosynthesis; blue, ICHG; green, soyasaponin biosynthesis.

## 3 RESULTS AND DISCUSSION

### 3.1 Time-dependent transcriptome analysis

The RNA-seq analysis yielded over 1.5 billion reads from 80 samples, that is, eight time points, two tissues, and five replicates. We conducted PCA and found that the leaf and root transcriptomes were clearly separated (Figure S1). To provide an overview of the diurnal changes of gene expressions in soybean, diurnal transcription patterns were examined by extracting DEGs that fit 12 diurnal patterns (Figure S2A). A total of 1,073 and 197 genes were found in the leaves and roots, respectively (Table 1, S2, Figure S2B, S2C). Most of the DEGs in leaves belonged to pattern 2 and 6, which showed the expression induction from midnight to dawn and its reduction from dawn to evening. In Table 1, the number of genes in the roots was lower than those in the leaves except for pattern 3, in which the expression peaked at ZT (hours after dawn) 6 (12:00 am).

**Table 1.** Gene numbers in the respective 12 different diurnal patterns in soybean leaves and roots. All genes in this table are listed in Table S2.

| Pattern | # Leaf Genes | # Root Genes |
| --- | --- | --- |
| 1 | 6 | 0 |
| 2 | 219 | 12 |
| 3 | 64 | 76 |
| 4 | 39 | 5 |
| 5 | 140 | 11 |
| 6 | 207 | 39 |
| 7 | 143 | 22 |
| 8 | 54 | 5 |
| 9 | 34 | 0 |
| 10 | 23 | 0 |
| 11 | 110 | 2 |
| 12 | 34 | 25 |

The expression of key genes related to the circadian clock in soybeans (Locke et al., 2018) showed divergent diurnal changes both in the leaves and roots. Late elongated hypocotyl/circadian clock associated 1-like 1 (LCL1, *Glyma07G048500*) belonged to pattern 5 in both the leaves and roots (Figure 2). PRR9 (*Glyma.06G136600*) belonged to pattern 7 and 3 in the leaves and roots, respectively (Figure 2). LCL2 (*Glyma.03G261800*), PRR3 (*Glyma.U034500*), PRR7 (*Glyma.10G048100*), and TOC1 (*Glyma.04G166300*) showed diurnal variation only in the leaves, whereas LCL2 belonged to pattern 5, PRR3/7 belonged to pattern 3, and TOC1 belonged to pattern 8 (Figure 2). These gene expressions in the leaves exhibited the same trends as previous studies (Marcolino-Gomes et al., 2014; Locke, Slattery, and Ort, 2018). The diurnal fluctuation of the gene expressions for LCL2, PRR3/7, and TOC1 in the roots showed the same pattern as in the leaves, although statistically insignificant. The genes encoding transcription factors GIGANTEA (GI) (*Glyma.20G170000*), Jumonji (*Glyma.11G130600*), ZEITLUPE (ZTL) (*Glyma.09G056100*), LUX ARRHYTHMO (LUX) (*Glyma.12G060200*), and Early flowering 4 (ELF4) (*Glyma.18G027500*) did not fall within the 12 expression patterns but displayed a similar expression tendency as observed in previous studies (Marcolino-Gomes et al., 2014; Locke et al., 2018) (Figure 2).

**Figure 2.**
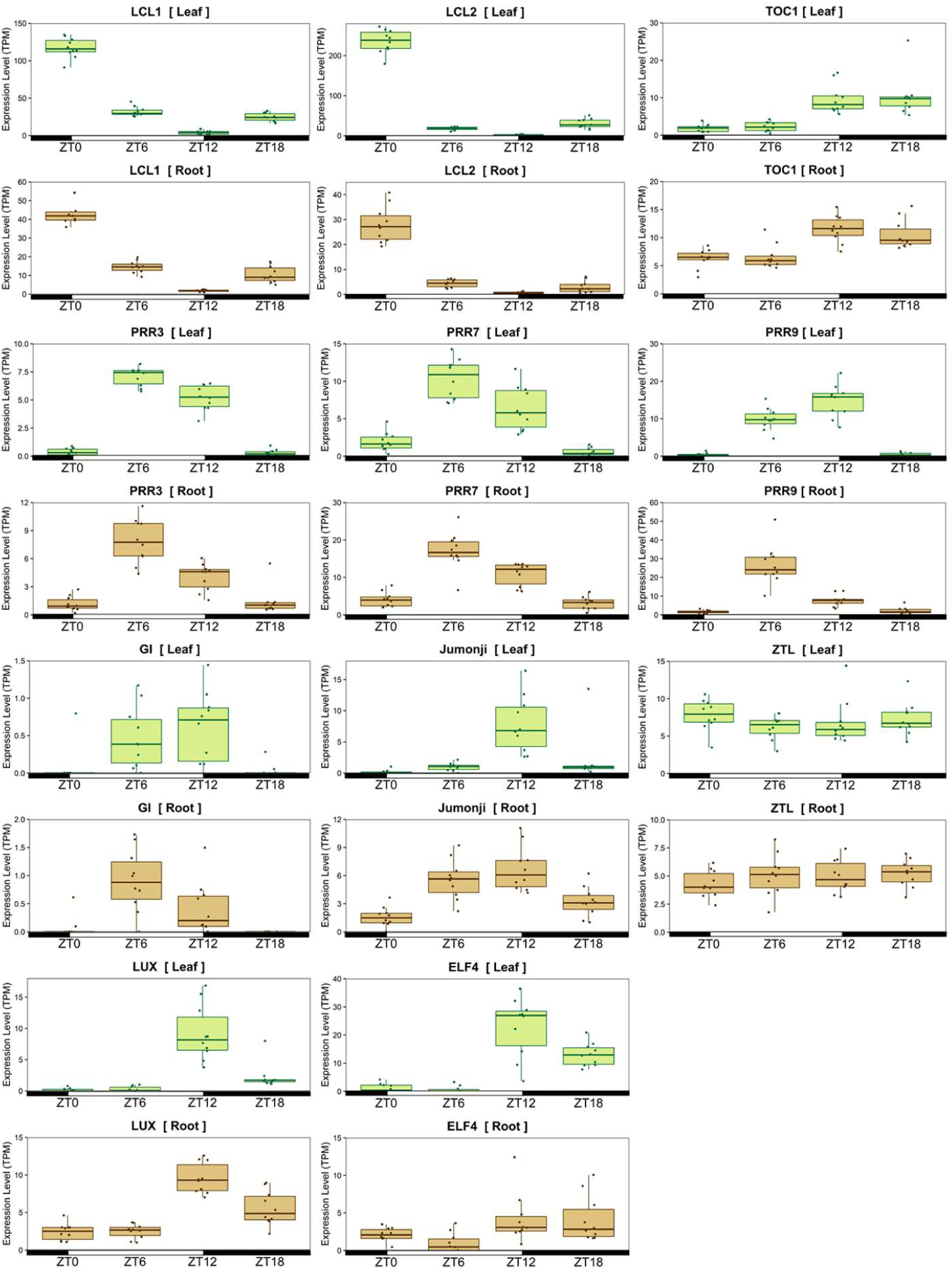
Diurnal variations of circadian rhythm-related genes. Each boxplot was constructed by 10 replicates (n = 5, 2 time points). The individual dots indicate raw data. The outliers were identified using the 1.5*IQR (interquartile range) rule. The green and brown boxplots exhibit gene expression levels in the leaves and roots. ZT, hours after dawn. Abbreviations: TPM, Transcripts per million; LCL, Late elongated hypocotyl/circadian clock associated; TOC, Timing of CAB expression; PRR, Pseudo-response regulator; GI, GIGANTEA; ZTL, ZEITLUPE; LUX, LUX ARRHYTHMO; ELF, Early flowering.

### 3.2 Gene ontology enrichment analysis on 12 diurnal patterns

To understand the characteristic biological processes of the respective variation patterns, we carried out GO enrichment analysis for the genes listed in Supplementary Table 2. In the leaves, the GO terms related to light response and photosynthesis were enriched from midnight to noon. The GO terms associated with ion transport were represented in pattern 6 and had increased expression in the dawn and noon, which consistent with the GO enrichment analysis on diurnal transcripts in rice leaves (Xu et al., 2011). As observed in maize leaves (Feng et al., 2017), the GO terms associated with carbon metabolism and DNA/RNA processing were induced from dawn to dusk. The GO terms linked with plant hormone signal transduction were highlighted from midnight to dawn (Table 2). In the roots, the GO terms associated with light response and photosynthesis were represented in pattern 3, 5, and 6; DNA/RNA processing was represented in pattern 7; and auxin biosynthetic process was represented in pattern 5 (Table 2). The trends in the roots were consistent with those in the leaves.

**Table 2.**
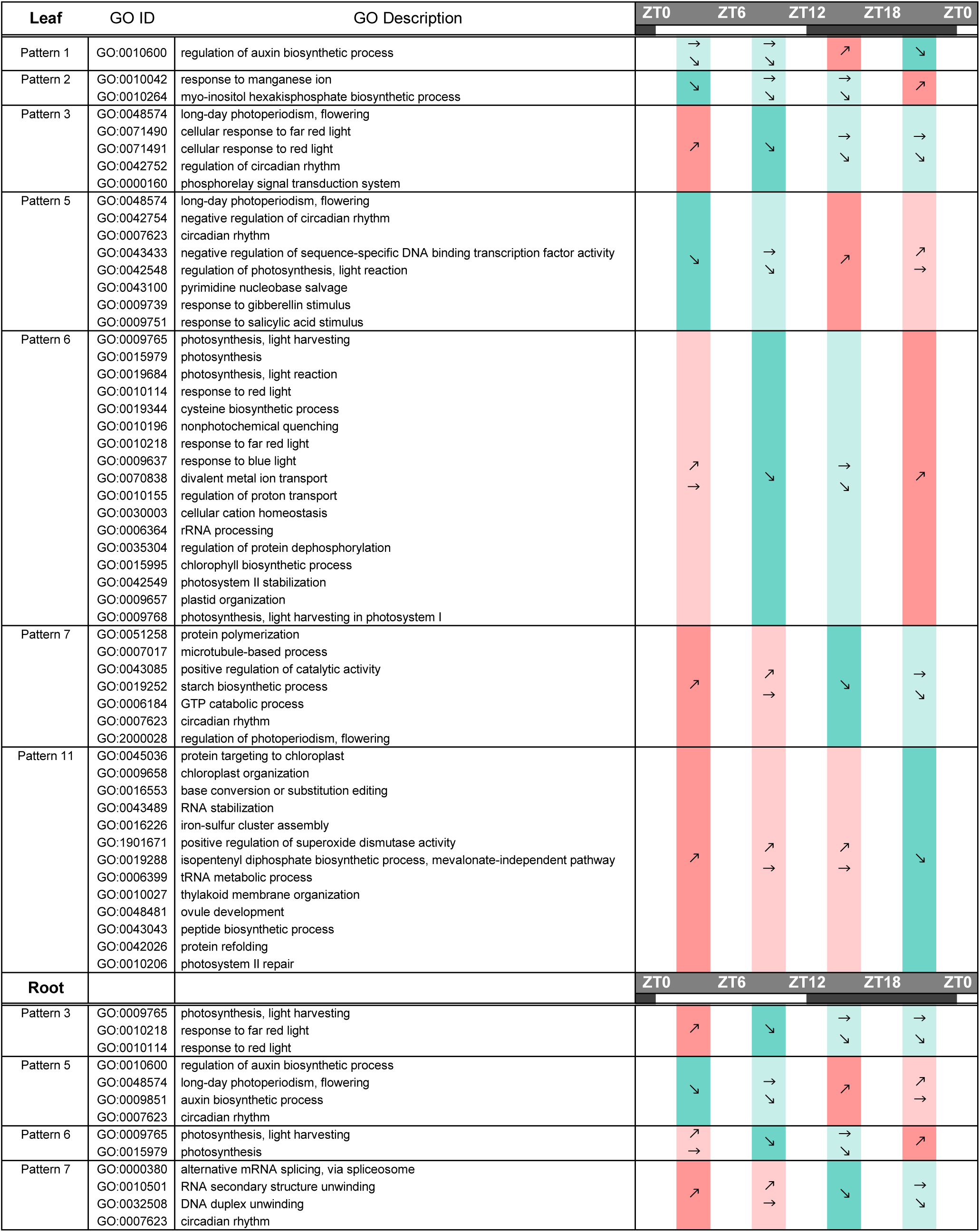
Gene ontology (GO) enrichment analysis for soybean leaf and root genes on 12 diurnal regulation patterns.

The GO terms associated with specialized metabolism were not found in any pattern in the GO enrichment analysis, but the expression of genes involved in isoflavone and soyasaponin biosynthesis displayed diurnal variations (Table S2). Among the genes in isoflavone biosynthetic pathway, *Glyma.14G205200* encoding *trans*-cinnamate 4-monooxygenase belonged to pattern 5, and *Glyma.13G095600* encoding 4-coumarate-CoA ligase 1 and *Glyma.16G219500* encoding chalcone reductase CHR2 belonged to pattern 6 in the leaves. In the roots, *Glyma.11G093100* encoding isoflavone 3’-hydroxylase belonged to pattern 3. In turn, among the genes involved in the soyasaponin biosynthetic pathway, *Glyma.11G112000* and *Glyma.13G253100*, which code for squalene synthase and squalene monooxygenase, respectively, were found in pattern 6, and *Glyma.14G090400* encoding squalene monooxygenase were found in pattern 2. *Glyma.15G221300* encoding soyasapogenol B glucuronide galactosyltransferase-like was listed in pattern 3 in the leaves. No soyasaponin biosynthetic gene was found in any of the 12 variation patterns in the roots.

### 3.3 Diurnal variation of biosynthetic gene expression for isoflavone and soyasaponin

RNA-seq analysis revealed diurnal variations in the gene expression of isoflavone and soyasaponin biosynthetic pathways. To further analyze the diurnal variation patterns, we investigated the fluctuations of gene expression for all the reported genes involved in isoflavone and soyasaponin biosynthesis. No diurnal variation pattern common to the isoflavone or soyasaponin biosynthetic genes were observed in the leaves (Figure 3). In contrast, the expression profiles in the roots exhibited clear diurnal variation patterns for the genes involved in isoflavone and soyasaponin biosynthesis (Figure 3). Isoflavone biosynthetic genes, such as chalcone reductase (CHR), chalcone synthase (CHS), isoflavone synthase (IFS), and 2-hydroxyisoflavanone dehydratase (HIDH) showed high expression at ZT6 and low expression at night. This variation pattern is inconsistent with that of the flavonoid biosynthetic genes in Arabidopsis (Harmer et al., 2000), which were highly expressed at night under constant light conditions after 7 days of culture in a 12-hours light/dark condition.

**Figure 3.**
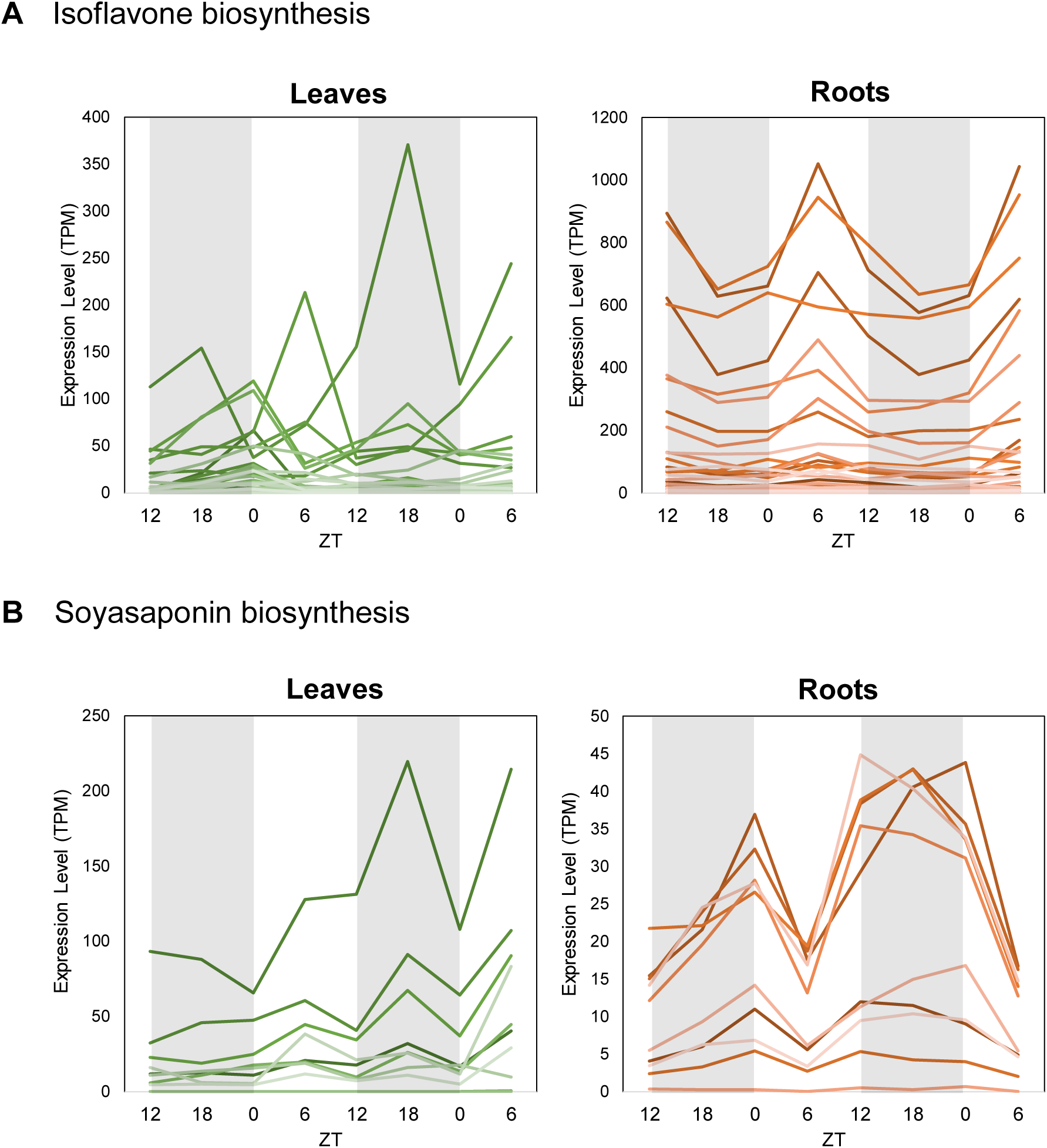
Diurnally differentially expressed genes related to isoflavone and soyasaponin biosynthesis in soybean leaves (green lines) and roots (brown lines). (A) Isoflavone biosynthetic genes described in Figure 1A. Gene IDs were obtained from NCBI (https://www.ncbi.nlm.nih.gov/). (B) Soyasaponin biosynthetic genes listed in Krishnamurthy et al. (2019) and Sundaramoorthy et al. (2019). The shaded areas show nighttime. The data points indicate the average of expression levels of five replicates. ZT, hours after dawn. Abbreviations: TPM, Transcripts per million.

The expression levels of genes encoding β-Amyrin synthase (*BAS*), cytochrome P450 (*CYP*) for triterpenes, and UDP-glucuronosyltransferase (*UGT*) involved in soyasaponin biosynthesis (Sundaramoothy et al., 2019; Krishnamurthy et al., 2019) increased from ZT18 (0:00 am) to ZT0 (6:00 am) and decreased at ZT6, which was an inverse pattern to that of isoflavone biosynthesis.

The transcription factors of the MYB family play crucial roles in the regulation of isoflavone biosynthesis. GmMYB29, GmMYB102, GmMYB133, GmMYB176, GmMYB280, and GmMYB502 are positive regulators of isoflavone biosynthetic genes (Yi et al., 2010; Chu et al., 2017; Bian et al, 2018; Vadivel et al., 2019; Sarkar et al., 2019), while GmMYB29B, GmMYB39, and GmMYB100 negatively regulate those genes (Liu et al., 2013; Yan et al., 2015; Jahan et al., 2020). Among these MYBs, *GmMYB176* showed the highest expression level in the roots (Figure 4, S3). *GmMYB176* exhibited a remarkable diurnal variation, with a higher expression level at ZT0 and ZT6 than at ZT12 (Figure 4), indicating the roles of this MYB member in the induction of isoflavone biosynthetic genes during the daytime.

**Figure 4.**
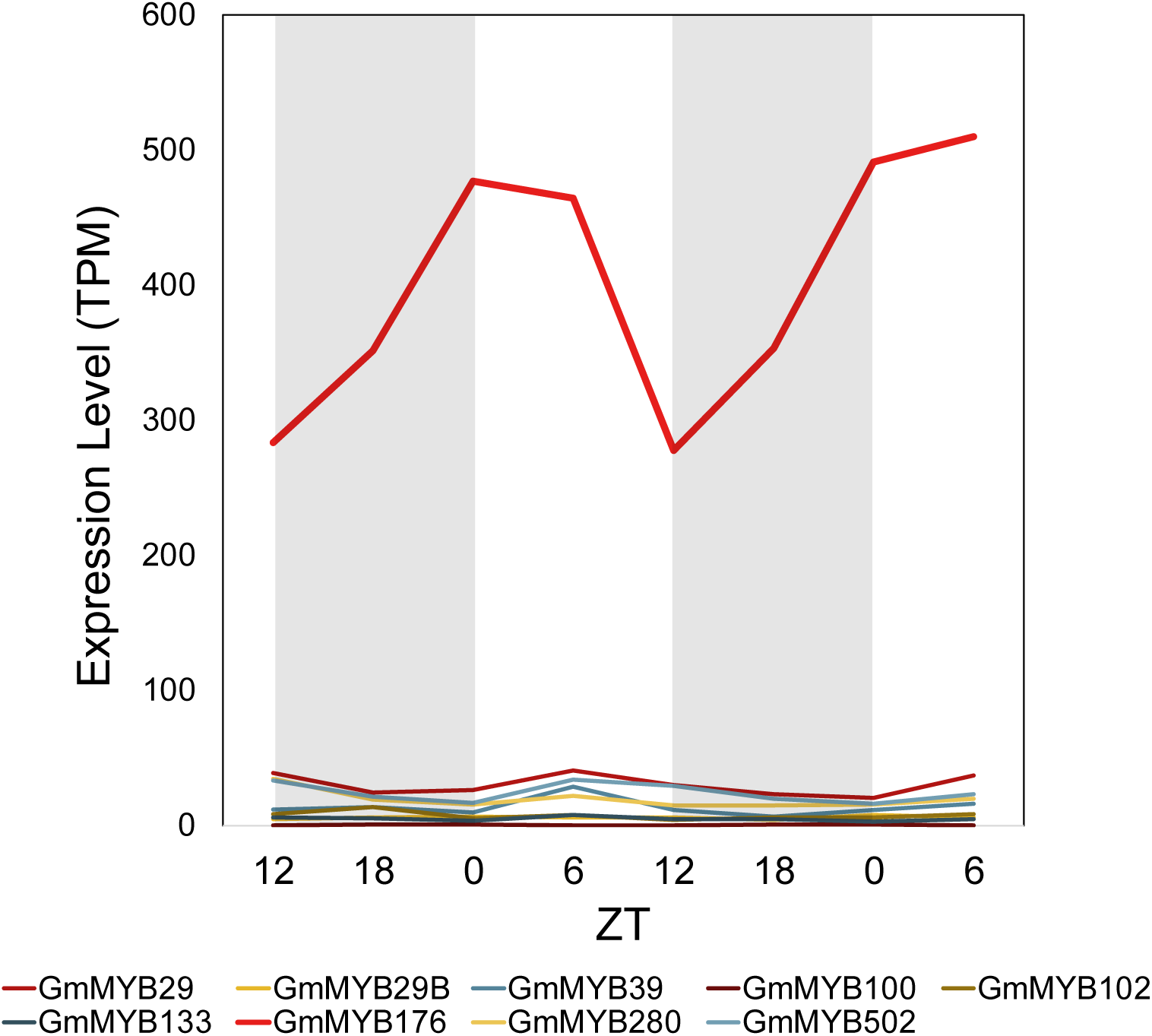
Diurnally differentially expressed genes of MYB transcription factors of isoflavone biosynthesis in soybean roots. The shaded areas show nighttime. The data points indicate the average of expression levels of five replicates. ZT, hours after dawn. Abbreviations: TPM, Transcripts per million.

### 3.4 Co-expression network of isoflavone and soyasaponin metabolism-related genes

The transporters engaged in the accumulation or secretion of both isoflavones and soyasaponins in soybeans have not been reported to date. In this study, co-expression network analysis was performed using a Spearman’s correlation coefficient threshold value of 0.8 to explore the candidate genes responsible for isoflavone and soyasaponin transport. Isoflavone biosynthetic genes displayed a separated co-expression network from those of soyasaponin biosynthesis (Figure 5). On the one hand, the isoflavone biosynthetic gene cluster included three genes coding for ABC transporters (*Glyma.10G019000, Glyma.13G361900*, and *Glyma.15G011900*), and a MATE-type transporter gene *Glyma.01G026200* (Figure 5A, Table 3). On the other hand, the cluster of soyasaponin biosynthetic genes only contained an ABC transporter gene, *Glyma.15G148500* (Figure 5B, Table 3). When co-expression network analysis was performed using a Spearman’s correlation coefficient threshold value of 0.7, the isoflavone biosynthetic gene cluster contained five additional ABC transporter genes, namely *Glyma.01G008200, Glyma.03G101000, Glyma.07G233900, Glyma.13G043800*, and *Glyma.20G242000*, whereas the soyasaponin biosynthetic gene cluster included four more ABC transporter genes, namely *Glyma.04G069800, Glyma.08G101500, Glyma.19G021500*, and *Glyma.19G184300* (Figure S4, Table S3). The correlation analysis using publicly open transcriptome data has recently become available in SoyCSN for soybean (Wang et al., 2019). The co-expression of the ABC transporter genes (*Glyma.10G019000, Glyma.13G043800, Glyma.13G361900*, and *Glyma.15G011900*) with CHS7 (*Glyma.01G228700*) and IFS1 (*Glyma.07G202300*), and the co-expression of two other ABC transporter genes (*Glyma.15G148500* and *Glyma.19G021500*) with BAS1 (*Glyma.07G001300*) and CYP93E1 (*Glyma.08G350800*) were also obtained from SoyCSN (Wang et al., 2019). These findings suggest that the ABC transporter genes are co-expressed with the biosynthetic genes of isoflavone or soyasaponin not only in diurnal variations but also in the symbiosis or other tissues. It is thus expected that those transporters mediate in the vacuolar accumulation and secretion into the rhizosphere of isoflavones or soyasaponins.

**Table 3.**
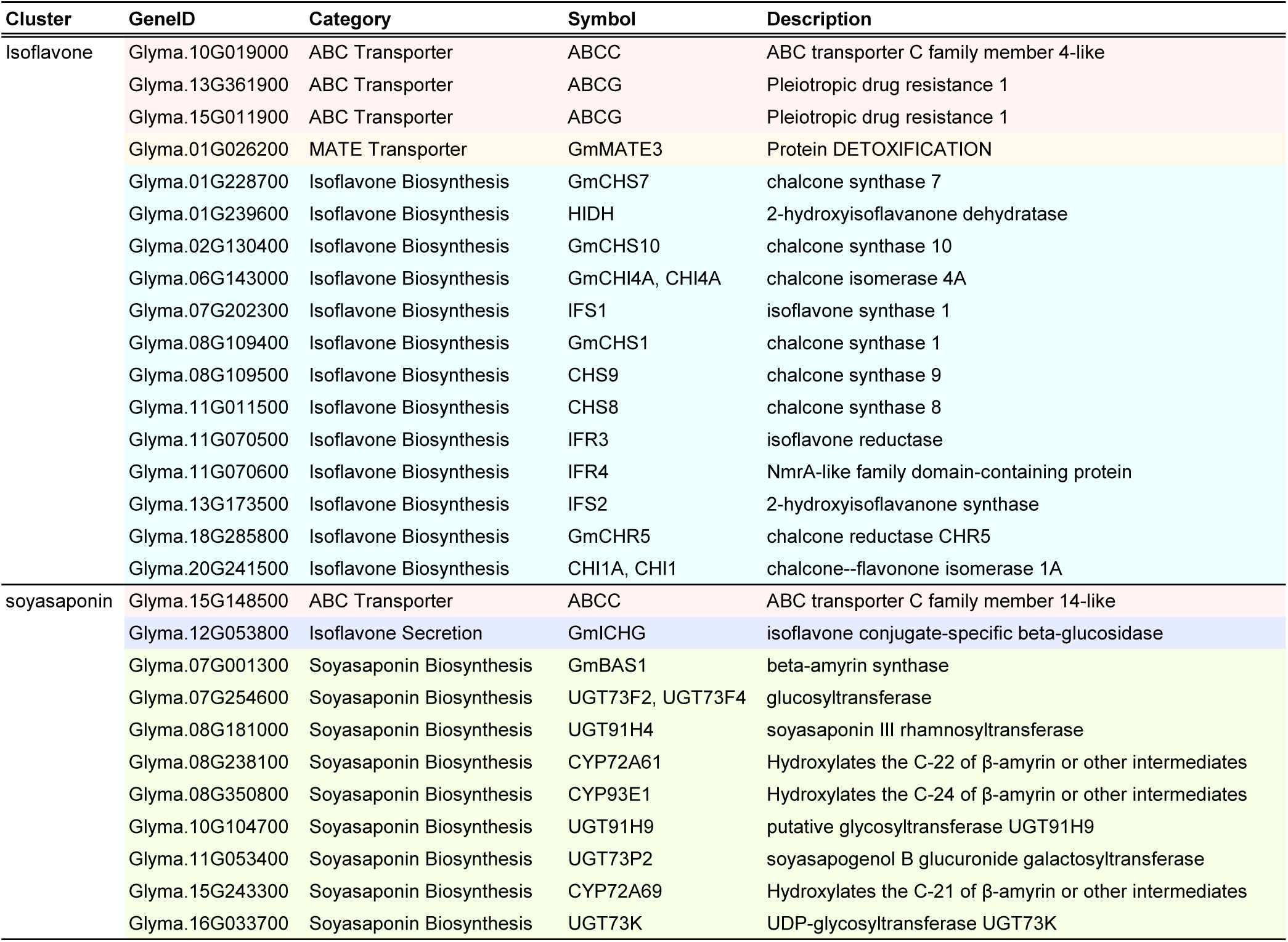
Co-expressed biosynthetic and transporter genes in soybean roots (Spearman correlation coefficient > 0.8). The genes are exhibited in Figure 5. Pink, ABC transporter; orange, MATE transporter; light blue, isoflavone biosynthesis; blue, ICHG; green, soyasaponin biosynthesis.

**Figure 5.**
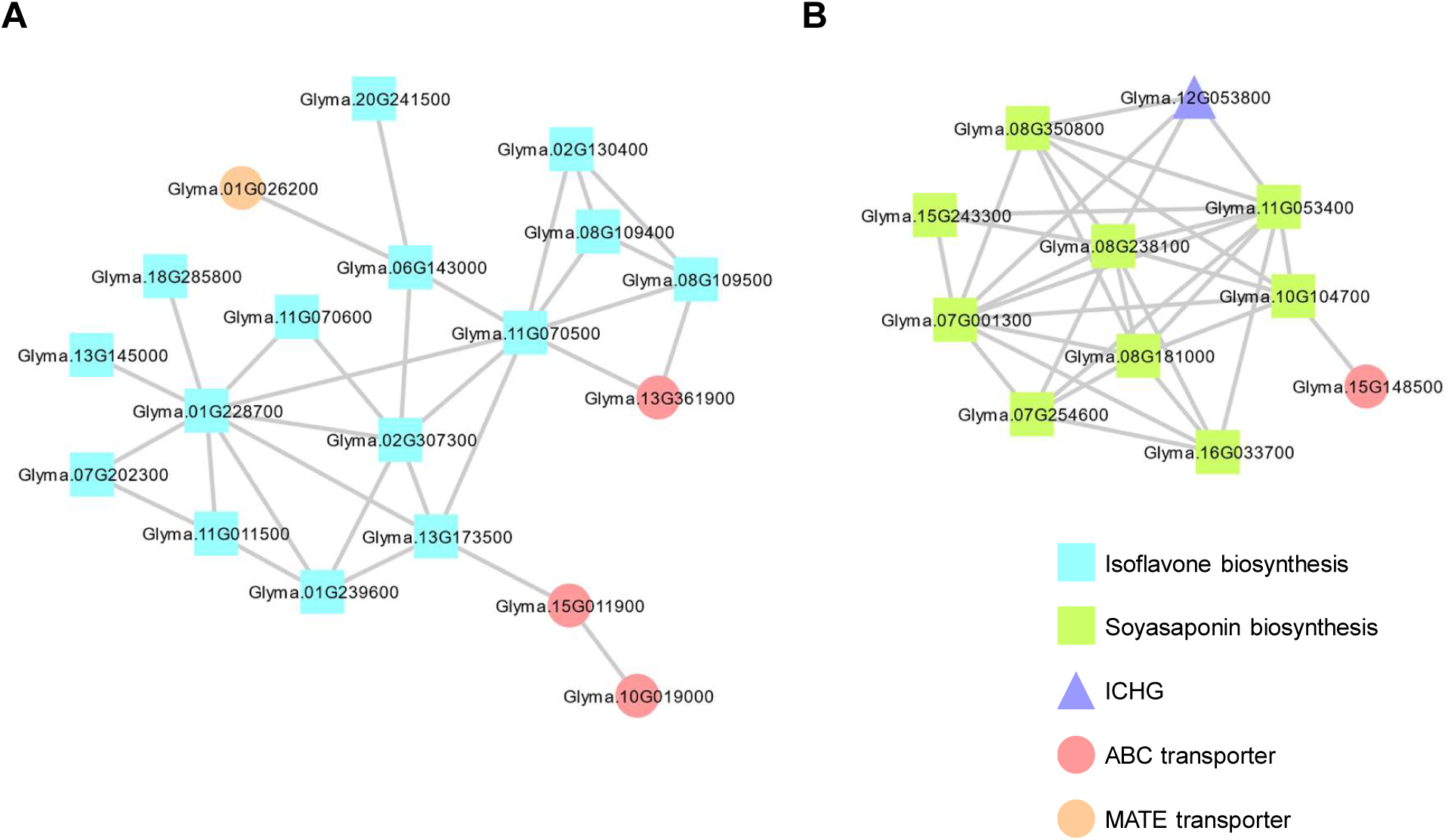
Co-expression network analysis in soybean roots. Isoflavone and soyasaponin biosynthetic genes, ICHG, ABC, and MATE transporters were analyzed. (A) Isoflavone and (B) soyasaponin biosynthetic gene clusters. The exhibited relationships have the Spearman’s rank correlation coefficient > 0.8. Light blue square, isoflavone biosynthetic gene; green square, soyasaponin biosynthetic gene; blue triangle, ICHG gene; red circle, ABC transporter gene; orange circle, MATE transporter gene. Abbreviations: ICHG, Isoflavone conjugate-hydrolyzing beta-glucosidase; ABC, ATP-binding cassette transporter; MATE, Multidrug and toxic compound extrusion transporter.

The coordinated gene expression for biosynthesis and transport has been reported in several studies. For instance, MtMATE1 of *Medicago truncatula* was reported as an epicatechin 3’-*O*-glucoside transporter identified from the co-expression with its biosynthetic genes (Zhao and Dixon, 2009). A glucosinolate transporter 1 (GTR1) in Arabidopsis was identified as jasmonoyl-isoleucine and gibberellin transporter based on its co-expression with jasmonate biosynthetic genes (Saito et al., 2015). Furthermore, *Catharanthus roseus* CrNPF2.9 was identified by co-expression analysis of the biosynthetic genes of monoterpene indole alkaloids (Payne et al., 2017), and saffron (*Crocus sativus*) CsABCC4a was revealed as a transporter for crocins by narrowing down the candidate transporter genes that were highly expressed in pistils (Demurtas et al., 2019). These reports prompted us to genetically and biochemically investigate candidate transporter genes for isoflavone and soyasaponin transport further.

*ICHG* is grouped within the soyasaponin biosynthetic gene cluster, although it hydrolyzes isoflavone glucosides and not soyasaponins, presumably (Suzuki et al., 2006). Nevertheless, the expression level of *ICHG* in the roots from ZT18 to ZT0 was high, whereas it was low at ZT6 (Figure 6), which was consistent with the pattern of the soyasaponin biosynthetic genes (Table 3, Figure 3, 5). This could be a coincidence because *ICHG* is the most downstream gene in the secretion of isoflavones and could be expressed later than those for isoflavone biosynthesis, while there remains a possibility that ICHG hydrolyzes the glucoside linkage of soyasaponins in the apoplast.

**Figure 6.**
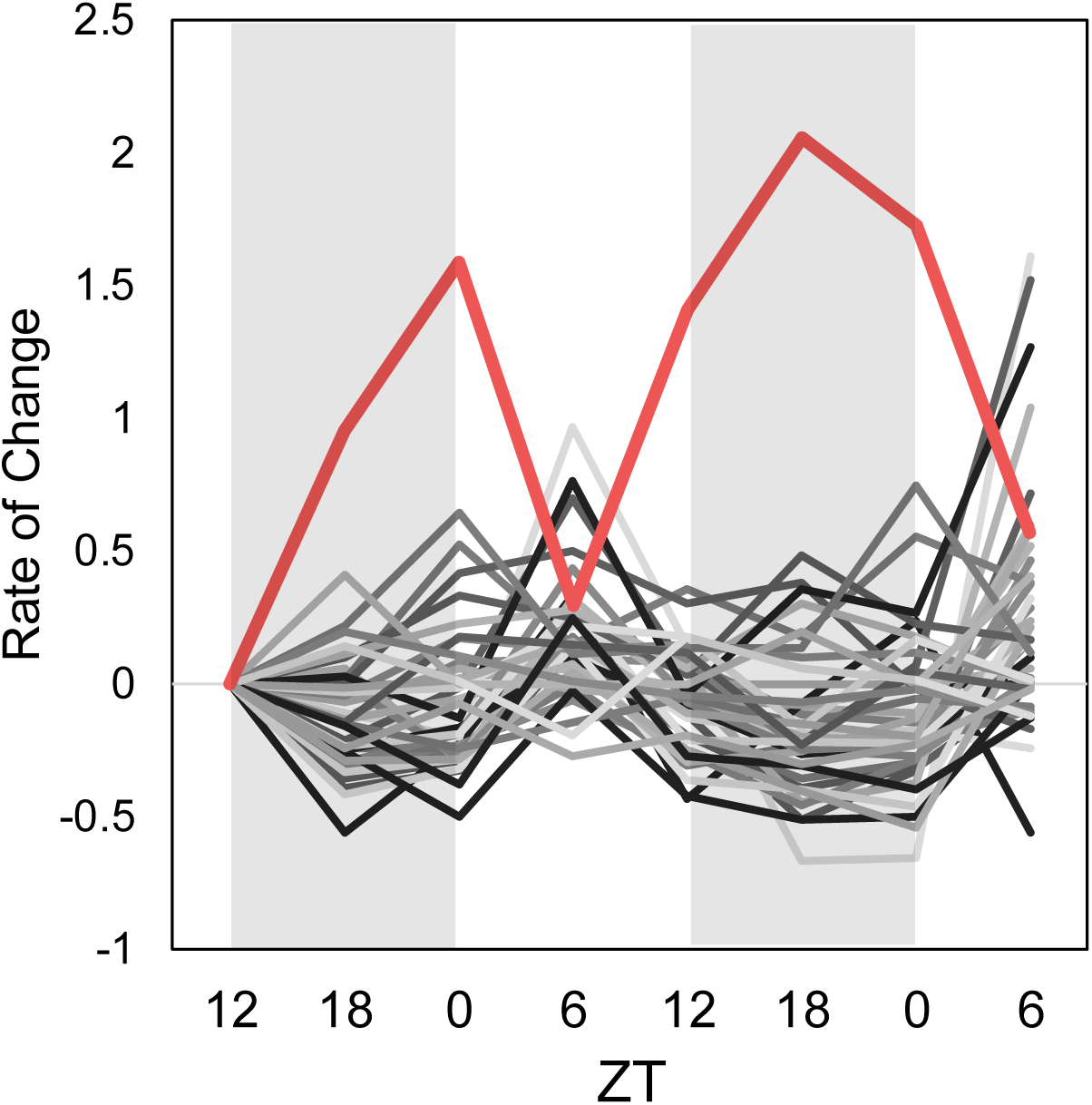
Change rate for ICHG and isoflavone biosynthetic genes in soybean roots. The shaded areas show nighttime. Red lines, ICHG gene. Grey lines, isoflavone biosynthetic genes. The data points indicate average of expression levels of five replicates. ZT, hours after dawn.

### 3.5 Diurnal variation of isoflavones and soyasaponins in roots and root exudates

The gene expression in isoflavone and soyasaponin biosynthesis displayed diurnal variations in the roots. We analyzed the contents of these specialized metabolites in both the roots and root exudates using LC-MS/MS. The daidzein content increased from ZT0 to ZT18 (Figure 7A), followed by the stimulation of isoflavone biosynthetic genes between ZT0 and ZT6 (Figure 4). This temporal difference between gene expression and metabolite accumulation was consistent with the flavonoid biosynthesis in Arabidopsis (Nakabayashi et al. 2017). The contents of its glycosides, daidzin and malonyldaidzin, increased from ZT18 to ZT0. The difference in the variation patterns of aglycone and glycosides in root content was likely due to the biosynthetic order. The sum of daidzein and its glucosides remained constant, suggesting the homeostatic regulation of isoflavone levels in the root throughout the day. The contents of genistein and its glucosides were about 5–10-fold lower than those of daidzein and its glucosides and did not show apparent diurnal variations (Figure 7A). In contrast to the accumulation of isoflavone glucosides in the roots, the aglycone daidzein was the predominant isoflavone form found in root exudates as observed previously (Sugiyama et al. 2016). The amount of daidzein and genistein in root exudates did not show clear diurnal variations (Figure 7B). In contrast, the contents of glucosides, such as daidzin, malonyldaidzin, genistin, and malonylgenistin, were highest during ZT6 to ZT12 (Figure 7B), which is in concordance with the decreased *ICHG* expression at ZT6 (Figure 6). These findings implicate the possible involvement of two modes in the secretion of isoflavones to the rhizosphere, that is, one for the secretion of aglycones and another for the secretion of glucosides, because daidzein content constantly remained at its highest among isoflavones even at ZT6 when *ICHG* expression was suppressed.

**Figure 7.**
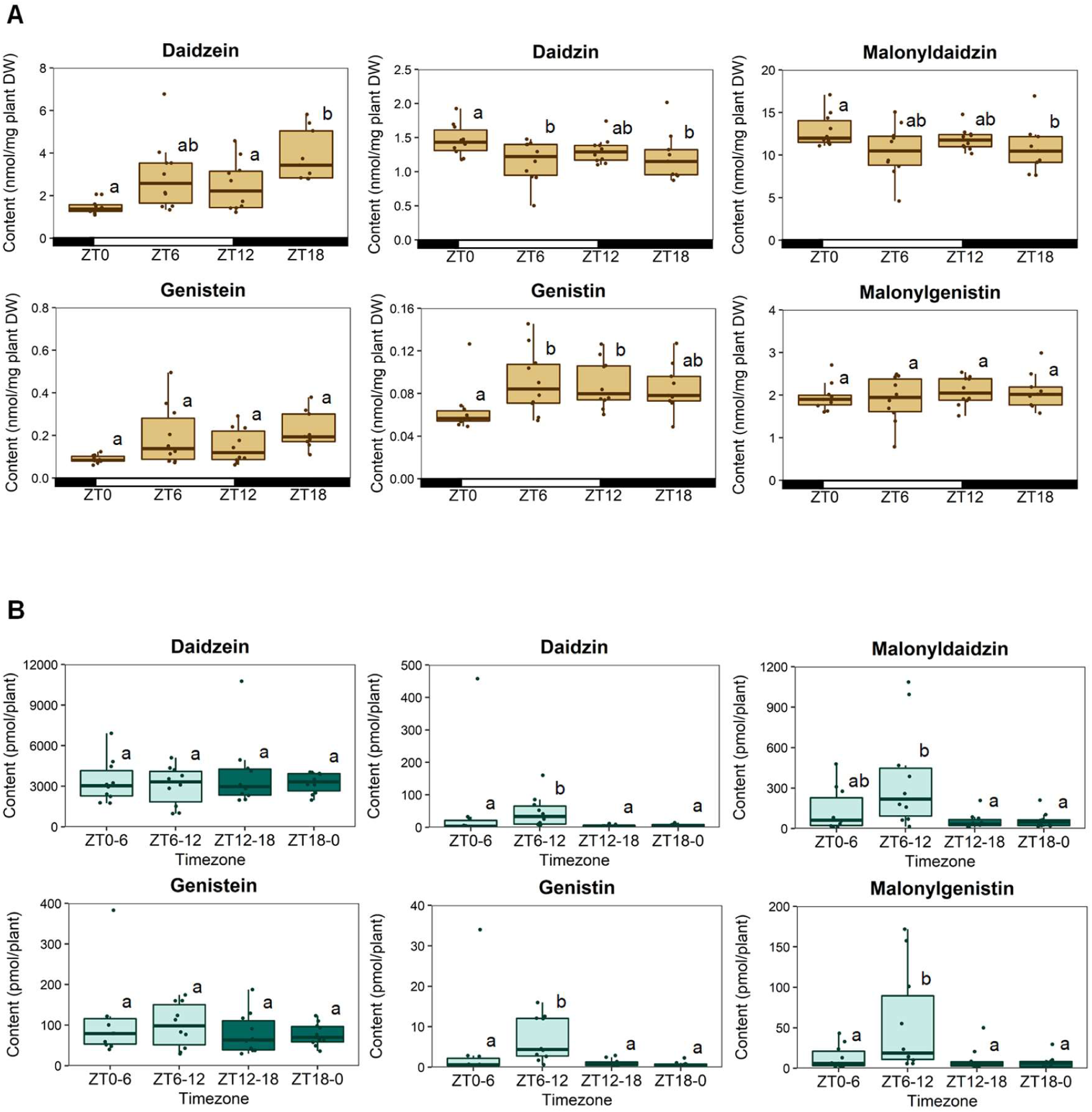
Isoflavone content in soybean roots and root exudates. Isoflavone content in (A) soybean roots and (B) root exudates. The brown boxplots exhibit root content. Pale blue and dark green boxplots show root exudate content in the light and dark time. Each boxplot was constructed by ten replicates (n = 5, 2 time points) except for ZT18 of the root content, which lacked one replicate due to a technical error. The individual dots indicate raw data. The outliers were identified using the 1.5*IQR (interquartile range) rule. Tukey’s HSD test was used for statistical analysis (p < 0.05). ZT, hours after dawn.

We also determined the contents of soyasaponins using LC-MS/MS. We focused on two major soyasaponins in root exudates of soybean, soyasaponin Ab and soyasaponin Bb, and their aglycones, soyasapogenol A and soyasapogenol B (Tsuno et al. 2018), due to the limited availability of authentic samples for soyasaponins. The root content of soyasaponin Ab increased from ZT0 to ZT18, and that of soyasaponin Bb was the least at ZT0 but was significantly increased at ZT6 and maintained until ZT18 (Figure 8A). Soyasaponin Ab and Bb were the major soyasaponins in the root exudates throughout a day, with a slight reduction of soyasaponin Bb from ZT18 to ZT0 (Figure 8B). The amount of soyasaponin aglycones in the roots was much lower than that of their glucosides. Soyasapogenol A was not detected. Soyasapogenol B was present in small quantities, and it was at its highest content at ZT6 and lowest at ZT18 (Figure 8A). The content of soyasapogenol B in the root exudates was highest from ZT0 to ZT6, which would reflect the induction of soyasaponin biosynthetic genes during the night and the slight increase of soyasapogenol B in ZT6 (Figure 8B).

**Figure 8.**
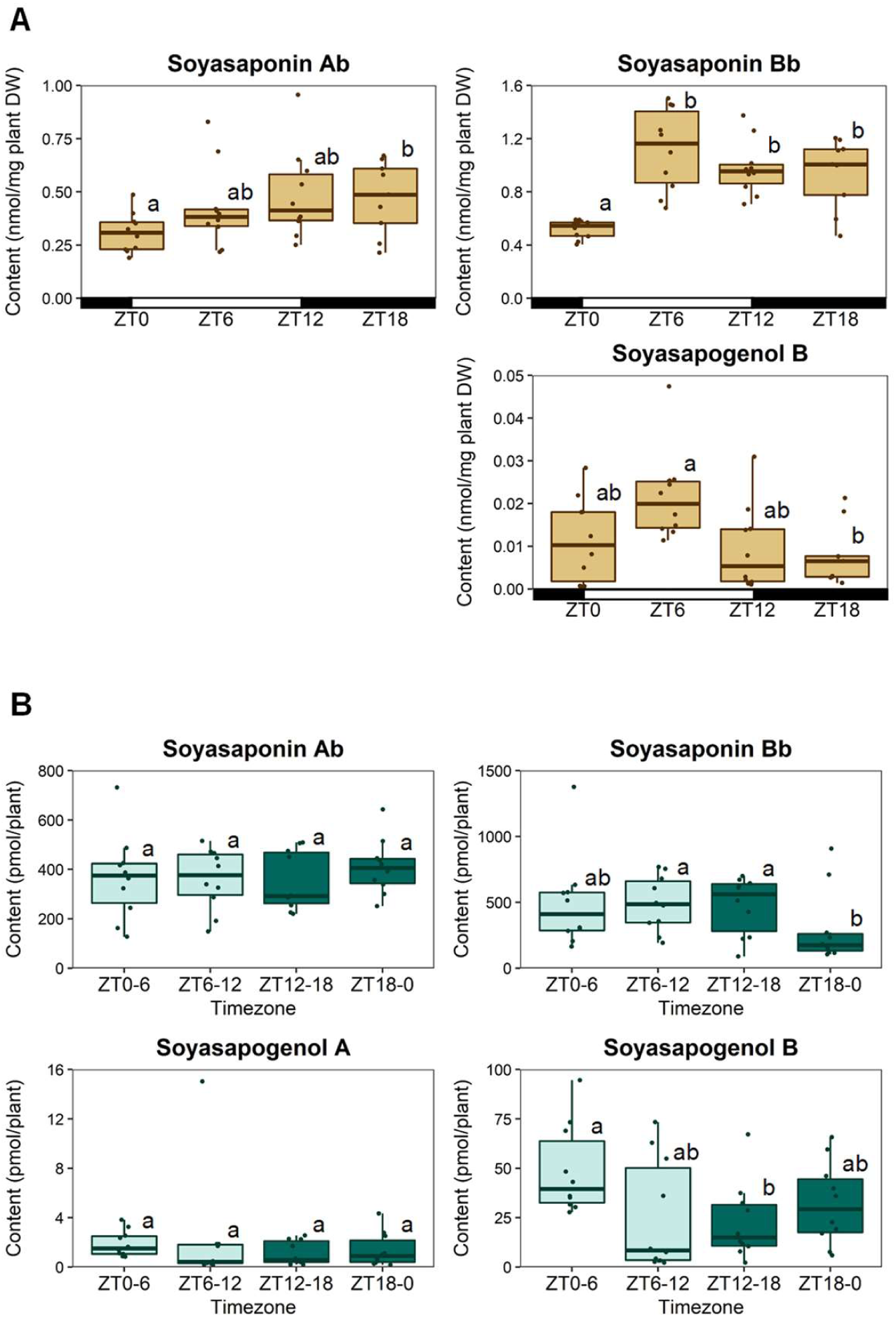
Soyasaponin content in soybean roots and root exudates. The brown boxplots exhibit root content. Pale blue and dark green boxplots show root exudate content in the light and dark time. Soyasaponin content in (A) soybean roots and (B) root exudates. Each boxplot was constructed by ten replicates (n = 5, 2 time points) except for ZT18 of the root content, which lacked one replicate due to a technical error. The individual dots indicate raw data. The outliers were identified using the 1.5*IQR (interquartile range) rule. Soyasapogenol A was not detected in soybean root. Tukey’s HSD test was used for statistical analysis (p < 0.05). ZT, hours after dawn.

## 4 Conclusions

In this study, we elucidated the diurnal variability of isoflavone biosynthesis in soybean roots. GmMYB176, a major transcription factor of isoflavone biosynthesis, stimulates the isoflavone biosynthetic genes from ZT0 to ZT6, followed by the induction of isoflavone biosynthetic genes at ZT6, the increment of daidzein content at ZT18, and the slight increase of its glucosides at ZT0 (Figure 9). In contrast, soyasaponin biosynthetic genes were highly expressed from ZT18 to ZT0. Co-expression network analysis revealed that the clusters for isoflavone and soyasaponin biosynthesis were separated; that is, the former was induced during daytime, and the latter was activated at nighttime. The network analyses highlighted several genes encoding ABC and MATE transporters, which showed closely correlated expression patterns with isoflavone and soyasaponin biosynthetic genes. These genes are promising candidates for further characterization in future studies.

**Figure 9.**
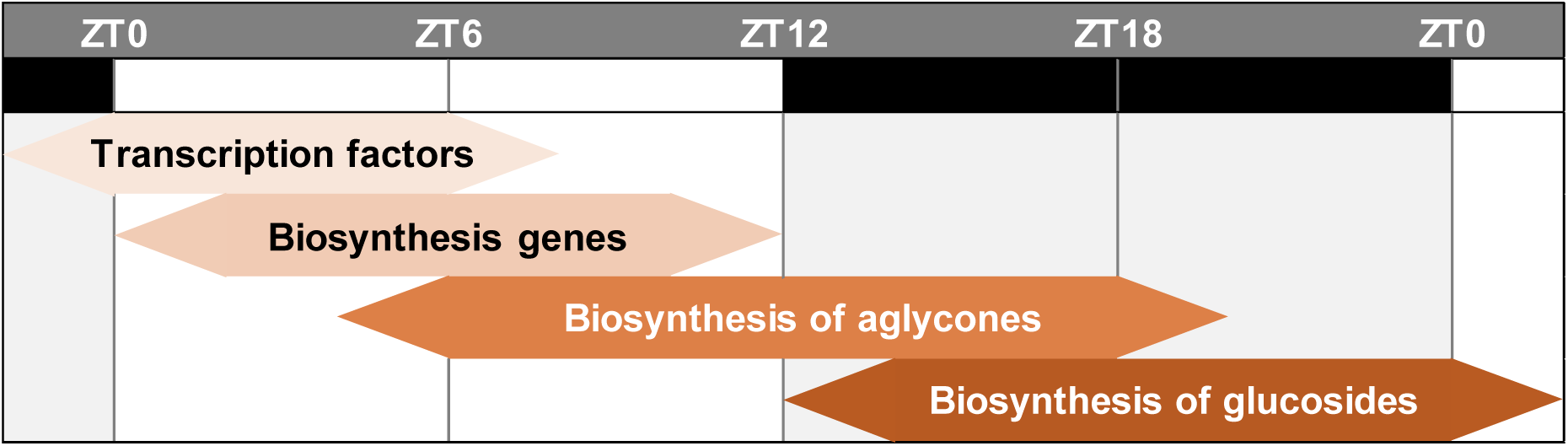
Diurnal cycle of isoflavone biosynthesis in soybean roots. ZT, hours after dawn.

## Acknowledgments

This study was supported in part by grants from JST-CREST (JPMJCR17O2 to A.S. and Y.A., JPMJCR15O2 to A.J.N.), and JSPS KAKENHI (18H02313 to A.S.); from the Research Institute for Sustainable Humanosphere (Mission 1).

We thank Ms. Keiko Kanai and Ms. Kyoko Y-Mogami for the technical assistance. We also thank DASH/FBAS, the Research Institute for Sustainable Humanosphere, Kyoto University, for supporting institutional setting.

## Author contributions

H.M., M.N. and A.S. conceived and designed the research; K.Y. and A.S. supervised the experiments; H.M., M.N. and A.S. conducted plant sampling and extraction; A.J.N. conducted RNA-seq experiments; Y.A. and S.Y. performed RNA-seq data analysis; H.M. and M.N. conducted LC-MS/MS analysis; H.M. and Y.A. constructed the correlation network and performed data analysis; H.M., Y.A., A.J.N. and A.S. wrote the article with contributions of all the authors; A.S. agrees to serve as the author responsible for contact and ensures communication.

## Conflict of Interest

The authors have no conflicts of interest directly relevant to the content of this article.

## Notes

https://www.ddbj.nig.ac.jp

### Competing Interest Statement

The authors have declared no competing interest.

