## Supplemental Figures and Tables for "Diurnal metabolic regulation of isoflavones and soyasaponins in soybean roots"

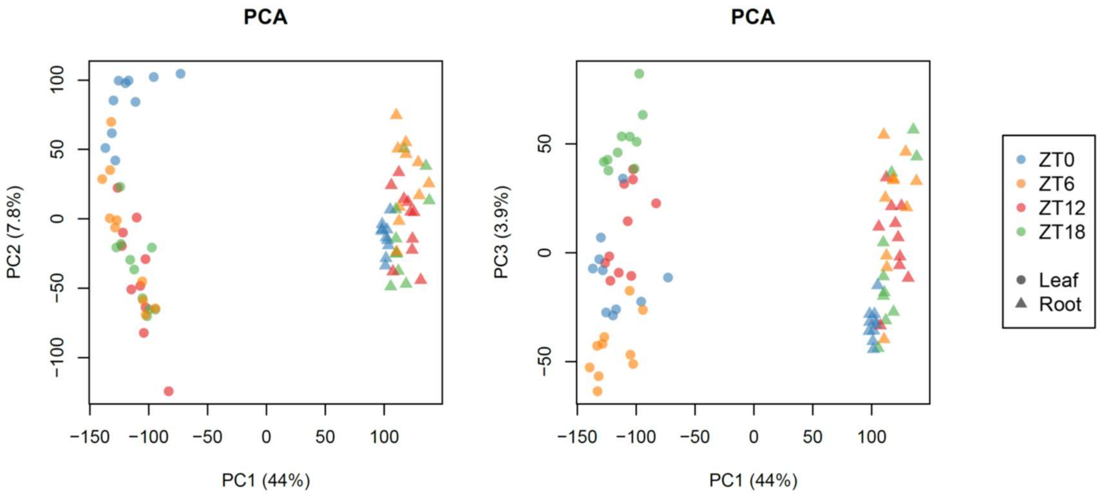


Supplemental Figure S1. Principal component analysis (PCA) for the leaf and root transcriptomes.

**A**

| | ZT12 | ZT18 | ZT0 | ZT6 | ZT12 | ZT18 | ZT0 | ZT6 |
|---|---|---|---|---|---|---|---|---|
| **Pattern 1** | | ↗ | ↘ | → ↘ | → ↘ | ↗ | ↘ | → ↘ |
| **Pattern 2** | | → ↘ | ↗ | ↘ | → ↘ | → ↘ | ↗ | ↘ |
| **Pattern 3** | | → ↘ | → ↘ | ↗ | ↘ | → ↘ | → ↘ | ↗ |
| **Pattern 4** | | ↘ | → ↘ | → ↘ | ↗ | ↘ | → ↘ | → ↘ |
| **Pattern 5** | | ↗ | ↗ → | ↘ | → ↘ | ↗ | ↗ → | ↘ |
| **Pattern 6** | | → ↘ | ↗ | ↗ → | ↘ | → ↘ | ↗ | ↗ → |
| **Pattern 7** | | ↘ | → ↘ | ↗ | ↗ → | ↘ | → ↘ | ↗ |
| **Pattern 8** | | ↗ → | ↘ | → ↘ | ↗ | ↗ → | ↘ | → ↘ |
| **Pattern 9** | | ↗ | ↗ → | ↗ → | ↘ | ↗ | ↗ → | ↗ → |
| **Pattern 10** | | ↘ | ↗ | ↗ → | ↗ → | ↘ | ↗ | ↗ → |
| **Pattern 11** | | ↗ → | ↘ | ↗ | ↗ → | ↗ → | ↘ | ↗ |
| **Pattern 12** | | ↗ → | ↗ → | ↘ | ↗ | ↗ → | ↗ → | ↘ |

Supplemental Figure S2. The 12 different diurnal patterns set for this study (A) and gene expression variations in the leaves (B) and roots (C). (B, C) The shaded areas show nighttime. The data points indicate the average of expression levels of five replicates.

Figure S2 continued

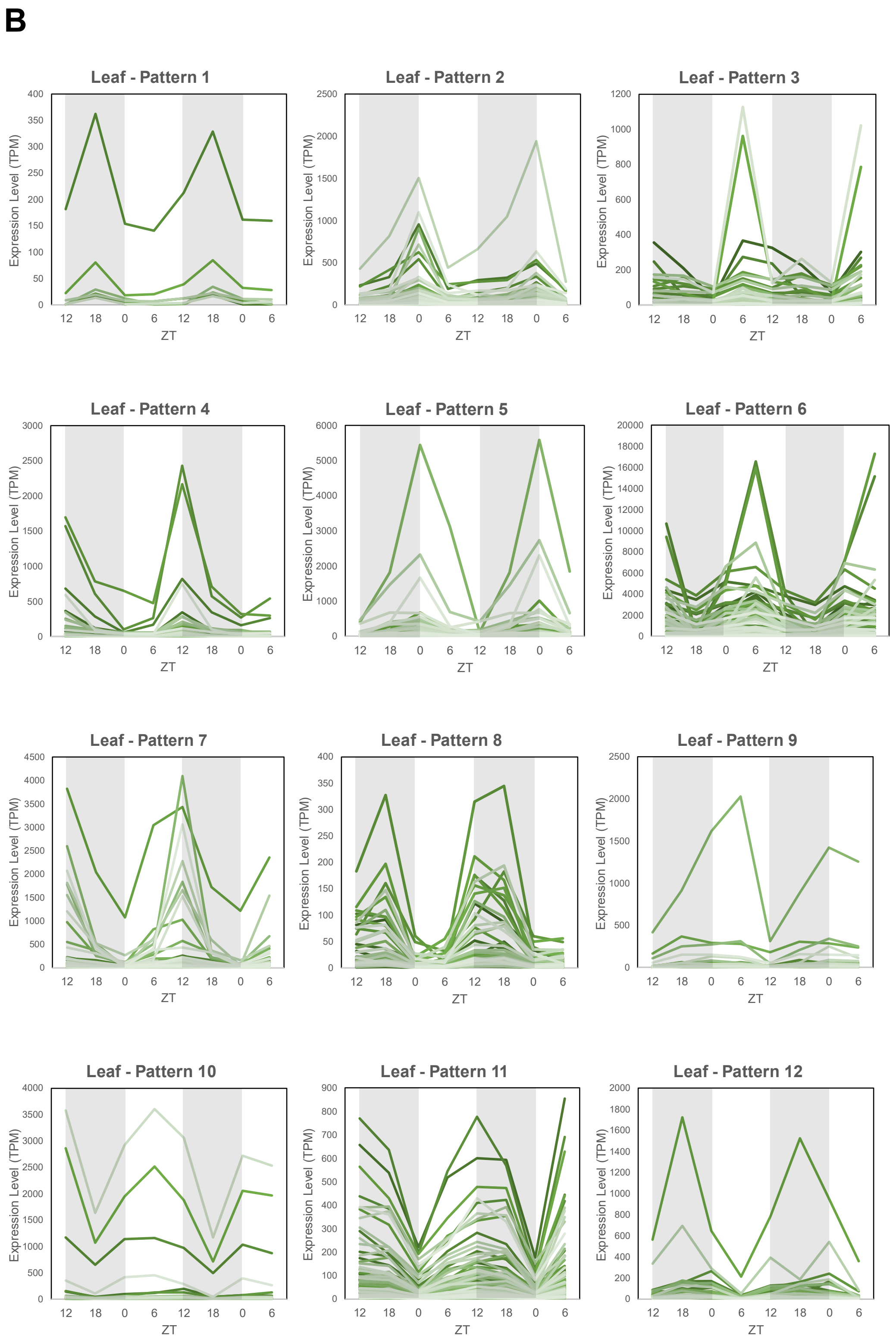

Figure S2 continued

**C**

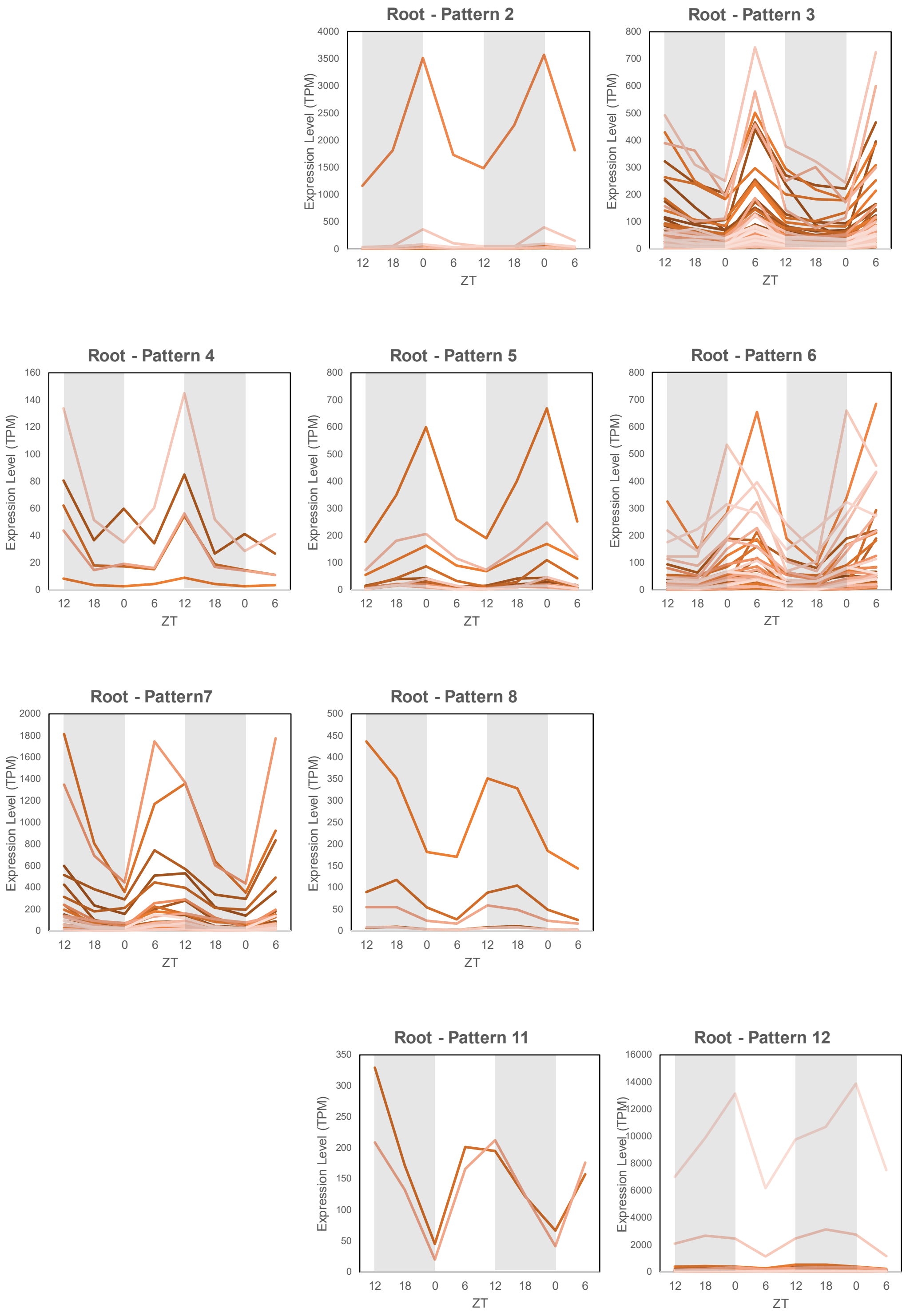

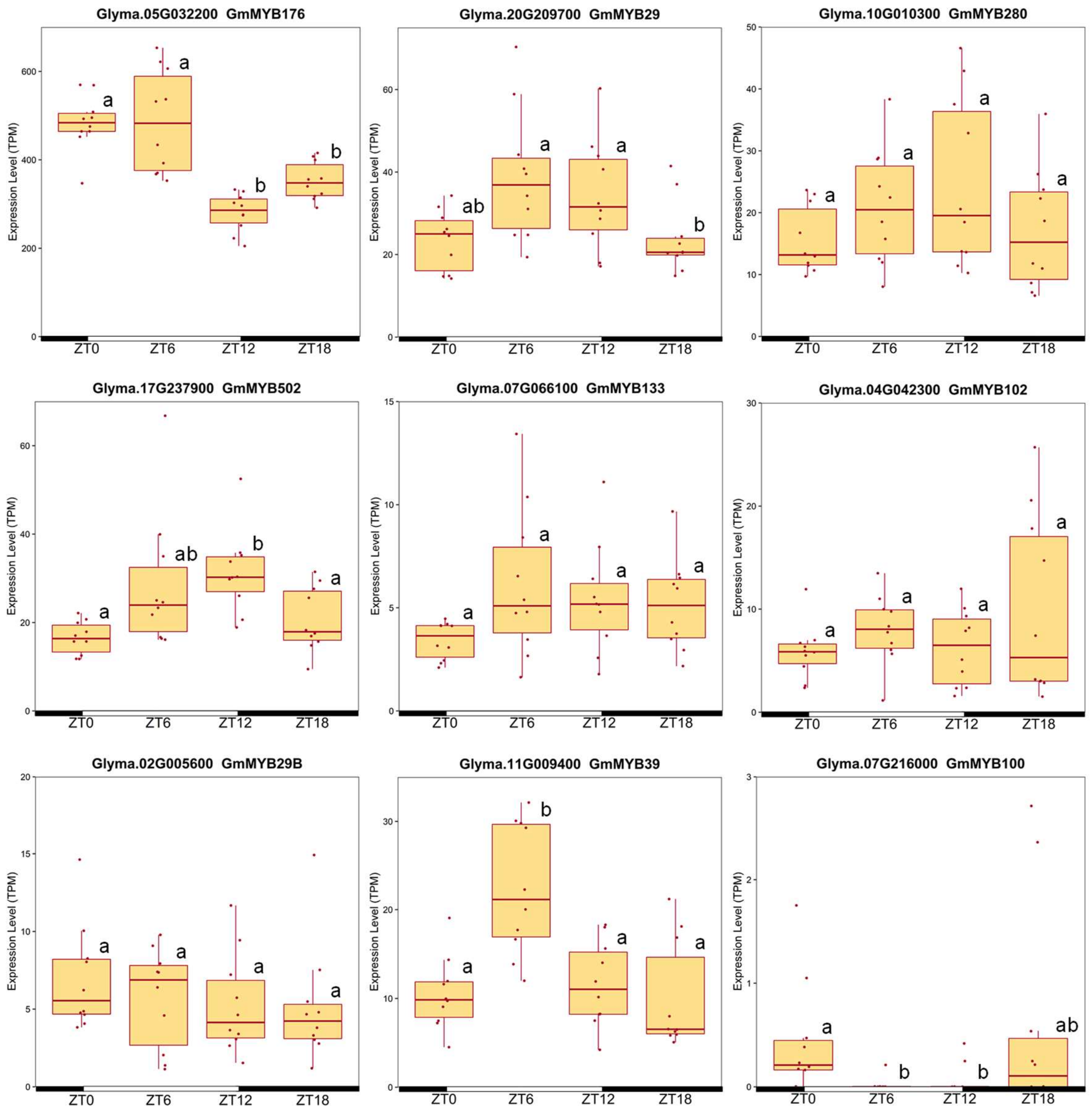


Supplemental Figure S3. The diurnal variations of MYB transcription factors regulating isoflavone biosynthesis in soybean roots. Each boxplot was constructed by ten replicates (n = 5, 2 time points). The individual dots indicate raw data. The outliers were identified with the 1.5*IQR (interquartile range) rule. Tukey's HSD test was used for statistical analysis ($p < 0.05$). ZT, hours after dawn. Abbreviations: TPM, Transcripts per million.

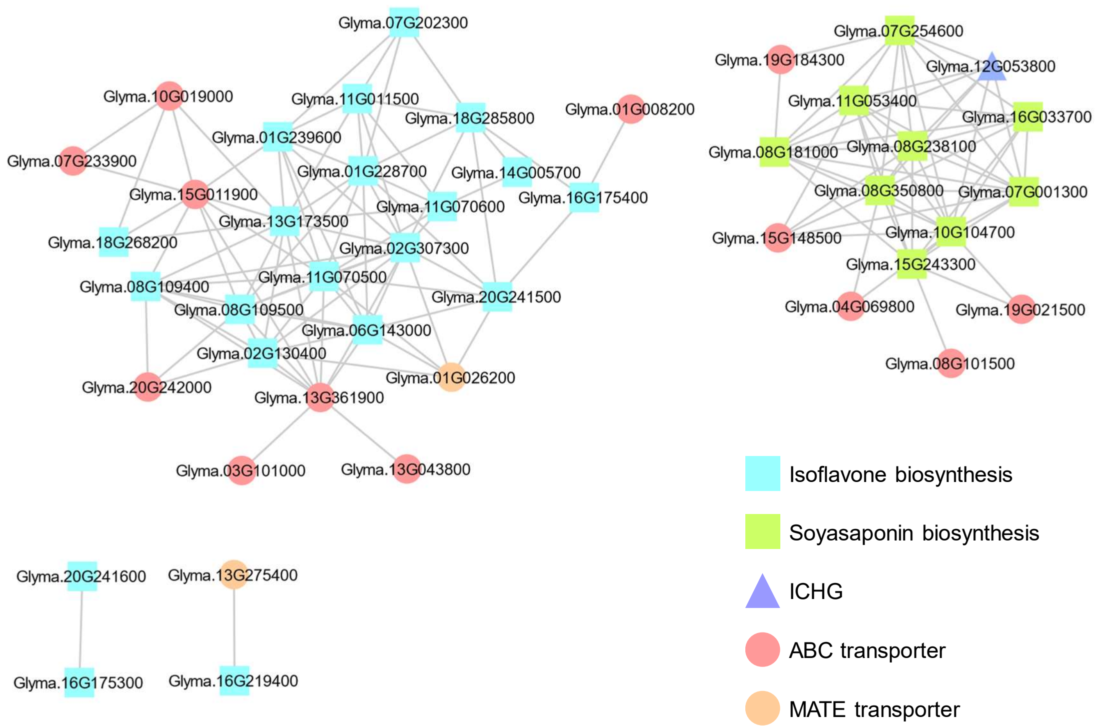


Supplemental Figure S4. Co-expression network analysis in soybean roots. Isoflavone and soyasaponin biosynthetic genes, ICHG, ABC, and MATE transporters were analyzed. The exhibited relationships have the Spearman's rank correlation coefficient > 0.7. Light blue square, isoflavone biosynthetic gene; green square, soyasaponin biosynthetic gene; blue triangle, ICHG gene; red circle, ABC transporter gene; orange circle, MATE transporter gene. Abbreviations: ICHG, Isoflavone conjugate-hydrolyzing beta-glucosidase; ABC, ATP-binding cassette transporter; MATE, Multidrug and toxic compound extrusion transporter.

Table S1 MRM conditions for LC-MS/MS analysis.

| Compound name | Parent(m/z) | Daughter(m/z) | Dwell(s) | Cone(V) | Collision(V) |
|---|---|---|---|---|---|
| daidzein | 255.3 | 137.1 | 0.025 | 30 | 30 |
| genistein | 271.3 | 153.2 | 0.025 | 30 | 30 |
| daidzin | 417.4 | 255.3 | 0.025 | 30 | 30 |
| genistin | 433.4 | 271.3 | 0.025 | 30 | 30 |
| malonyl daidzin | 503.4 | 255.3 | 0.025 | 30 | 30 |
| malonyl genistin | 519.5 | 271.3 | 0.025 | 30 | 30 |
| soyasaponin Ab | 1439.2 | 331.2 | 0.025 | 30 | 20 |
| soyasaponin Bb | 944.2 | 441.6 | 0.025 | 30 | 20 |
| soyasapogenol A | 457.7 | 95.2 | 0.025 | 30 | 30 |
| soyasapogenol B | 441.7 | 95.2 | 0.025 | 30 | 30 |

Table S2 Genes in accordance with 12 diurnal patterns.

**Leaf Pattern-01**

| GeneID | EntrezID | GeneSymbol | Description |
|---|---|---|---|
| GLYMA_01G055400 | 100790671 | LOC100790671 | uncharacterized LOC100790671 |
| GLYMA_02G241000 | 100785040 | LOC100785040 | protein REVEILLE 1 |
| GLYMA_03G194800 | NA | NA | NA |
| GLYMA_14G210600 | 778060 | MYB177 | MYB transcription factor MYB177 |
| GLYMA_18G181300 | 100819082 | PHAN3 | MYB/HD-like transcription factor |
| GLYMA_18G300800 | 100778270 | LOC100778270 | leucine-rich repeat extensin-like protein 6 |

**Leaf Pattern-02**

| GeneID | EntrezID | GeneSymbol | Description |
|---|---|---|---|
| GLYMA_01G004200 | 100800911 | LOC100800911 | probable caffeoyl-CoA O-methyltransferase At4g26220 |
| GLYMA_01G025200 | 100796652 | LOC100796652 | protein DETOXIFICATION 27 |
| GLYMA_01G033800 | NA | NA | NA |
| GLYMA_01G080500 | 102663973 | LOC102663973 | uncharacterized LOC102663973 |
| GLYMA_01G157900 | 100793320 | LOC100793320 | glutamate receptor 3.6 |
| GLYMA_01G179900 | 100786231 | LOC100786231 | BEL1-like homeodomain protein 4 |
| GLYMA_01G196600 | 100784129 | LOC100784129 | homeobox-leucine zipper protein HAT22 |
| GLYMA_01G208200 | 100527338 | LOC100527338 | aquaporin TIP2-1 |
| GLYMA_02G008000 | 100800034 | LOC100800034 | ABC transporter B family member 4 |
| GLYMA_02G017100 | 100527032 | LOC100527032 | indole-3-acetic acid-induced protein ARG2-like |
| GLYMA_02G082800 | NA | NA | NA |
| GLYMA_02G089500 | 100527161 | LOC100527161 | uncharacterized LOC100527161 |
| GLYMA_02G095400 | 100791229 | LOC100791229 | metal tolerance protein 5-like |
| GLYMA_02G099500 | 100801107 | LOC100801107 | AP2/ERF and B3 domain-containing transcription repressor TEM1 |
| GLYMA_02G123700 | 100816579 | LOC100816579 | phosphatidylinositol 4-kinase gamma 5 |
| GLYMA_02G153100 | 100810325 | LOC100810325 | glutaredoxin-C11 |
| GLYMA_02G155600 | 100816212 | LOC100816212 | universal stress protein A-like protein-like |
| GLYMA_02G240200 | 100780932 | LOC100780932 | phytoene synthase, chloroplastic-like |
| GLYMA_02G263900 | 100792450 | LOC100792450 | probable inactive receptor-like protein kinase At3g56050 |
| GLYMA_02G275900 | 100794566 | LOC100794566 | protein kinase superfamily protein |
| GLYMA_02G282100 | 100811777 | LOC100811777 | transcription factor PIF4 |
| GLYMA_03G017500 | 100810553 | LOC100810553 | RING-H2 finger protein ATL39 |
| GLYMA_03G042600 | 100803617 | LOC100803617 | F-box/LRR-repeat protein 3 |
| GLYMA_03G084300 | 100789670 | LOC100789670 | ninja-family protein 4 |
| GLYMA_03G127800 | 102661022 | LOC102661022 | sigma factor binding protein 1, chloroplastic |
| GLYMA_03G128800 | 100527837 | LOC100527837 | uncharacterized LOC100527837 |
| GLYMA_03G174500 | 100802370 | LOC100802370 | ankyrin repeat-containing protein At5g02620 |
| GLYMA_03G179100 | 102667501 | LOC102667501 | uncharacterized LOC102667501 |
| GLYMA_03G263000 | 100797420 | LOC100797420 | RING-H2 finger protein ATL3 |
| GLYMA_04G075400 | 100787213 | LOC100787213 | E3 ubiquitin-protein ligase MBR2 |
| GLYMA_04G112600 | 100791973 | LOC100791973 | actin-related protein 8 |
| GLYMA_04G123700 | 100784391 | LOC100784391 | urea-proton symporter DUR3 |
| GLYMA_04G133400 | 100785636 | LOC100785636 | protein LNK3 |
| GLYMA_04G168300 | 100803454 | LOC100803454 | cyclic dof factor 3 |
| GLYMA_04G190400 | 100811980 | LOC100811980 | leucine-rich repeat receptor-like serine/threonine/tyrosine-protein kinase SOBIR1 |
| GLYMA_04G225400 | 100795129 | LOC100795129 | uncharacterized LOC100795129 |
| GLYMA_04G229900 | NA | NA | NA |
| GLYMA_04G249400 | 100806298 | LOC100806298 | chlorophyllide a oxygenase, chloroplastic |
| GLYMA_04G254700 | NA | NA | NA |
| GLYMA_05G043800 | NA | NA | NA |
| GLYMA_05G068400 | 100788122 | LOC100788122 | BOI-related E3 ubiquitin-protein ligase 1 |
| GLYMA_05G070600 | 100808276 | LOC100808276 | protein NRT1/ PTR FAMILY 7.3 |
| GLYMA_05G096500 | 100170685 | WRKY11 | WRKY transcription factor 11 |
| GLYMA_05G112000 | 548096 | PM29 | seed maturation protein |
| GLYMA_05G180600 | 100800820 | LOC100800820 | inositol-3-phosphate synthase |
| GLYMA_05G190100 | NA | NA | NA |
| GLYMA_05G243300 | 100785302 | LOC100785302 | uncharacterized LOC100785302 |
| GLYMA_06G018700 | 100788155 | LOC100788155 | uncharacterized LOC100788155 |
| GLYMA_06G046300 | 100787252 | LOC100787252 | interferon-related developmental regulator 1 |
| GLYMA_06G095700 | 100776749 | LOC100776749 | MADS-box protein SVP-like |
| GLYMA_06G105500 | NA | NA | NA |
| GLYMA_06G106200 | 100805269 | LOC100805269 | low affinity sulfate transporter 3 |
| GLYMA_06G116000 | 100788329 | LOC100788329 | replication factor C subunit 2 |
| GLYMA_06G124000 | 100806166 | LOC100806166 | glycolipid transfer protein 3 |
| GLYMA_06G179200 | 100800308 | LOC100800308 | galactinol--sucrose galactosyltransferase |
| GLYMA_06G199600 | 102667566 | LOC102667566 | transcription factor PIF7 |
| GLYMA_06G210200 | 102669584 | LOC102669584 | uncharacterized LOC102669584 |
| GLYMA_06G255300 | 100818267 | LOC100818267 | zinc finger protein CONSTANS-LIKE 3 |
| GLYMA_06G258900 | 100794112 | LOC100794112 | G-type lectin S-receptor-like serine/threonine-protein kinase At4g27290 |
| GLYMA_06G298600 | 100817911 | LOC100817911 | uncharacterized LOC100817911 |
| GLYMA_06G319500 | 100776945 | LOC100776945 | uncharacterized LOC100776945 |
| GLYMA_07G021300 | 100808320 | LOC100808320 | uncharacterized LOC100808320 |
| GLYMA_07G139200 | 100305688 | LOC100305688 | uncharacterized LOC100305688 |
| GLYMA_07G165300 | 100808308 | LOC100808308 | uncharacterized LOC100808308 |
| GLYMA_07G184800 | 100798214 | LOC100798214 | K(+) efflux antiporter 3, chloroplastic |
| GLYMA_07G189000 | 100806717 | LOC100806717 | uncharacterized LOC100806717 |

| | | | |
|---|---|---|---|
| GLYMA_07G191000 | 100814037 | LOC100814037 | vacuolar protein sorting-associated protein 28 homolog 2 |
| GLYMA_07G194300 | NA | NA | NA |
| GLYMA_07G205400 | 100803704 | LOC100803704 | ervatamin-B |
| GLYMA_07G212800 | 100775519 | LOC100775519 | ferredoxin--nitrite reductase, chloroplastic |
| GLYMA_07G243900 | NA | NA | NA |
| GLYMA_08G047200 | 100306590 | LOC100306590 | uncharacterized LOC100306590 |
| GLYMA_08G057900 | NA | NA | NA |
| GLYMA_08G060300 | 100803004 | LOC100803004 | uncharacterized LOC100803004 |
| GLYMA_08G060500 | NA | NA | NA |
| GLYMA_08G071300 | 100776258 | LOC100776258 | protein EXORDIUM-like 2 |
| GLYMA_08G072200 | 548077 | AOX3 | alternative oxidase 3, mitochondrial |
| GLYMA_08G112500 | NA | NA | NA |
| GLYMA_08G137600 | NA | NA | NA |
| GLYMA_08G150800 | NA | NA | NA |
| GLYMA_08G152500 | 100812802 | LOC100812802 | transcription factor bHLH130 |
| GLYMA_08G182100 | NA | NA | NA |
| GLYMA_08G239000 | 100305840 | LOC100305840 | uncharacterized LOC100305840 |
| GLYMA_08G261000 | 102667538 | LOC102667538 | uncharacterized LOC102667538 |
| GLYMA_08G317200 | 100788195 | LOC100788195 | uncharacterized LOC100788195 |
| GLYMA_08G328000 | 100809940 | LOC100809940 | zinc transporter 11 |
| GLYMA_09G005700 | NA | NA | NA |
| GLYMA_09G014100 | NA | NA | NA |
| GLYMA_09G038300 | 100776839 | LOC100776839 | calmodulin-binding transcription activator 5 |
| GLYMA_09G038600 | 100812823 | LOC100812823 | beta-D-xylosidase 1 |
| GLYMA_09G061500 | 100776295 | LOC100776295 | plant intracellular Ras-group-related LRR protein 6 |
| GLYMA_09G069500 | 100795426 | LOC100795426 | uncharacterized LOC100795426 |
| GLYMA_09G071800 | 100801983 | LOC100801983 | uncharacterized LOC100801983 |
| GLYMA_09G138100 | 100779306 | LOC100779306 | oxalate--CoA ligase |
| GLYMA_09G187700 | NA | NA | NA |
| GLYMA_09G198900 | 100817263 | LOC100817263 | dirigent protein 19 |
| GLYMA_09G204100 | 100788576 | LOC100788576 | abscisic acid receptor PYR1 |
| GLYMA_09G213200 | 100811917 | LOC100811917 | uncharacterized LOC100811917 |
| GLYMA_09G250600 | 100784149 | LOC100784149 | protein METHYLENE BLUE SENSITIVITY 1 |
| GLYMA_09G250700 | 100785746 | LOC100785746 | probable tyrosine-protein phosphatase At1g05000-like |
| GLYMA_09G251200 | 100786261 | LOC100786261 | zinc finger CCCH domain-containing protein ZFN-like |
| GLYMA_09G258000 | NA | NA | NA |
| GLYMA_10G001700 | NA | NA | NA |
| GLYMA_10G003400 | 100784359 | LOC100784359 | VAN3-binding protein |
| GLYMA_10G019000 | 100793361 | LOC100793361 | ABC transporter C family member 4 |
| GLYMA_10G041200 | 100795642 | LOC100795642 | uncharacterized LOC100795642 |
| GLYMA_10G155600 | 100819956 | LOC100819956 | root phototropism protein 3 |
| GLYMA_10G183900 | 547471 | NRT1-3 | putative nitrate transporter NRT1-3 |
| GLYMA_10G185100 | 100798639 | LOC100798639 | uncharacterized LOC100798639 |
| GLYMA_10G199500 | 100785771 | LOC100785771 | uncharacterized LOC100785771 |
| GLYMA_10G231200 | 547758 | BCH3 | beta-carotene hydroxylase BCH3 |
| GLYMA_10G249500 | NA | NA | NA |
| GLYMA_10G270200 | 100775954 | LOC100775954 | neurogenic protein mastermind |
| GLYMA_10G276000 | 100775242 | LOC100775242 | uncharacterized LOC100775242 |
| GLYMA_10G289900 | 100815539 | LOC100815539 | WUSCHEL-related homeobox 13 |
| GLYMA_11G001200 | 100812510 | LOC100812510 | leucine-rich repeat protein 2 |
| GLYMA_11G045700 | 100801518 | LOC100801518 | LRR receptor-like serine/threonine-protein kinase GSO2-like |
| GLYMA_11G050500 | 100819301 | LOC100819301 | protein CONSERVED IN THE GREEN LINEAGE AND DIATOMS 27, chloroplastic |
| GLYMA_11G060100 | NA | NA | NA |
| GLYMA_11G086600 | NA | NA | NA |
| GLYMA_11G095100 | NA | NA | NA |
| GLYMA_11G139300 | 100809679 | LOC100809679 | protein kinase APK1A, chloroplastic-like |
| GLYMA_11G142900 | NA | NA | NA |
| GLYMA_11G155000 | NA | NA | NA |
| GLYMA_11G158200 | NA | NA | NA |
| GLYMA_11G163800 | 100810027 | LOC100810027 | uncharacterized LOC100810027 |
| GLYMA_11G178400 | 100796378 | LOC100796378 | non-specific phospholipase C1 |
| GLYMA_11G179300 | 100817682 | LOC100817682 | apoptosis-inducing factor homolog A |
| GLYMA_11G228900 | 100795125 | CYP90A15 | cytochrome P450 90A1 |
| GLYMA_11G239200 | NA | NA | NA |
| GLYMA_11G252200 | 100526959 | LOC100526959 | uncharacterized LOC100526959 |
| GLYMA_11G252700 | 100792859 | LOC100792859 | MACPF domain-containing protein At1g14780 |
| GLYMA_12G009000 | NA | NA | NA |
| GLYMA_12G013200 | 100799211 | LOC100799211 | putative glycerol-3-phosphate transporter 1 |

| | | | |
|---|---|---|---|
| GLYMA_12G063600 | 100815043 | LOC100815043 | putative E3 ubiquitin-protein ligase XBAT31 |
| GLYMA_12G063700 | 100815575 | LOC100815575 | U-box domain-containing protein 13 |
| GLYMA_12G133500 | 100777976 | LOC100777976 | probable serine/threonine-protein kinase abkC |
| GLYMA_12G170100 | 100810394 | LOC100810394 | uncharacterized LOC100810394 |
| GLYMA_12G172500 | NA | NA | NA |
| GLYMA_12G181500 | 100796036 | LOC100796036 | uncharacterized protein At4g22758 |
| GLYMA_12G196200 | 100305571 | LOC100305571 | uncharacterized LOC100305571 |
| GLYMA_12G212100 | 100786494 | LOC100786494 | chaperone protein dnaJ C76, chloroplastic |
| GLYMA_12G222500 | 100787229 | LOC100787229 | CBS domain-containing protein CBSX5 |
| GLYMA_12G224000 | 102660202 | LOC102660202 | uncharacterized LOC102660202 |
| GLYMA_12G234500 | NA | NA | NA |
| GLYMA_12G238400 | 100787421 | LOC100787421 | zinc finger A20 and AN1 domain-containing stress-associated protein 5 |
| GLYMA_13G033400 | 106794110 | LOC106794110 | LEAF RUST 10 DISEASE-RESISTANCE LOCUS RECEPTOR-LIKE PROTEIN KINASE-like 2.5 |
| GLYMA_13G076200 | 100793798 | LOC100793798 | putative disease resistance protein At4g11170 |
| GLYMA_13G112600 | 100805456 | LOC100805456 | auxin response factor 19 |
| GLYMA_13G155600 | 100797115 | LOC100797115 | alpha/beta hydrolase domain-containing protein 17B |
| GLYMA_13G172000 | 100802963 | IQD44 | protein IQ-DOMAIN 44 |
| GLYMA_13G191900 | 100810608 | LOC100810608 | protein LNK3 |
| GLYMA_13G233400 | 100818619 | LOC100818619 | glutamate receptor 2.7 |
| GLYMA_13G253900 | 100799584 | LOC100799584 | protein RMD5 homolog |
| GLYMA_13G267100 | 100305673 | LOC100305673 | uncharacterized LOC100305673 |
| GLYMA_13G289600 | NA | NA | NA |
| GLYMA_13G310100 | 100127384 | WRKY36 | WRKY transcription factor 36 |
| GLYMA_13G339800 | 100170750 | MATE | aluminum-activated citrate transporter |
| GLYMA_13G340500 | 100800843 | LOC100800843 | ADP,ATP carrier protein 1, mitochondrial |
| GLYMA_14G017700 | 100782666 | LOC100782666 | syntaxin-51 |
| GLYMA_14G032200 | 100811345 | LOC100811345 | transcription factor PIF4 |
| GLYMA_14G041500 | 100798037 | LOC100798037 | ETHYLENE INSENSITIVE 3-like 1 protein |
| GLYMA_14G049500 | NA | NA | NA |
| GLYMA_14G090400 | 100804406 | LOC100804406 | squalene monooxygenase |
| GLYMA_14G106500 | 100799961 | LOC100799961 | polyadenylate-binding protein-interacting protein 7 |
| GLYMA_14G154600 | 100782844 | LOC100782844 | NAD kinase 2, chloroplastic |
| GLYMA_15G016800 | 100788713 | FWL8 | protein FW2.2-like 8 |
| GLYMA_15G036200 | NA | NA | NA |
| GLYMA_15G041600 | 100786573 | LOC100786573 | monofunctional riboflavin biosynthesis protein RIBA 3, chloroplastic |
| GLYMA_15G050300 | NA | NA | CYP71D10 |
| GLYMA_15G055400 | NA | NA | NA |
| GLYMA_15G063400 | 100500019 | LOC100500019 | ribonuclease activity regulator protein RraA-like |
| GLYMA_15G141600 | 100305676 | LOC100305676 | uncharacterized LOC100305676 |
| GLYMA_15G142200 | NA | NA | NA |
| GLYMA_15G172900 | 100814577 | LOC100814577 | protein CASC3 |
| GLYMA_15G229400 | 100817056 | LOC100817056 | protein LNK3 |
| GLYMA_15G238600 | 100797696 | LOC100797696 | repetitive proline-rich cell wall protein 1 |
| GLYMA_15G271900 | 100785696 | LOC100785696 | transcription factor bHLH130 |
| GLYMA_15G272300 | 100787293 | LOC100787293 | zinc finger A20 and AN1 domain-containing stress-associated protein 5 |
| GLYMA_16G009000 | 100799109 | LOC100799109 | uncharacterized LOC100799109 |
| GLYMA_16G107300 | 100781246 | LOC100781246 | pectin acetylesterase 8 |
| GLYMA_16G109300 | 100807446 | LOC100807446 | abscisic acid 8'-hydroxylase 1 |
| GLYMA_17G006300 | NA | NA | NA |
| GLYMA_17G011600 | 100799838 | LOC100799838 | S-type anion channel SLAH4 |
| GLYMA_17G021100 | 100787515 | LOC100787515 | uncharacterized protein At4g15545 |
| GLYMA_17G064800 | 100799482 | LOC100799482 | CASP-like protein 2C1 |
| GLYMA_17G075400 | 100790159 | LOC100790159 | protein RTF2 homolog |
| GLYMA_17G090600 | 100788033 | LOC100788033 | serine/threonine-protein kinase STY13 |
| GLYMA_17G124900 | 100802333 | LOC100802333 | high-affinity nitrate transporter 3.1 |
| GLYMA_17G169700 | 100793579 | LOC100793579 | SUMO-conjugating enzyme SCE1 |
| GLYMA_17G170300 | 100792442 | LOC100792442 | floral homeotic protein APETALA 2 |
| GLYMA_17G179200 | NA | NA | NA |
| GLYMA_17G250100 | 100790867 | LOC100790867 | phosphatidylinositol transfer protein 3 |
| GLYMA_17G258500 | 100785732 | LOC100785732 | thaumatin-like protein 1 |
| GLYMA_18G020900 | 100793353 | LOC100793353 | transcription factor TGA4-like |
| GLYMA_18G022400 | NA | NA | NA |
| GLYMA_18G071900 | 100818535 | LOC100818535 | probable amino acid permease 7 |
| GLYMA_18G182000 | NA | NA | NA |
| GLYMA_18G189800 | 100500199 | LOC100500199 | uncharacterized LOC100500199 |
| GLYMA_18G192800 | 100797212 | LOC100797212 | phosphatidylinositol:ceramide inositolphosphotransferase 1 |
| GLYMA_18G200500 | 100816231 | LOC100816231 | vacuolar cation/proton exchanger 3 |
| GLYMA_18G211000 | 100799162 | LOC100799162 | cationic peroxidase 1 |

| | | | |
|---|---|---|---|
| GLYMA_18G228700 | 100793709 | LOC100793709 | SPX domain-containing membrane protein At4g22990 |
| GLYMA_18G284600 | 100815891 | LOC100815891 | glucose-6-phosphate 1-dehydrogenase 2, chloroplastic |
| GLYMA_18G286100 | 100776339 | LOC100776339 | metal tolerance protein 11 |
| GLYMA_18G289000 | NA | NA | NA |
| GLYMA_19G011700 | 100794600 | LOC100794600 | alpha-dioxygenase 1 |
| GLYMA_19G068600 | NA | NA | NA |
| GLYMA_19G129300 | 100804683 | LOC100804683 | dual specificity protein phosphatase 12 |
| GLYMA_19G141100 | NA | NA | NA |
| GLYMA_19G180000 | 100803627 | LOC100803627 | probable E3 ubiquitin-protein ligase RHG1A |
| GLYMA_19G213500 | 102668305 | LOC102668305 | chaperone protein DnaJ |
| GLYMA_19G232400 | 100814324 | LOC100814324 | dormancy-associated protein homolog 3 |
| GLYMA_19G252900 | 100785274 | LOC100785274 | NDR1/HIN1-like protein 13 |
| GLYMA_19G253000 | 100783492 | LOC100783492 | ubiquitin-conjugating enzyme E2-17 kDa |
| GLYMA_20G107600 | 100811116 | LOC100811116 | uncharacterized LOC100811116 |
| GLYMA_20G177800 | 100807197 | LOC100807197 | mitochondrial substrate carrier family protein C |
| GLYMA_20G190800 | 100804534 | LOC100804534 | cyclin-dependent protein kinase inhibitor SMR1 |
| GLYMA_20G206300 | 100800628 | LOC100800628 | protein NRT1/ PTR FAMILY 5.2 |

**Leaf Pattern-03**

| GeneID | EntrezID | GeneSymbol | Description |
|---|---|---|---|
| GLYMA_01G170200 | NA | NA | NA |
| GLYMA_02G014900 | NA | NA | NA |
| GLYMA_02G133800 | 100800733 | LOC100800733 | probable carboxylesterase 2 |
| GLYMA_02G279700 | NA | NA | NA |
| GLYMA_03G098300 | 100526914 | LOC100526914 | uncharacterized LOC100526914 |
| GLYMA_03G098800 | 100812337 | LOC100812337 | 2-alkenal reductase (NADP(+)-dependent) |
| GLYMA_03G151900 | 100800584 | LOC100800584 | basic 7S globulin |
| GLYMA_03G163300 | 100776685 | LOC100776685 | uncharacterized LOC100776685 |
| GLYMA_03G167900 | 100791074 | LOC100791074 | tyrosine decarboxylase 1 |
| GLYMA_03G180900 | 100785783 | PIP2-7 | aquaporin PIP2-7 |
| GLYMA_03G255700 | 100775980 | LOC100775980 | arabinogalactan protein 22 |
| GLYMA_03G261300 | 100779346 | LOC100779346 | two-component response regulator-like APRR5 |
| GLYMA_04G232400 | NA | NA | NA |
| GLYMA_04G247800 | 100801334 | LOC100801334 | two-component response regulator ORR9 |
| GLYMA_05G157800 | NA | NA | NA |
| GLYMA_07G089500 | 100785379 | LOC100785379 | ATP-dependent Clp protease adapter protein CLPS1, chloroplastic |
| GLYMA_08G022300 | NA | NA | NA |
| GLYMA_08G040800 | 100785888 | LOC100785888 | uncharacterized LOC100785888 |
| GLYMA_08G108900 | 100305380 | SHMT | serine hydroxylmethyltransferase |
| GLYMA_08G287600 | 100813692 | LOC100813692 | dual-specificity RNA methyltransferase RlmN |
| GLYMA_09G096300 | 100790524 | LOC100790524 | probable E3 ubiquitin-protein ligase RHY1A |
| GLYMA_09G152100 | 100306446 | LOC100306446 | uncharacterized LOC100306446 |
| GLYMA_09G187300 | 100785379 | LOC100785379 | ATP-dependent Clp protease adapter protein CLPS1, chloroplastic |
| GLYMA_09G192800 | NA | NA | NA |
| GLYMA_09G213100 | 100811384 | LOC100811384 | probable methyltransferase PMT20 |
| GLYMA_09G282900 | 100782541 | LOC100782541 | abscisic acid 8'-hydroxylase 2 |
| GLYMA_10G048100 | 100806454 | LOC100806454 | two-component response regulator-like APRR7 |
| GLYMA_10G064200 | 100779860 | LOC100779860 | probable glycosyltransferase At5g03795 |
| GLYMA_10G157900 | 100781292 | LOC100781292 | golgin IMH1 |
| GLYMA_10G221500 | 100800578 | E2 | protein GIGANTEA |
| GLYMA_10G243800 | NA | NA | NA |
| GLYMA_10G252400 | NA | NA | NA |
| GLYMA_10G269700 | 100805385 | LOC100805385 | amino-acid permease BAT1 homolog |
| GLYMA_11G175800 | 100787020 | LOC100787020 | sodium-dependent phosphate transport protein 1, chloroplastic |
| GLYMA_11G234200 | 100811436 | LOC100811436 | uncharacterized LOC100811436 |
| GLYMA_11G238800 | 547604 | MIPS | myo-inositol-3-phosphate synthase |
| GLYMA_12G028300 | 100814521 | LOC100814521 | delta(24)-sterol reductase |
| GLYMA_12G140100 | 100790931 | LOC100790931 | uncharacterized LOC100790931 |
| GLYMA_12G184800 | 100810928 | LOC100810928 | uncharacterized LOC100810928 |
| GLYMA_13G002200 | 100784781 | LOC100784781 | calcium permeable stress-gated cation channel 1 |
| GLYMA_13G029900 | 100794221 | LOC100794221 | linamarin synthase 1 |
| GLYMA_13G100700 | 100784788 | LOC100784788 | protein CHLOROPLAST IMPORT APPARATUS 2 |
| GLYMA_13G165800 | 100784254 | LOC100784254 | uncharacterized LOC100784254 |
| GLYMA_15G080200 | 100815274 | LOC100815274 | magnesium-chelatase subunit ChlI, chloroplastic |
| GLYMA_15G128000 | 100787822 | LOC100787822 | tyrosine/DOPA decarboxylase 2 |
| GLYMA_15G201100 | 100812619 | LOC100812619 | probable flavin-containing monooxygenase 1 |
| GLYMA_15G221300 | 100810117 | LOC100810117 | soyasapogenol B glucuronide galactosyltransferase-like |
| GLYMA_16G038300 | 547643 | LOC547643 | methionine synthase |
| GLYMA_16G070800 | 100812628 | LOC100812628 | uncharacterized LOC100812628 |
| GLYMA_16G134000 | 100790489 | LOC100790489 | salicylate carboxymethyltransferase |
| GLYMA_17G113600 | 100809777 | LOC100809777 | uncharacterized LOC100809777 |
| GLYMA_18G066000 | 100792104 | LOC100792104 | probable anion transporter 1, chloroplastic |
| GLYMA_18G113100 | NA | NA | NA |
| GLYMA_19G152100 | 100818913 | LOC100818913 | receptor-like cytosolic serine/threonine-protein kinase RBK2 |
| GLYMA_19G187000 | 100776894 | LOC100776894 | UDP-glycosyltransferase 73C2 |
| GLYMA_19G260200 | 100803098 | LOC100803098 | 28 kDa ribonucleoprotein, chloroplastic-like |
| GLYMA_19G260400 | 100809490 | LOC100809490 | two-component response regulator-like PRR95 |
| GLYMA_20G023100 | 100806316 | LOC100806316 | pentatricopeptide repeat-containing protein At4g18520, chloroplastic |
| GLYMA_20G094500 | 100804007 | LOC100804007 | galactinol synthase 2 |
| GLYMA_20G150600 | NA | NA | NA |
| GLYMA_20G162800 | 778166 | LOC778166 | protein REVEILLE 6 |
| GLYMA_20G164100 | 100808097 | LOC100808097 | uncharacterized LOC100808097 |
| GLYMA_20G179800 | 100816118 | LOC100816118 | uncharacterized LOC100816118 |
| GLYMA_U034500 | NA | NA | NA |

**Leaf Pattern-04**

| GeneID | EntrezID | GeneSymbol | Description |
|---|---|---|---|
| GLYMA_02G136700 | 100305614 | LOC100305614 | uncharacterized LOC100305614 |
| GLYMA_02G241200 | 100786103 | LOC100786103 | isocitrate dehydrogenase [NADP], chloroplastic |
| GLYMA_02G283500 | 100817652 | LOC100817652 | shikimate O-hydroxycinnamoyltransferase |
| GLYMA_04G092600 | NA | NA | NA |
| GLYMA_04G100400 | NA | NA | NA |
| GLYMA_04G138600 | 100306185 | LOC100306185 | uncharacterized LOC100306185 |
| GLYMA_04G238900 | 100812525 | LOC100812525 | myosin IC heavy chain |
| GLYMA_05G102100 | 100813994 | LOC100813994 | lysine-rich arabinogalactan protein 18 |
| GLYMA_06G049400 | 100818436 | LOC100818436 | probable xyloglucan glycosyltransferase 12 |
| GLYMA_06G118300 | 100795906 | LOC100795906 | PHD finger protein EHD3 |
| GLYMA_06G123700 | 100805093 | LOC100805093 | pathogen-related protein |
| GLYMA_06G185300 | 100789576 | LOC100789576 | snakin-2 |
| GLYMA_07G043100 | 100779251 | LOC100779251 | dnaJ homolog subfamily B member 9 |
| GLYMA_07G099500 | 100811519 | LOC100811519 | uncharacterized LOC100811519 |
| GLYMA_07G132000 | NA | NA | NA |
| GLYMA_07G255400 | NA | NA | NA |
| GLYMA_08G050500 | 100792588 | LOC100792588 | protein DETOXIFICATION 16 |
| GLYMA_08G343900 | NA | NA | NA |
| GLYMA_08G345000 | 100500214 | LOC100500214 | uncharacterized LOC100500214 |
| GLYMA_09G044800 | 100813899 | LOC100813899 | xyloglucan 6-xylosyltransferase 2 |
| GLYMA_09G143000 | 100783074 | LOC100783074 | uncharacterized LOC100783074 |
| GLYMA_09G181400 | 100776297 | LOC100776297 | LEAF RUST 10 DISEASE-RESISTANCE LOCUS RECEPTOR-LIKE PROTEIN KINASE-like 1.3 |
| GLYMA_09G235200 | 100780548 | PHO1-H3 | PHO1 family protein |
| GLYMA_10G230600 | 100306170 | LOC100306170 | heavy-metal-associated domain-containing protein |
| GLYMA_10G256400 | NA | NA | NA |
| GLYMA_11G029000 | 100806283 | LOC100806283 | ACT domain-containing protein ACR8 |
| GLYMA_11G039400 | 100784912 | LOC100784912 | beta-amylase 3, chloroplastic |
| GLYMA_11G096700 | NA | NA | NA |
| GLYMA_13G304400 | 100801567 | LOC100801567 | xyloglucan endotransglucosylase/hydrolase protein 9 |
| GLYMA_13G357500 | 100808818 | LOC100808818 | putative dynein light chain type 1 |
| GLYMA_14G026100 | 100788166 | LOC100788166 | uncharacterized LOC100788166 |
| GLYMA_14G083000 | 100781222 | LOC100781222 | uncharacterized LOC100781222 |
| GLYMA_15G088100 | 100789061 | LOC100789061 | MACPF domain-containing protein CAD1 |
| GLYMA_15G111400 | NA | NA | NA |
| GLYMA_16G050900 | 100801439 | COL16 | zinc finger protein CONSTANS-LIKE 15 |
| GLYMA_17G102400 | 100819400 | LOC100819400 | probable xyloglucan 6-xylosyltransferase 5 |
| GLYMA_17G242100 | 100814610 | LOC100814610 | uncharacterized LOC100814610 |
| GLYMA_18G067500 | 100811254 | LOC100811254 | uncharacterized LOC100811254 |
| GLYMA_18G151900 | 100808737 | LOC100808737 | uncharacterized LOC100808737 |

**Leaf Pattern-05**

| GeneID | EntrezID | GeneSymbol | Description |
|---|---|---|---|
| GLYMA_01G071300 | NA | NA | NA |
| GLYMA_01G087900 | 100778588 | LOC100778588 | serine/threonine-protein kinase STN7, chloroplastic |
| GLYMA_01G167700 | NA | NA | NA |
| GLYMA_01G179400 | 100816377 | LOC100816377 | cytochrome P450 71D8-like |
| GLYMA_01G208000 | 100791905 | LOC100791905 | formyltetrahydrofolate deformylase 2, mitochondrial |
| GLYMA_01G213700 | 100783063 | LOC100783063 | probable aminotransferase ACS10 |
| GLYMA_01G221400 | 100800012 | LOC100800012 | phosphoglycerate mutase-like protein AT74H |
| GLYMA_02G026300 | 778178 | MYB138 | MYB transcription factor MYB138 |
| GLYMA_02G035000 | 100803945 | LOC100803945 | carbonic anhydrase 2 |
| GLYMA_02G100000 | 100810714 | LOC100810714 | serine/threonine-protein kinase STN7, chloroplastic |
| GLYMA_02G128000 | 547981 | SAMDC | S-adenosylmethionine decarboxylase |
| GLYMA_02G192300 | 100803582 | LOC100803582 | E3 ubiquitin-protein ligase RHF2A |
| GLYMA_02G202500 | 100780041 | ALDH11A1 | aldehyde dehydrogenase 11A1 |
| GLYMA_02G278300 | 100801819 | LOC100801819 | F-box protein At2g26850 |
| GLYMA_02G283400 | 100817119 | LOC100817119 | phytoene synthase 2, chloroplastic |
| GLYMA_03G018800 | 100814309 | LOC100814309 | transcription factor TCP9 |
| GLYMA_03G063000 | 100780774 | LOC100780774 | uncharacterized LOC100780774 |
| GLYMA_03G068100 | 100795298 | RCA03 | ribulose bisphosphate carboxylase/oxygenase activase 1, chloroplastic-like |
| GLYMA_03G105200 | NA | NA | NA |
| GLYMA_03G138200 | 100808931 | LOC100808931 | guanylate kinase 1 |
| GLYMA_03G145500 | 100781689 | LOC100781689 | uncharacterized LOC100781689 |
| GLYMA_03G216900 | 100794942 | LOC100794942 | chaperone protein DnaJ |
| GLYMA_03G237900 | 100780594 | LOC100780594 | uncharacterized LOC100780594 |
| GLYMA_03G261800 | 778086 | MYB114 | MYB transcription factor MYB114 |
| GLYMA_04G058900 | 100794431 | LOC100794431 | zinc finger protein CONSTANS-LIKE 4 |
| GLYMA_04G075700 | 102668990 | LOC102668990 | F-box protein AFR |
| GLYMA_04G190500 | 100813072 | LOC100813072 | alpha/beta hydrolase domain-containing protein 17C |
| GLYMA_04G199400 | 100793556 | LOC100793556 | uncharacterized LOC100793556 |
| GLYMA_04G215900 | 100789000 | LOC100789000 | actin-7 |
| GLYMA_05G056300 | 100788844 | LOC100788844 | acid beta-fructofuranosidase |
| GLYMA_05G085400 | 100787957 | LOC100787957 | kinesin-like protein KIN-4A |
| GLYMA_05G156300 | 100784940 | LOC100784940 | protochlorophyllide-dependent translocon component 52, chloroplastic |
| GLYMA_05G169400 | 100782430 | LOC100782430 | phosphatidylinositol 4-phosphate 5-kinase 7-like |
| GLYMA_05G195900 | 100797103 | LOC100797103 | uncharacterized LOC100797103 |
| GLYMA_05G218400 | 100809536 | LOC100809536 | uncharacterized LOC100809536 |
| GLYMA_05G230700 | 100820389 | LOC100820389 | pentatricopeptide repeat-containing protein At5g21222-like |
| GLYMA_05G235800 | NA | NA | NA |
| GLYMA_05G245300 | 100778331 | LOC100778331 | probable serine/threonine-protein kinase SIS8 |
| GLYMA_06G113400 | 100786028 | LOC100786028 | TLC domain-containing protein 2 |
| GLYMA_06G113500 | 100780485 | LOC100780485 | chlorophyllide a oxygenase, chloroplastic |
| GLYMA_06G120600 | 100797840 | LOC100797840 | F-box protein SKP2A |
| GLYMA_06G139300 | 100813653 | LOC100813653 | protein RESPONSE TO LOW SULFUR 3 |
| GLYMA_06G302800 | 100788156 | LOC100788156 | protein translation factor SUI1 homolog 1 |
| GLYMA_07G048500 | 100101859 | LCL4 | LHY1/CCA1-like protein |
| GLYMA_07G091400 | 100802113 | COL21 | CONSTANS-like zinc finger protein |
| GLYMA_07G101600 | 100816332 | LOC100816332 | probable serine/threonine-protein kinase SIS8 |
| GLYMA_07G116700 | 100786209 | LOC100786209 | transcription factor bHLH18 |
| GLYMA_07G217900 | 100785146 | PIN3A | auxin efflux carrier component 3a |
| GLYMA_08G003500 | 100785172 | LOC100785172 | uncharacterized LOC100785172 |
| GLYMA_08G038200 | 100809945 | LOC100809945 | pentatricopeptide repeat-containing protein At5g21222 |
| GLYMA_08G053000 | 100782499 | LOC100782499 | uncharacterized LOC100782499 |
| GLYMA_08G167100 | 100798053 | LOC100798053 | thiosulfate/3-mercaptopyruvate sulfurtransferase 2 |
| GLYMA_08G188000 | 100811199 | LOC100811199 | uncharacterized LOC100811199 |
| GLYMA_08G226600 | 100779812 | LOC100779812 | uncharacterized LOC100779812 |
| GLYMA_08G241200 | 100783218 | LOC100783218 | UPF0301 protein Plut_0637 |
| GLYMA_08G241600 | 100810471 | LOC100810471 | cyclic nucleotide-gated ion channel 2 |
| GLYMA_08G255200 | 100301885 | COL2a | CONSTANS-like 2a |
| GLYMA_08G314700 | 100793638 | LOC100793638 | fatty acid desaturase 4, chloroplastic |
| GLYMA_08G320700 | 100795762 | LOC100795762 | ethylene-responsive transcription factor 3 |
| GLYMA_09G010600 | 100812846 | LOC100812846 | F-box protein CPR1 |
| GLYMA_09G010700 | 100790349 | LOC100790349 | tonoplast dicarboxylate transporter |
| GLYMA_09G050800 | NA | NA | AAH2 |
| GLYMA_09G063200 | 100305450 | LOC100305450 | protein kinase |
| GLYMA_09G145700 | 100787859 | LOC100787859 | F-box protein PP2-A13 |
| GLYMA_09G216300 | 100775393 | LOC100775393 | protein NUCLEAR FUSION DEFECTIVE 4 |
| GLYMA_09G237000 | 100784308 | LOC100784308 | cyclic dof factor 2 |

| | | | |
|---|---|---|---|
| GLYMA_10G048500 | 778050 | MYB118 | MYB transcription factor MYB118 |
| GLYMA_10G063100 | NA | NA | NA |
| GLYMA_10G084500 | 100802025 | LOC100802025 | E3 ubiquitin-protein ligase RHF2A |
| GLYMA_10G149200 | 100790197 | LOC100790197 | protein DEHYDRATION-INDUCED 19 homolog 3 |
| GLYMA_10G170800 | 100802536 | LOC100802536 | zinc finger CCCH domain-containing protein ZFN-like |
| GLYMA_11G008300 | 100811424 | LOC100811424 | uncharacterized LOC100811424 |
| GLYMA_11G022300 | 100785802 | LOC100785802 | phosphoglycerate mutase-like protein AT74H |
| GLYMA_11G029500 | NA | NA | NA |
| GLYMA_11G060200 | 100777753 | LOC100777753 | cytochrome P450 CYP82D47 |
| GLYMA_11G074900 | 100795117 | LOC100795117 | probable salt tolerance-like protein At1g78600-like |
| GLYMA_11G145100 | 100819644 | LOC100819644 | cytokinin riboside 5'-monophosphate phosphoribohydrolase LOG1 |
| GLYMA_11G192600 | 100782408 | LOC100782408 | zinc finger protein ZAT5 |
| GLYMA_11G221000 | 100818034 | RCA11 | ribulose bisphosphate carboxylase/oxygenase activase, chloroplastic-like |
| GLYMA_11G233800 | 100810366 | LOC100810366 | protein RETICULATA-RELATED 5, chloroplastic |
| GLYMA_12G012800 | 100790410 | LOC100790410 | peroxisomal membrane protein 11B |
| GLYMA_12G081400 | 100785286 | LOC100785286 | zinc finger protein ZAT5 |
| GLYMA_12G117700 | 100777595 | LOC100777595 | transcription activator GLK1 |
| GLYMA_12G195900 | 100804706 | LOC100804706 | cyclin-U1-1 |
| GLYMA_13G079900 | 100803677 | LOC100803677 | long chain acyl-CoA synthetase 9, chloroplastic |
| GLYMA_13G089200 | NA | NA | NA |
| GLYMA_13G119600 | 100778006 | LOC100778006 | probable GTP diphosphokinase CRSH, chloroplastic |
| GLYMA_13G136300 | 100797466 | LOC100797466 | protein REVEILLE 8 |
| GLYMA_13G142400 | 547884 | LOC547884 | heavy metal-associated isoprenylated plant protein 5 |
| GLYMA_13G147700 | 100819871 | LOC100819871 | purine-uracil permease NCS1 |
| GLYMA_13G152300 | 778058 | MYB173 | MYB transcription factor MYB173 |
| GLYMA_13G176100 | NA | NA | NA |
| GLYMA_13G210300 | 100814013 | LOC100814013 | mitogen-activated protein kinase 15 |
| GLYMA_13G211000 | 100815072 | LOC100815072 | F-box and associated interaction domain-containing protein |
| GLYMA_13G264400 | NA | NA | NA |
| GLYMA_13G310500 | 100306390 | LOC100306390 | uncharacterized LOC100306390 |
| GLYMA_13G325900 | 732581 | PIP2-6 | aquaporin PIP2-6 |
| GLYMA_13G345400 | 100816314 | GLYII-7 | putative hydroxyacylglutathione hydrolase |
| GLYMA_13G354900 | 100800844 | LOC100800844 | NADP-dependent malic enzyme |
| GLYMA_14G036300 | 100790643 | LOC100790643 | F-box protein At2g26850 |
| GLYMA_14G047200 | 100794669 | LOC100794669 | protein NRT1/ PTR FAMILY 6.4 |
| GLYMA_14G067000 | NA | NA | NA |
| GLYMA_14G086300 | 547703 | ASP1 | L-asparaginase |
| GLYMA_14G115900 | NA | NA | NA |
| GLYMA_14G127800 | 100815093 | LOC100815093 | ABC transporter G family member 39 |
| GLYMA_14G198600 | NA | NA | NA |
| GLYMA_14G198700 | 100802106 | LOC100802106 | shewanella-like protein phosphatase 1 |
| GLYMA_14G205200 | 100818985 | LOC100818985 | trans-cinnamate 4-monooxygenase |
| GLYMA_14G206100 | 100776768 | LOC100776768 | root phototropism protein 2 |
| GLYMA_14G215400 | 100803696 | LOC100803696 | uncharacterized LOC100803696 |
| GLYMA_15G096400 | NA | NA | NA |
| GLYMA_15G101600 | 100779800 | LOC100779800 | F-box/kelch-repeat protein At3g23880 |
| GLYMA_15G156900 | NA | NA | AAH1 |
| GLYMA_16G017400 | 100271888 | LCL1 | late elongated hypocotyl and circadian clock associated-1-like protein 1 |
| GLYMA_16G056600 | 100775724 | LOC100775724 | B2 protein-like |
| GLYMA_16G077900 | 102661081 | LOC102661081 | pcbC domain-containing protein |
| GLYMA_16G171900 | 100813700 | LOC100813700 | PH, RCC1 and FYVE domains-containing protein 1 |
| GLYMA_16G217700 | 778089 | LOC778089 | protein REVEILLE 7 |
| GLYMA_17G075300 | 100789100 | ALDH11A2 | aldehyde dehydrogenase 11A2 |
| GLYMA_17G211900 | 100789623 | LOC100789623 | spindle pole body component 110 |
| GLYMA_17G238200 | 100803038 | LOC100803038 | probable isoaspartyl peptidase/L-asparaginase 2 |
| GLYMA_18G005200 | 100306350 | LOC100306350 | uncharacterized LOC100306350 |
| GLYMA_18G010500 | 100798298 | LOC100798298 | small heat shock protein, chloroplastic |
| GLYMA_18G014900 | 100777924 | LOC100777924 | homeobox-leucine zipper protein ATHB-6 |
| GLYMA_18G023200 | 100790538 | LOC100790538 | protein RETICULATA-RELATED 5, chloroplastic |
| GLYMA_18G037900 | 100795471 | LOC100795471 | telomere repeat-binding protein 3 |
| GLYMA_18G044200 | 780539 | LOC780539 | protein REVEILLE 1 |
| GLYMA_18G070500 | NA | NA | NA |
| GLYMA_18G091900 | NA | NA | NA |
| GLYMA_18G116900 | NA | NA | NA |
| GLYMA_18G216000 | 100802703 | LOC100802703 | ubiquitin-conjugating enzyme E2 28 |
| GLYMA_18G238000 | 100305540 | LOC100305540 | heavy metal-associated domain-containing protein |
| GLYMA_18G260500 | 100788247 | LOC100788247 | cyclic dof factor 2 |
| GLYMA_18G263900 | 100798823 | LOC100798823 | cyclic nucleotide-gated ion channel 2 |

| | | | |
|---|---|---|---|
| GLYMA_18G266900 | 100808584 | LOC100808584 | probable endo-1,3(4)-beta-glucanase ARB_01444 |
| GLYMA_18G278100 | NA | NA | NA |
| GLYMA_19G027400 | 100818397 | LOC100818397 | uncharacterized LOC100818397 |
| GLYMA_19G035800 | 100798148 | LOC100798148 | 7-deoxyloganetin glucosyltransferase |
| GLYMA_19G260900 | 780540 | LCL3 | LHY1/CCA1-like protein |
| GLYMA_20G146100 | NA | NA | NA |

**Leaf Pattern-06**

| GeneID | EntrezID | GeneSymbol | Description |
|---|---|---|---|
| GLYMA_01G009200 | 100805334 | LOC100805334 | lachrymatory-factor synthase |
| GLYMA_01G040000 | 100500281 | GSTU1 | tau class glutathione S-transferase |
| GLYMA_01G079100 | 100803551 | LOC100803551 | protein ACTIVITY OF BC1 COMPLEX KINASE 3, chloroplastic |
| GLYMA_01G115900 | 100785180 | LOC100785180 | chlorophyll a-b binding protein CP29.2, chloroplastic |
| GLYMA_01G121700 | 100795945 | LOC100795945 | rhodanese-like domain-containing protein 10 |
| GLYMA_01G154900 | 100777875 | LOC100777875 | probable carotenoid cleavage dioxygenase 4, chloroplastic |
| GLYMA_01G166000 | 100816709 | LOC100816709 | chaperone protein dnaJ 11, chloroplastic |
| GLYMA_02G020000 | 100499775 | LOC100499775 | uncharacterized LOC100499775 |
| GLYMA_02G061100 | 100811386 | LOC100811386 | oxygen-evolving enhancer protein 1, chloroplastic-like |
| GLYMA_02G064700 | NA | NA | NA |
| GLYMA_02G080800 | NA | NA | NA |
| GLYMA_02G088900 | 100306409 | LOC100306409 | alpha-amylase inhibitor/lipid transfer/seed storage family protein |
| GLYMA_02G141700 | NA | NA | NA |
| GLYMA_02G168000 | 732619 | VTE2-2 | homogentisate phytyltransferase VTE2-2 |
| GLYMA_02G235100 | NA | NA | NA |
| GLYMA_02G305400 | 100793702 | LOC100793702 | chlorophyll a-b binding protein 151, chloroplastic-like |
| GLYMA_03G003500 | 100787389 | PHR7 | MYB-CC domain-containing transcription factor PHR7 |
| GLYMA_03G060300 | NA | NA | NA |
| GLYMA_03G114600 | 100499794 | LOC100499794 | uncharacterized LOC100499794 |
| GLYMA_03G136900 | 100804325 | LOC100804325 | cytoplasmic tRNA 2-thiolation protein 2 |
| GLYMA_03G138700 | NA | NA | NA |
| GLYMA_03G148800 | 100787372 | LOC100787372 | receptor-like cytosolic serine/threonine-protein kinase RBK2 |
| GLYMA_03G156700 | 100804670 | LOC100804670 | sulfate transporter 3.1 |
| GLYMA_03G168700 | NA | NA | NA |
| GLYMA_03G170300 | NA | NA | NA |
| GLYMA_03G219400 | 100803973 | LOC100803973 | calvin cycle protein CP12-2, chloroplastic |
| GLYMA_03G244400 | 100787907 | LOC100787907 | benzyl alcohol O-benzoyltransferase |
| GLYMA_03G256500 | 100101895 | UGT1 | glucosyltransferase |
| GLYMA_04G022900 | NA | NA | NA |
| GLYMA_04G055300 | NA | NA | NA |
| GLYMA_04G112800 | NA | NA | NA |
| GLYMA_04G141400 | 100812174 | LOC100812174 | protein LNK2 |
| GLYMA_04G167900 | 100802922 | LOC100802922 | chlorophyll a-b binding protein P4, chloroplastic |
| GLYMA_04G174400 | 100816458 | LOC100816458 | uncharacterized LOC100816458 |
| GLYMA_04G221700 | 100500559 | LOC100500559 | uncharacterized LOC100500559 |
| GLYMA_04G231800 | 100499823 | CB5-A2 | cytochrome b5 |
| GLYMA_04G241000 | 100785447 | LOC100785447 | psbP domain-containing protein 4, chloroplastic |
| GLYMA_05G003900 | 100790775 | LOC100790775 | galactinol--sucrose galactosyltransferase |
| GLYMA_05G022900 | 100807732 | LOC100807732 | photosystem I reaction center subunit III, chloroplastic |
| GLYMA_05G035600 | 100500416 | LOC100500416 | RING-H2 finger protein ATL48 |
| GLYMA_05G051300 | 100810598 | LOC100810598 | PGR5-like protein 1A, chloroplastic |
| GLYMA_05G063100 | NA | NA | NA |
| GLYMA_05G119000 | NA | NA | NA |
| GLYMA_05G127900 | 100527929 | LOC100527929 | sm-like protein LSM5 |
| GLYMA_05G128000 | 547862 | CAB3 | chlorophyll a/b-binding protein |
| GLYMA_05G166800 | NA | NA | NA |
| GLYMA_05G172300 | 100779397 | LOC100779397 | photosystem II core complex proteins psbY, chloroplastic-like |
| GLYMA_05G184700 | 100811312 | LOC100811312 | iron-sulfur assembly protein IscA, chloroplastic-like |
| GLYMA_05G192700 | 100789722 | LOC100789722 | ABC transporter G family member 5-like |
| GLYMA_05G247800 | 100817359 | LOC100817359 | purple acid phosphatase 3-like |
| GLYMA_06G141600 | 100775497 | LOC100775497 | uncharacterized LOC100775497 |
| GLYMA_06G190300 | 100783904 | LOC100783904 | uncharacterized LOC100783904 |
| GLYMA_06G194900 | 100798548 | LOC100798548 | chlorophyll a-b binding protein P4, chloroplastic |
| GLYMA_06G214700 | 100789568 | LOC100789568 | uncharacterized LOC100789568 |
| GLYMA_06G247100 | 100804565 | LOC100804565 | protochlorophyllide reductase, chloroplastic |
| GLYMA_06G263300 | 100785133 | LOC100785133 | uncharacterized LOC100785133 |
| GLYMA_06G300900 | 100782655 | LOC100782655 | zinc finger protein CONSTANS-LIKE 9 |
| GLYMA_06G312300 | NA | NA | NA |
| GLYMA_07G016900 | 100802992 | LOC100802992 | uncharacterized LOC100802992 |
| GLYMA_07G019700 | 100499832 | LOC100499832 | uncharacterized LOC100499832 |
| GLYMA_07G034400 | 100796602 | LOC100796602 | F-box/kelch-repeat protein At1g80440 |
| GLYMA_07G049400 | 100791698 | LOC100791698 | two-component response regulator-like PRR95 |
| GLYMA_07G052300 | 100798388 | LOC100798388 | cytochrome P450 78A3 |
| GLYMA_07G112000 | NA | NA | NA |
| GLYMA_07G121900 | NA | NA | NA |
| GLYMA_07G136500 | 100819003 | LOC100819003 | ultraviolet-B receptor UVR8 |

| | | | |
|---|---|---|---|
| GLYMA_07G183200 | NA | NA | NA |
| GLYMA_07G204300 | 100781940 | LOC100781940 | magnesium-chelatase subunit ChlI, chloroplastic |
| GLYMA_07G257700 | 100807247 | LOC100807247 | stem-specific protein TSJT1 |
| GLYMA_07G262000 | 100777136 | LOC100777136 | uncharacterized LOC100777136 |
| GLYMA_08G073300 | 100800895 | LOC100800895 | protein LNK1 |
| GLYMA_08G074000 | 100783223 | LOC100783223 | chlorophyll a-b binding protein CP24 10A, chloroplastic |
| GLYMA_08G082900 | 100805310 | LOC100805310 | chlorophyll a-b binding protein 3, chloroplastic-like |
| GLYMA_08G140900 | 100782151 | LOC100782151 | protein ACTIVITY OF BC1 COMPLEX KINASE 8, chloroplastic |
| GLYMA_08G164800 | 100797536 | LOC100797536 | metal tolerance protein 10 |
| GLYMA_08G179400 | 100781982 | LOC100781982 | protein LNK1 |
| GLYMA_08G180000 | NA | NA | NA |
| GLYMA_08G202200 | 102668214 | LOC102668214 | uncharacterized LOC102668214 |
| GLYMA_08G304000 | 100801237 | LOC100801237 | uncharacterized oxidoreductase At1g06690, chloroplastic |
| GLYMA_08G326000 | 100806206 | LOC100806206 | zinc finger CCCH domain-containing protein 18 |
| GLYMA_09G067700 | 100792794 | LOC100792794 | uncharacterized LOC100792794 |
| GLYMA_09G079400 | 100815660 | LOC100815660 | CBL-interacting serine/threonine-protein kinase 1 |
| GLYMA_09G087200 | 100783605 | LOC100783605 | probable inositol transporter 2 |
| GLYMA_09G154700 | NA | NA | NA |
| GLYMA_09G186200 | NA | NA | NA |
| GLYMA_09G186500 | NA | NA | NA |
| GLYMA_09G203300 | 100785203 | LOC100785203 | auxin-responsive protein IAA8 |
| GLYMA_09G207700 | 100799653 | LOC100799653 | uncharacterized protein At1g32220, chloroplastic |
| GLYMA_09G250800 | 100527240 | LOC100527240 | uncharacterized LOC100527240 |
| GLYMA_09G252100 | NA | NA | NA |
| GLYMA_10G028900 | 100810196 | LOC100810196 | sulfate transporter 3.1 |
| GLYMA_10G032200 | 100305788 | LOC100305788 | uncharacterized LOC100305788 |
| GLYMA_10G042000 | NA | NA | NA |
| GLYMA_10G080000 | 100788592 | LOC100788592 | NDR1/HIN1-like protein 1 |
| GLYMA_10G094900 | 100813025 | LOC100813025 | carboxyl-terminal-processing peptidase 3, chloroplastic |
| GLYMA_10G097800 | 100806079 | LOC100806079 | magnesium-chelatase subunit ChlH, chloroplastic |
| GLYMA_10G180000 | NA | NA | NA |
| GLYMA_10G244300 | 100500632 | LOC100500632 | 1,2-dihydroxy-3-keto-5-methylthiopentene dioxygenase 4 |
| GLYMA_10G274400 | 100815887 | LOC100815887 | 3-ketoacyl-CoA synthase 6 |
| GLYMA_11G000800 | 100816766 | LOC100816766 | putative cyclase |
| GLYMA_11G021400 | 100306378 | LOC100306378 | uncharacterized LOC100306378 |
| GLYMA_11G031500 | 100813789 | LOC100813789 | protein NRT1/ PTR FAMILY 6.3 |
| GLYMA_11G044400 | 102661842 | LOC102661842 | uncharacterized LOC102661842 |
| GLYMA_11G049300 | 100802764 | LOC100802764 | LOB domain-containing protein 38 |
| GLYMA_11G071400 | 100784545 | LOC100784545 | lipid phosphate phosphatase epsilon 2, chloroplastic |
| GLYMA_11G075300 | 100796361 | LOC100796361 | V-type proton ATPase 16 kDa proteolipid subunit |
| GLYMA_11G092900 | 100804333 | LOC100804333 | uncharacterized LOC100804333 |
| GLYMA_11G112000 | 100775983 | LOC100775983 | squalene synthase |
| GLYMA_11G127100 | NA | NA | NA |
| GLYMA_11G133800 | 100796365 | LOC100796365 | dicarboxylate transporter 1, chloroplastic |
| GLYMA_11G154700 | 100800977 | LOC100800977 | protein LNK2 |
| GLYMA_11G228000 | 100791447 | PIP1-8 | aquaporin PIP1-8 |
| GLYMA_11G232600 | 547476 | SFERH-3 | ferritin |
| GLYMA_11G241900 | NA | NA | NA |
| GLYMA_11G254900 | 100786653 | LOC100786653 | UDP-glucuronate 4-epimerase 6 |
| GLYMA_12G046000 | 100818067 | LOC100818067 | 4-hydroxy-3-methylbut-2-enyl diphosphate reductase, chloroplastic |
| GLYMA_12G051700 | NA | NA | NA |
| GLYMA_12G059100 | 100780806 | LOC100780806 | tropinone reductase |
| GLYMA_12G074000 | 100804000 | LOC100804000 | probable serine/threonine-protein kinase PBL5 |
| GLYMA_12G084900 | 100792165 | LOC100792165 | uncharacterized protein At4g22758 |
| GLYMA_12G103800 | 100789532 | LOC100789532 | B-box zinc finger protein 32 |
| GLYMA_12G151400 | NA | NA | NA |
| GLYMA_12G198400 | 100809520 | LOC100809520 | zinc finger protein CONSTANS-LIKE 3 |
| GLYMA_12G210600 | 547756 | PEPC4 | phosphoenolpyruvate carboxylase |
| GLYMA_12G218900 | 100778859 | LOC100778859 | dynein light chain 2, cytoplasmic |
| GLYMA_12G219300 | 100779387 | LOC100779387 | chlorophyll a-b binding protein 13, chloroplastic-like |
| GLYMA_12G227800 | 100800452 | LOC100800452 | sodium/calcium exchanger NCL2 |
| GLYMA_12G236700 | NA | NA | NA |
| GLYMA_13G059000 | 100811507 | LOC100811507 | uncharacterized LOC100811507 |
| GLYMA_13G090700 | 100778882 | LOC100778882 | glycine-rich cell wall structural protein 2 |
| GLYMA_13G095600 | 100810418 | LOC100810418 | 4-coumarate--CoA ligase 1 |
| GLYMA_13G127200 | 100306491 | LOC100306491 | photosystem II reaction center PSB28-like |
| GLYMA_13G129400 | 100526993 | LOC100526993 | uncharacterized LOC100526993 |
| GLYMA_13G153400 | 100792377 | LOC100792377 | uncharacterized LOC100792377 |

| | | | |
|---|---|---|---|
| GLYMA_13G183200 | NA | NA | NA |
| GLYMA_13G199300 | 100781565 | LOC100781565 | protein LNK2 |
| GLYMA_13G202200 | 100789210 | LOC100789210 | carotenoid 9,10(9',10')-cleavage dioxygenase 1 |
| GLYMA_13G207800 | NA | NA | NA |
| GLYMA_13G248000 | 100306080 | LOC100306080 | B-box type zinc finger family protein |
| GLYMA_13G253100 | 100796946 | LOC100796946 | squalene monooxygenase |
| GLYMA_13G282000 | 100790960 | LOC100790960 | chlorophyll a-b binding protein 13, chloroplastic-like |
| GLYMA_13G297400 | NA | NA | NA |
| GLYMA_13G357300 | NA | NA | NA |
| GLYMA_14G003400 | NA | NA | NA |
| GLYMA_14G008000 | 100815789 | LOC100815789 | chlorophyll a-b binding protein |
| GLYMA_14G009300 | NA | NA | NA |
| GLYMA_14G185700 | 100815985 | LOC100815985 | glutamyl-tRNA reductase 1, chloroplastic |
| GLYMA_15G008300 | NA | NA | NA |
| GLYMA_15G034400 | 100819197 | ALDH3J4 | aldehyde dehydrogenase family 3 member J4 |
| GLYMA_15G041100 | 100775900 | LOC100775900 | transcription factor MYB48-like |
| GLYMA_15G052400 | NA | NA | NA |
| GLYMA_15G053000 | 100816531 | LOC100816531 | protein LNK1 |
| GLYMA_15G061500 | NA | NA | NA |
| GLYMA_15G072500 | 100783934 | LOC100783934 | acetolactate synthase 2, chloroplastic |
| GLYMA_15G116300 | 100819190 | LOC100819190 | scarecrow-like protein 13 |
| GLYMA_15G118400 | 100776437 | LOC100776437 | pollen-specific leucine-rich repeat extensin-like protein 1 |
| GLYMA_15G161100 | NA | NA | NA |
| GLYMA_15G205900 | 100305903 | TIL' | temperature-induced lipocalin' |
| GLYMA_15G237600 | 100793985 | LOC100793985 | protein LNK2 |
| GLYMA_15G240600 | 100802304 | LOC100802304 | protein SRG1 |
| GLYMA_15G253700 | 100305752 | LOC100305752 | photosystem II subunit R-like |
| GLYMA_15G262300 | 100803531 | LOC100803531 | metal tolerance protein 10 |
| GLYMA_16G016100 | 100802138 | LOC100802138 | chlorophyll a-b binding protein 7, chloroplastic |
| GLYMA_16G020700 | NA | NA | NA |
| GLYMA_16G021200 | 100815292 | LOC100815292 | cytochrome P450 78A3-like |
| GLYMA_16G031100 | 100801965 | LOC100801965 | uncharacterized LOC100801965 |
| GLYMA_16G056900 | 100776801 | LOC100776801 | protease HtpX homolog |
| GLYMA_16G089000 | NA | NA | NA |
| GLYMA_16G143600 | 100792252 | LOC100792252 | oxygen-evolving enhancer protein 1, chloroplastic |
| GLYMA_16G145800 | 100800350 | LOC100800350 | chlorophyll a-b binding protein 6A, chloroplastic |
| GLYMA_16G162600 | NA | NA | NA |
| GLYMA_16G165200 | NA | NA | NA |
| GLYMA_16G165800 | 100800351 | LHCB1-7 | photosystem II type I chlorophyll a/b-binding protein |
| GLYMA_16G205200 | 100789786 | LOC100789786 | chlorophyll a-b binding protein CP26, chloroplastic |
| GLYMA_16G219500 | 100786750 | CHR2 | chalcone reductase CHR2 |
| GLYMA_17G012100 | NA | NA | NA |
| GLYMA_17G021800 | 100789279 | LOC100789279 | probable 1-deoxy-D-xylulose-5-phosphate synthase, chloroplastic |
| GLYMA_17G034900 | NA | NA | NA |
| GLYMA_17G042100 | 100782546 | LOC100782546 | polyubiquitin |
| GLYMA_17G069500 | NA | NA | NA |
| GLYMA_17G088400 | 100782176 | LOC100782176 | uncharacterized LOC100782176 |
| GLYMA_17G094400 | 780536 | MYB175 | MYB transcription factor MYB173 |
| GLYMA_17G096700 | 100804450 | LOC100804450 | homeobox-leucine zipper protein HAT5 |
| GLYMA_17G108300 | 100792616 | LOC100792616 | probable anion transporter 6, chloroplastic |
| GLYMA_17G174500 | NA | NA | NA |
| GLYMA_17G201600 | 100800917 | LOC100800917 | probable plastid-lipid-associated protein 8, chloroplastic |
| GLYMA_18G024600 | NA | NA | NA |
| GLYMA_18G031300 | 100778997 | LOC100778997 | proline transporter 1 |
| GLYMA_18G048800 | 100784357 | LOC100784357 | calcium uniporter protein 6, mitochondrial |
| GLYMA_18G057900 | NA | NA | NA |
| GLYMA_18G115500 | 100783464 | LOC100783464 | uncharacterized oxidoreductase At1g06690, chloroplastic |
| GLYMA_18G240500 | 100527586 | LOC100527586 | uncharacterized LOC100527586 |
| GLYMA_18G243200 | 100306555 | LOC100306555 | uncharacterized LOC100306555 |
| GLYMA_19G118700 | 100815187 | LOC100815187 | uncharacterized LOC100815187 |
| GLYMA_19G123800 | NA | NA | NA |
| GLYMA_19G139300 | 100787735 | LOC100787735 | magnesium-chelatase subunit ChlH, chloroplastic |
| GLYMA_19G153300 | 100777424 | LOC100777424 | uncharacterized LOC100777424 |
| GLYMA_19G177600 | 100795126 | LOC100795126 | UPF0308 protein At2g37240, chloroplastic-like |
| GLYMA_19G242800 | 100800250 | LOC100800250 | uncharacterized LOC100800250 |
| GLYMA_19G252700 | 100782415 | LOC100782415 | putative glucose-6-phosphate 1-epimerase |
| GLYMA_20G008300 | NA | NA | NA |
| GLYMA_20G027200 | 100804187 | LOC100804187 | nodulin-related protein 1 |

| | | | |
|---|---|---|---|
| GLYMA_20G043000 | 100777073 | LOC100777073 | hydroxymethylglutaryl-CoA lyase, mitochondrial |
| GLYMA_20G144700 | 100800992 | LOC100800992 | photosystem I reaction center subunit II, chloroplastic |
| GLYMA_20G150100 | NA | NA | NA |
| GLYMA_20G194000 | 100812913 | LOC100812913 | uncharacterized LOC100812913 |
| GLYMA_20G210100 | 100809706 | LOC100809706 | basic 7S globulin |

**Leaf Pattern-07**

| GeneID | EntrezID | GeneSymbol | Description |
|---|---|---|---|
| GLYMA_01G003400 | 100802330 | LOC100802330 | pentatricopeptide repeat-containing protein At2g15690, mitochondrial |
| GLYMA_01G017400 | NA | NA | NA |
| GLYMA_01G069500 | 100796464 | LOC100796464 | ruBisCO large subunit-binding protein subunit beta, chloroplastic |
| GLYMA_01G106700 | 100808693 | LOC100808693 | copper transporter 2 |
| GLYMA_01G162800 | 100499625 | LOC100499625 | aconitate hydratase 1 |
| GLYMA_01G182500 | 100779657 | LOC100779657 | thioredoxin-like protein HCF164, chloroplastic |
| GLYMA_01G184600 | 100794704 | LOC100794704 | uncharacterized LOC100794704 |
| GLYMA_01G188500 | 100802500 | LOC100802500 | CBS domain-containing protein CBSX1, chloroplastic |
| GLYMA_01G192100 | 100813528 | LOC100813528 | casein kinase II subunit alpha-2 |
| GLYMA_01G193600 | NA | NA | NA |
| GLYMA_01G210800 | 100526858 | LOC100526858 | uncharacterized LOC100526858 |
| GLYMA_02G074000 | 100803930 | LOC100803930 | uncharacterized LOC100803930 |
| GLYMA_02G126300 | 100785937 | LOC100785937 | ruBisCO large subunit-binding protein subunit beta, chloroplastic |
| GLYMA_02G256000 | 100778800 | LOC100778800 | probable zinc metalloprotease EGY2, chloroplastic |
| GLYMA_02G292400 | 100798628 | LOC100798628 | uncharacterized LOC100798628 |
| GLYMA_02G296500 | NA | NA | NA |
| GLYMA_02G299100 | NA | NA | NA |
| GLYMA_02G300300 | 100777912 | LOC100777912 | protein TIC 20-v, chloroplastic |
| GLYMA_03G002600 | 100810556 | LOC100810556 | protein WHAT'S THIS FACTOR 1 homolog |
| GLYMA_03G171100 | 100798307 | LOC100798307 | heat shock cognate 70 kDa protein 2 |
| GLYMA_03G195600 | 100779885 | LOC100779885 | ribonucleoside-diphosphate reductase small chain |
| GLYMA_03G232700 | 100798656 | DNAJ | chaperone protein DnaJ-like |
| GLYMA_04G007400 | 100783494 | LOC100783494 | tRNA-dihydrouridine(47) synthase [NAD(P)(+)]-like |
| GLYMA_04G023900 | 100781525 | LOC100781525 | tubulin beta chain |
| GLYMA_04G038400 | 100802935 | LOC100802935 | protein TIC 22, chloroplastic |
| GLYMA_04G178400 | 100780801 | LOC100780801 | mannose-1-phosphate guanyltransferase alpha |
| GLYMA_04G194600 | 100783850 | LOC100783850 | ribonuclease J |
| GLYMA_04G228300 | 100784383 | LOC100784383 | two-component response regulator-like APRR9 |
| GLYMA_05G025000 | 100786522 | LOC100786522 | two-component response regulator-like APRR1 |
| GLYMA_05G074800 | 100797268 | LOC100797268 | ribulose-1,5 bisphosphate carboxylase/oxygenase large subunit N-methyltransferase, chloroplastic |
| GLYMA_05G129100 | NA | NA | NA |
| GLYMA_05G152000 | NA | NA | NA |
| GLYMA_05G157300 | NA | NA | NA |
| GLYMA_05G189900 | 100781745 | LOC100781745 | uridylate kinase |
| GLYMA_05G202000 | 100789359 | LOC100789359 | probable serine/threonine-protein kinase At1g54610 |
| GLYMA_05G233500 | 100813100 | LOC100813100 | uncharacterized LOC100813100 |
| GLYMA_05G239400 | NA | NA | NA |
| GLYMA_06G024100 | NA | NA | NA |
| GLYMA_06G046000 | NA | NA | NA |
| GLYMA_06G081500 | 100784426 | LOC100784426 | GTP-binding protein At2g22870 |
| GLYMA_06G094300 | 100818441 | LOC100818441 | glucose-6-phosphate isomerase 1, chloroplastic |
| GLYMA_06G114800 | 100788687 | LOC100788687 | protein disulfide-isomerase |
| GLYMA_06G136600 | 100777119 | LOC100777119 | two-component response regulator-like PRR95 |
| GLYMA_06G163500 | 100797666 | LOC100797666 | random slug protein 5-like |
| GLYMA_06G171400 | 100779771 | LOC100779771 | ribonuclease J |
| GLYMA_06G183900 | 100811501 | ALDH2B4 | aldehyde dehydrogenase family 2 member B4, mitochondrial |
| GLYMA_06G186400 | 100817201 | LOC100817201 | mannose-1-phosphate guanyltransferase alpha-B |
| GLYMA_07G017800 | 102668733 | LOC102668733 | bZIP transcription factor 17 |
| GLYMA_07G031100 | 100794849 | LOC100794849 | phospholipase D alpha 1 |
| GLYMA_07G033100 | 100793284 | LOC100793284 | ADP,ATP carrier protein 1, chloroplastic |
| GLYMA_07G058200 | 100799267 | LOC100799267 | protein SUPPRESSOR OF PHYA-105 1 |
| GLYMA_07G077400 | 100813493 | LOC100813493 | probable methyltransferase PMT15 |
| GLYMA_07G100500 | 100814740 | LOC100814740 | beta-glucosidase 44 |
| GLYMA_07G126200 | 100777131 | LOC100777131 | glutathione hydrolase 3 |
| GLYMA_07G128600 | 100783921 | LOC100783921 | pentatricopeptide repeat-containing protein |
| GLYMA_07G211100 | 100815625 | LOC100815625 | probable carboxylesterase 2 |
| GLYMA_07G226600 | 100813859 | LOC100813859 | uncharacterized LOC100813859 |
| GLYMA_07G256200 | 547521 | LOC547521 | ribonucleotide reductase small subunit |
| GLYMA_08G046500 | 100812253 | LOC100812253 | adagio protein 3 |
| GLYMA_08G197100 | NA | NA | NA |
| GLYMA_08G205700 | 100813698 | LOC100813698 | kinesin-like protein KIN-13B |
| GLYMA_08G209100 | 100777154 | LOC100777154 | ADP,ATP carrier protein 1, chloroplastic |
| GLYMA_08G283700 | 100802675 | LOC100802675 | alpha-glucan water dikinase, chloroplastic |
| GLYMA_08G285000 | 100801953 | LOC100801953 | CRS2-associated factor 2, chloroplastic |
| GLYMA_08G316400 | 100797332 | LOC100797332 | NEP1-interacting protein-like 1 |
| GLYMA_08G334000 | 100789789 | LOC100789789 | alpha-glucan phosphorylase, H isozyme |

| | | | |
|---|---|---|---|
| GLYMA_09G005200 | 100775933 | LOC100775933 | peroxisomal membrane protein PMP22 |
| GLYMA_09G026100 | 100818878 | LOC100818878 | tubulin beta chain |
| GLYMA_09G065900 | 100801104 | LOC100801104 | metalloendoproteinase 1 |
| GLYMA_09G069800 | 547734 | LOC547734 | thiol protease aleurain-like |
| GLYMA_09G129400 | 100809627 | LOC100809627 | pentatricopeptide repeat-containing protein At4g21190 |
| GLYMA_09G186700 | 100783250 | LOC100783250 | probable 3-beta-hydroxysteroid-Delta(8),Delta(7)-isomerase |
| GLYMA_09G230500 | 100813721 | LOC100813721 | cryptochrome DASH, chloroplastic/mitochondrial |
| GLYMA_09G275300 | 100814602 | LOC100814602 | RING-H2 finger protein ATL74 |
| GLYMA_10G003200 | 100790901 | LOC100790901 | chlorophyllase-2, chloroplastic |
| GLYMA_10G012900 | 100814289 | LOC100814289 | calvin cycle protein CP12-2, chloroplastic |
| GLYMA_10G048000 | NA | NA | NA |
| GLYMA_10G058500 | NA | NA | NA |
| GLYMA_10G072700 | 100789895 | LOC100789895 | uncharacterized LOC100789895 |
| GLYMA_10G179200 | 100782566 | LOC100782566 | protein EXORDIUM-like 5 |
| GLYMA_10G224100 | 100306580 | LOC100306580 | thioredoxin Y1-like protein |
| GLYMA_10G251500 | 100788953 | LOC100788953 | thiamine thiazole synthase 2, chloroplastic |
| GLYMA_11G031200 | NA | NA | NA |
| GLYMA_11G049900 | 100809155 | LOC100809155 | ribonuclease III domain-containing protein RNC1, chloroplastic |
| GLYMA_11G053700 | 100527375 | LOC100527375 | CBS domain-containing protein CBSX1, chloroplastic |
| GLYMA_11G087500 | 100786840 | LOC100786840 | MLO-like protein 10 |
| GLYMA_11G103500 | 100791799 | LOC100791799 | delta(24)-sterol reductase |
| GLYMA_11G117600 | NA | NA | NA |
| GLYMA_11G118000 | 100796176 | LOC100796176 | uncharacterized LOC100796176 |
| GLYMA_11G149600 | NA | NA | NA |
| GLYMA_11G165600 | 100805754 | LOC100805754 | gamma-interferon-responsive lysosomal thiol protein |
| GLYMA_11G240000 | 100789861 | LOC100789861 | uncharacterized LOC100789861 |
| GLYMA_12G008400 | 102668481 | LOC102668481 | uncharacterized LOC102668481 |
| GLYMA_12G026000 | 100500273 | LOC100500273 | uncharacterized LOC100500273 |
| GLYMA_12G158900 | 100800275 | LOC100800275 | transcription factor TCP3 |
| GLYMA_12G191700 | 100780817 | LOC100780817 | cellulose synthase-like protein B5 |
| GLYMA_12G227200 | 100799393 | LOC100799393 | uncharacterized LOC100799393 |
| GLYMA_13G047300 | 100803666 | AOC4 | allene oxide cyclase 3, chloroplastic-like |
| GLYMA_13G057800 | 100800659 | LOC100800659 | alpha-1,4 glucan phosphorylase L isozyme, chloroplastic/amyloplastic |
| GLYMA_13G157500 | NA | NA | NA |
| GLYMA_14G022200 | 100500642 | LOC100500642 | uncharacterized LOC100500642 |
| GLYMA_14G060500 | 100816506 | LOC100816506 | probable zinc metalloprotease EGY2, chloroplastic |
| GLYMA_14G197900 | NA | NA | NA |
| GLYMA_15G011500 | 100775892 | LOC100775892 | ADP,ATP carrier protein 1, chloroplastic |
| GLYMA_15G132200 | NA | NA | NA |
| GLYMA_15G176900 | 100806553 | GER10 | germin-like protein |
| GLYMA_15G251900 | 548030 | LOC548030 | glutathione S-transferase GST 16 |
| GLYMA_15G255800 | NA | NA | NA |
| GLYMA_15G263600 | 100806731 | LOC100806731 | uncharacterized LOC100806731 |
| GLYMA_16G011500 | 100791192 | LOC100791192 | dnaJ homolog subfamily B member 6 |
| GLYMA_16G027200 | 100802144 | LOC100802144 | protein SUPPRESSOR OF PHYA-105 1 |
| GLYMA_16G046000 | 100791900 | LOC100791900 | ATP-dependent DNA helicase At3g02060, chloroplastic |
| GLYMA_16G049000 | NA | NA | NA |
| GLYMA_16G083300 | 100795042 | LOC100795042 | protein PEP-RELATED DEVELOPMENT ARRESTED 1, chloroplastic |
| GLYMA_16G153000 | 100775728 | LOC100775728 | uncharacterized LOC100775728 |
| GLYMA_16G177300 | 100781797 | LOC100781797 | pentatricopeptide repeat-containing protein At4g21190 |
| GLYMA_16G177800 | 100782877 | LOC100782877 | uncharacterized LOC100782877 |
| GLYMA_17G022700 | 100805161 | LOC100805161 | tubby-like F-box protein 3-like |
| GLYMA_17G061900 | 732648 | HPT1 | homogentisate phytylprenyltransferase |
| GLYMA_17G066200 | 100804624 | LOC100804624 | xylose isomerase |
| GLYMA_17G128500 | 100814619 | LOC100814619 | ribonuclease J |
| GLYMA_17G134500 | 100789105 | LOC100789105 | uncharacterized LOC100789105 |
| GLYMA_17G153900 | 100796315 | LOC100796315 | pentatricopeptide repeat-containing protein At5g46580, chloroplastic |
| GLYMA_17G259000 | 100820475 | LOC100820475 | synaptotagmin-2 |
| GLYMA_18G002000 | NA | NA | NA |
| GLYMA_18G045900 | 100791599 | LOC100791599 | protein DJ-1 homolog D |
| GLYMA_18G065700 | 100787350 | LOC100787350 | phosphomethylpyrimidine synthase, chloroplastic |
| GLYMA_18G140500 | 100784875 | LOC100784875 | uncharacterized LOC100784875 |
| GLYMA_18G188800 | 100783806 | LOC100783806 | protein RETICULATA, chloroplastic |
| GLYMA_18G298500 | 100812497 | LOC100812497 | heparan-alpha-glucosaminide N-acetyltransferase |
| GLYMA_19G076300 | 100802221 | LOC100802221 | ribulose-1,5 bisphosphate carboxylase/oxygenase large subunit N-methyltransferase, chloroplastic |
| GLYMA_19G102400 | NA | NA | NA |
| GLYMA_19G113000 | 100796371 | LOC100796371 | tubulin alpha-6 chain |
| GLYMA_19G125800 | 100793021 | LOC100793021 | 4-alpha-glucanotransferase DPE2 |

| | | | |
|---|---|---|---|
| GLYMA_19G169800 | NA | NA | NA |
| GLYMA_19G172200 | 100777767 | LOC100777767 | heat shock cognate 70 kDa protein 2 |
| GLYMA_19G190800 | 100783307 | LOC100783307 | peroxiredoxin-2E, chloroplastic |
| GLYMA_19G229700 | NA | NA | NA |
| GLYMA_20G013600 | NA | NA | NA |
| GLYMA_20G018000 | 100810397 | LOC100810397 | phosphomannomutase/phosphoglucomutase |
| GLYMA_20G142000 | 100794288 | LOC100794288 | thiamine thiazole synthase 2, chloroplastic |
| GLYMA_20G167700 | 100500683 | LOC100500683 | uncharacterized LOC100500683 |
| GLYMA_20G198100 | 100808436 | LOC100808436 | protein disulfide isomerase pTAC5, chloroplastic |

**Leaf Pattern-08**

| GeneID | EntrezID | GeneSymbol | Description |
|---|---|---|---|
| GLYMA_01G054400 | 100819569 | LOC100819569 | uncharacterized protein At5g01610 |
| GLYMA_02G002100 | NA | NA | NA |
| GLYMA_02G212700 | 100813198 | LOC100813198 | uncharacterized LOC100813198 |
| GLYMA_02G308100 | NA | NA | NA |
| GLYMA_02G308300 | 100803585 | LOC100803585 | uncharacterized LOC100803585 |
| GLYMA_03G011200 | 100783475 | LOC100783475 | AT-hook motif nuclear-localized protein 9 |
| GLYMA_03G124300 | 100816432 | LOC100816432 | stomatal closure-related actin-binding protein 3 |
| GLYMA_03G142500 | 100818371 | LOC100818371 | FT-interacting protein 1 |
| GLYMA_04G004500 | 100813607 | LOC100813607 | protein NUCLEAR FUSION DEFECTIVE 4 |
| GLYMA_04G166300 | 100799728 | LOC100799728 | two-component response regulator-like APRR1 |
| GLYMA_04G167500 | 100801871 | LOC100801871 | thioredoxin-like 1-2, chloroplastic |
| GLYMA_04G236900 | 100815931 | LOC100815931 | glutamate synthase [NADH], amyloplastic |
| GLYMA_05G182100 | NA | NA | NA |
| GLYMA_06G088600 | 100803139 | LOC100803139 | ATP-dependent 6-phosphofructokinase 3 |
| GLYMA_06G127900 | 100820570 | LOC100820570 | CCT motif-containing protein |
| GLYMA_06G157200 | 100818098 | LOC100818098 | beta-galactosidase 10 |
| GLYMA_06G196200 | NA | NA | NA |
| GLYMA_07G032900 | 100776958 | LOC100776958 | uncharacterized LOC100776958 |
| GLYMA_07G117300 | 100499981 | LOC100499981 | FCS-like zinc-finger domain-containing protein |
| GLYMA_07G132400 | 732551 | PHANa | phantastica transcription factor a |
| GLYMA_07G166400 | 100810981 | LOC100810981 | probable NAD(P)H dehydrogenase subunit CRR3, chloroplastic |
| GLYMA_07G213000 | 100806882 | LOC100806882 | uncharacterized LOC100806882 |
| GLYMA_08G047500 | 100814227 | LOC100814227 | vacuolar iron transporter 1 |
| GLYMA_08G139800 | NA | NA | NA |
| GLYMA_08G209300 | 100778751 | LOC100778751 | uncharacterized LOC100778751 |
| GLYMA_08G234200 | 100795579 | LOC100795579 | E3 ubiquitin-protein ligase RGLG4 |
| GLYMA_08G345100 | NA | NA | NA |
| GLYMA_09G131900 | 100812294 | LOC100812294 | glutaminyl-peptide cyclotransferase |
| GLYMA_09G216800 | NA | NA | NA |
| GLYMA_09G242800 | 778134 | BZIP80 | putative bZIP domain class transcription factor |
| GLYMA_10G125000 | 100781845 | LOC100781845 | probable pectate lyase 8 |
| GLYMA_10G161200 | NA | NA | NA |
| GLYMA_10G165700 | 100789308 | LOC100789308 | ankyrin repeat domain-containing protein 13C-B |
| GLYMA_10G274300 | 100782921 | COL22 | zinc finger protein CONSTANS-LIKE 16-like |
| GLYMA_11G108500 | 100808226 | LOC100808226 | WEB family protein At3g02930, chloroplastic |
| GLYMA_11G118300 | 100793194 | LOC100793194 | cyclase-associated protein 1 |
| GLYMA_11G247400 | 100808950 | LOC100808950 | mitotic spindle checkpoint protein MAD2 |
| GLYMA_12G122300 | 100786668 | LOC100786668 | uncharacterized LOC100786668 |
| GLYMA_12G135000 | NA | NA | NA |
| GLYMA_13G009300 | 100800832 | LOC100800832 | zinc finger protein CONSTANS-LIKE 9 |
| GLYMA_13G304900 | 100802093 | LOC100802093 | scarecrow-like protein 32 |
| GLYMA_14G063900 | 100781579 | LOC100781579 | uncharacterized LOC100781579 |
| GLYMA_14G190400 | 100783016 | LOC100783016 | zinc finger protein CONSTANS-LIKE 9 |
| GLYMA_15G020300 | 100804593 | LOC100804593 | auxin-responsive protein IAA26 |
| GLYMA_15G089300 | NA | NA | NA |
| GLYMA_17G091000 | NA | NA | NA |
| GLYMA_17G218000 | 100788575 | LOC100788575 | U-box domain-containing protein 7 |
| GLYMA_18G208200 | NA | NA | NA |
| GLYMA_18G270000 | NA | NA | NA |
| GLYMA_20G109400 | 100799389 | LOC100799389 | protein indeterminate-domain 5, chloroplastic |
| GLYMA_20G112600 | 100808438 | LOC100808438 | probable pectate lyase 12 |
| GLYMA_20G115600 | 100817528 | LOC100817528 | zinc finger protein CONSTANS-LIKE 16 |
| GLYMA_20G211000 | 100786500 | LOC100786500 | protein EXORDIUM-like 5 |
| GLYMA_U027700 | 100814477 | LOC100814477 | serine/threonine-protein kinase BSK6 |

**Leaf Pattern-09**

| GeneID | EntrezID | GeneSymbol | Description |
|---|---|---|---|
| GLYMA_01G072800 | 102664861 | LOC102664861 | uncharacterized LOC102664861 |
| GLYMA_02G182400 | NA | NA | NA |
| GLYMA_03G128900 | 100783830 | LOC100783830 | lycopene beta cyclase, chloroplastic |
| GLYMA_03G181600 | NA | NA | NA |
| GLYMA_03G187000 | 100802029 | LOC100802029 | UDP-glycosyltransferase 73C1 |
| GLYMA_03G214300 | 100789324 | LOC100789324 | phytochrome A-associated F-box protein |
| GLYMA_05G025400 | NA | NA | NA |
| GLYMA_06G122400 | 100800838 | LOC100800838 | psbP domain-containing protein 4, chloroplastic |
| GLYMA_06G146900 | 100797121 | LOC100797121 | ABC transporter F family member 5 |
| GLYMA_06G307000 | 100797126 | SIP1-6 | aquaporin SIP1-6 |
| GLYMA_07G020400 | 100781783 | LOC100781783 | B-box zinc finger protein 32 |
| GLYMA_07G058100 | 100814032 | LOC100814032 | mitochondrial adenine nucleotide transporter ADNT1 |
| GLYMA_07G199900 | 100817214 | LOC100817214 | polyubiquitin |
| GLYMA_08G235000 | 100797159 | LOC100797159 | uncharacterized LOC100797159 |
| GLYMA_09G116400 | 100787329 | LOC100787329 | trihelix transcription factor GT-2 |
| GLYMA_09G157600 | 100812296 | LOC100812296 | probable carboxylesterase 18 |
| GLYMA_10G222100 | NA | NA | NA |
| GLYMA_10G293500 | 100781676 | LOC100781676 | transketolase, chloroplastic |
| GLYMA_11G166500 | 106795184 | LOC106795184 | amino acid transporter AVT1I |
| GLYMA_12G037400 | NA | NA | NA |
| GLYMA_12G169600 | 100808254 | LOC100808254 | ferredoxin-A |
| GLYMA_13G051000 | 100805275 | LOC100805275 | ATP sulfurylase 2 |
| GLYMA_13G225000 | 100816306 | LOC100816306 | pyridoxal 5'-phosphate synthase subunit PDX1 |
| GLYMA_13G330400 | 100817565 | LOC100817565 | phototropin-1 |
| GLYMA_14G200900 | 100808134 | LOC100808134 | probable O-methyltransferase 3 |
| GLYMA_14G207500 | 100781041 | LOC100781041 | transmembrane protein 56-B |
| GLYMA_16G155100 | 100782690 | PIP2-11 | aquaporin PIP2-11 |
| GLYMA_17G007400 | 100812100 | LOC100812100 | CASP-like protein 2B1 |
| GLYMA_17G099800 | 100812460 | LOC100812460 | myb-related protein 306 |
| GLYMA_17G159000 | 100809780 | LOC100809780 | phytosulfokines |
| GLYMA_17G243700 | 100817797 | LOC100817797 | probable carboxylesterase 13 |
| GLYMA_18G080000 | 606507 | SAT1 | serine O-acetyltransferase 1 |
| GLYMA_18G144100 | 100787349 | LOC100787349 | inorganic phosphate transporter 2-1, chloroplastic |
| GLYMA_19G147400 | 100805040 | FAD2-2B | fatty acid desaturase-2 |

**Leaf Pattern-10**

| GeneID | EntrezID | GeneSymbol | Description |
|---|---|---|---|
| GLYMA_03G173300 | 100526897 | LOC100526897 | C2H2-type zinc finger protein |
| GLYMA_04G075800 | 100787739 | LOC100787739 | uncharacterized LOC100787739 |
| GLYMA_05G098200 | 100806145 | LOC100806145 | transcription factor MYB44 |
| GLYMA_06G012600 | 100777821 | LOC100777821 | glycine-rich RNA-binding protein 2 |
| GLYMA_06G045400 | 100776038 | SCTF-1 | zinc finger protein ZAT10 |
| GLYMA_06G055500 | 100780841 | LOC100780841 | uncharacterized LOC100780841 |
| GLYMA_06G157400 | 100779770 | NAC018 | NAC transcription factor |
| GLYMA_06G293200 | NA | NA | NA |
| GLYMA_09G189700 | NA | NA | NA |
| GLYMA_10G031800 | NA | NA | NA |
| GLYMA_10G204200 | 100796683 | LOC100796683 | transcription factor HHO3 |
| GLYMA_11G181100 | NA | NA | NA |
| GLYMA_12G093100 | 100811298 | LOC100811298 | probable CCR4-associated factor 1 homolog 11 |
| GLYMA_13G003200 | 100785849 | LOC100785849 | B2 protein |
| GLYMA_13G226700 | 106795509 | LOC106795509 | uncharacterized LOC106795509 |
| GLYMA_13G236500 | NA | NA | NA |
| GLYMA_14G016300 | 100783376 | LOC100783376 | zinc finger CCCH domain-containing protein 29 |
| GLYMA_14G041700 | 100800674 | LOC100800674 | NDR1/HIN1-like protein 13 |
| GLYMA_17G237900 | NA | NA | NA |
| GLYMA_17G251800 | 100798264 | LOC100798264 | uncharacterized LOC100798264 |
| GLYMA_19G161400 | 100306223 | LOC100306223 | uncharacterized LOC100306223 |
| GLYMA_20G066100 | 547685 | NRP-A | N-rich protein |
| GLYMA_20G212900 | 100817348 | LOC100817348 | chlorophyll a-b binding protein CP26, chloroplastic |

**Leaf Pattern-11**

| GeneID | EntrezID | GeneSymbol | Description |
|---|---|---|---|
| GLYMA_01G035700 | 100776108 | LOC100776108 | ubiquitin carboxyl-terminal hydrolase 36 |
| GLYMA_01G109300 | 100816898 | LOC100816898 | tubulin beta-2 chain-like |
| GLYMA_01G109700 | 100819388 | LOC100819388 | fructokinase-like 1, chloroplastic |
| GLYMA_02G011800 | 100798621 | LOC100798621 | ATP-dependent Clp protease proteolytic subunit 5, chloroplastic-like |
| GLYMA_02G152400 | 100796340 | LOC100796340 | protease Do-like 2, chloroplastic |
| GLYMA_02G205700 | 100791411 | LOC100791411 | translation initiation factor IF3-4, chloroplastic |
| GLYMA_02G223300 | 100786274 | LOC100786274 | uncharacterized LOC100786274 |
| GLYMA_02G266200 | 100808196 | LOC100808196 | tRNase Z TRZ2, chloroplastic |
| GLYMA_02G291100 | 100527559 | LOC100527559 | RidA-like protein |
| GLYMA_02G294900 | 100805543 | LOC100805543 | trigger factor-like protein TIG, Chloroplastic |
| GLYMA_02G299000 | 100819784 | LOC100819784 | proteasome subunit beta type-7-B |
| GLYMA_03G005900 | 100794766 | LOC100794766 | protein GrpE |
| GLYMA_03G103600 | 100306026 | LOC100306026 | uncharacterized LOC100306026 |
| GLYMA_03G176400 | NA | NA | NA |
| GLYMA_03G181200 | 100786832 | LOC100786832 | diaminopimelate decarboxylase 2, chloroplastic-like |
| GLYMA_03G226600 | 100779706 | LOC100779706 | DEAD-box ATP-dependent RNA helicase 3, chloroplastic |
| GLYMA_04G056000 | 100802756 | LOC100802756 | ylmG homolog protein 2, chloroplastic |
| GLYMA_04G072300 | 100776365 | LOC100776365 | psbP domain-containing protein 5, chloroplastic-like |
| GLYMA_04G099400 | 100809150 | LOC100809150 | serine--tRNA ligase, chloroplastic/mitochondrial |
| GLYMA_04G190100 | NA | NA | NA |
| GLYMA_05G019600 | 100805978 | LOC100805978 | translation initiation factor IF-1, chloroplastic |
| GLYMA_05G041900 | NA | NA | NA |
| GLYMA_05G045200 | 100500478 | LOC100500478 | nucleoside diphosphate kinase 2, chloroplastic |
| GLYMA_05G059600 | 100803850 | LOC100803850 | 40S ribosomal protein S30 |
| GLYMA_05G160300 | NA | NA | NA |
| GLYMA_05G236700 | 100803308 | LOC100803308 | aspartate--tRNA ligase, chloroplastic/mitochondrial |
| GLYMA_06G040300 | 100527410 | LOC100527410 | uncharacterized LOC100527410 |
| GLYMA_06G047400 | 100801015 | LOC100801015 | serine carboxypeptidase-like 11 |
| GLYMA_06G072700 | 100807222 | LOC100807222 | protein TIFY 8 |
| GLYMA_06G175400 | 100790280 | LOC100790280 | 31 kDa ribonucleoprotein, chloroplastic-like |
| GLYMA_07G028500 | 100803691 | LOC100803691 | pyrophosphate-energized vacuolar membrane proton pump |
| GLYMA_07G056600 | 100795739 | LOC100795739 | DEAD-box ATP-dependent RNA helicase 31 |
| GLYMA_07G120400 | 100500567 | LOC100500567 | uncharacterized LOC100500567 |
| GLYMA_07G149500 | 100778912 | LOC100778912 | thylakoid ADP,ATP carrier protein, chloroplastic |
| GLYMA_07G178800 | 100803872 | LOC100803872 | pentatricopeptide repeat-containing protein At2g15820, chloroplastic |
| GLYMA_07G179700 | 100788347 | LOC100788347 | probable N-acetyl-gamma-glutamyl-phosphate reductase, chloroplastic |
| GLYMA_07G257500 | 100806719 | LOC100806719 | aspartyl protease domain-containing protein |
| GLYMA_07G261400 | 100793089 | GLYI-7 | putative lactoylglutathione lyase |
| GLYMA_07G267500 | 100795559 | LOC100795559 | pentatricopeptide repeat-containing protein At4g16390, chloroplastic |
| GLYMA_08G033500 | 100802486 | LOC100802486 | uncharacterized tRNA/rRNA methyltransferase slr0955 |
| GLYMA_08G064900 | 100527095 | LOC100527095 | phosphoglycerate mutase-like protein 1 |
| GLYMA_08G084300 | 547575 | KASI | beta-ketoacyl-ACP synthetase I |
| GLYMA_08G123100 | 100818130 | LOC100818130 | T-complex protein 1 subunit alpha |
| GLYMA_08G139900 | 100779108 | LOC100779108 | nucleolin |
| GLYMA_08G172200 | NA | NA | NA |
| GLYMA_08G175900 | 100777674 | LOC100777674 | ruBisCO large subunit-binding protein subunit beta, chloroplastic |
| GLYMA_08G178200 | NA | NA | NA |
| GLYMA_08G207200 | 100818669 | LOC100818669 | protein TIC 20, chloroplastic |
| GLYMA_08G262800 | 100813344 | LOC100813344 | alpha-xylosidase 1 |
| GLYMA_09G031800 | NA | NA | NA |
| GLYMA_09G072100 | 100803033 | LOC100803033 | 20 kDa chaperonin, chloroplastic-like |
| GLYMA_09G078300 | 100812461 | LOC100812461 | uncharacterized LOC100812461 |
| GLYMA_09G091100 | 100791916 | LOC100791916 | heparanase-like protein 3 |
| GLYMA_09G094300 | NA | NA | NA |
| GLYMA_09G132300 | NA | NA | NA |
| GLYMA_09G193300 | 100804285 | LOC100804285 | uncharacterized LOC100804285 |
| GLYMA_09G275000 | NA | NA | NA |
| GLYMA_10G037400 | 100795108 | LOC100795108 | rho-N domain-containing protein 1, chloroplastic |
| GLYMA_10G140900 | 100808574 | LOC100808574 | DEAD-box ATP-dependent RNA helicase 3, chloroplastic |
| GLYMA_10G197700 | 100782567 | LOC100782567 | malate dehydrogenase [NADP], chloroplastic |
| GLYMA_10G220900 | 100820145 | LOC100820145 | uncharacterized LOC100820145 |
| GLYMA_11G001000 | 100306198 | LOC100306198 | uncharacterized LOC100306198 |
| GLYMA_11G041000 | 100808080 | LOC100808080 | uncharacterized LOC100808080 |
| GLYMA_11G091500 | 100799018 | LOC100799018 | WD-40 repeat-containing protein MSI3 |
| GLYMA_11G101400 | 100785606 | LOC100785606 | enoyl-[acyl-carrier-protein] reductase [NADH], chloroplastic |
| GLYMA_11G110600 | 100787028 | LOC100787028 | pentatricopeptide repeat-containing protein At1g30610, chloroplastic |

| | | | |
|---|---|---|---|
| GLYMA_11G195900 | 100806471 | LOC100806471 | ruBisCO large subunit-binding protein subunit alpha |
| GLYMA_11G201500 | 100778483 | LOC100778483 | uncharacterized LOC100778483 |
| GLYMA_11G225200 | 100780972 | LOC100780972 | rop guanine nucleotide exchange factor 5 |
| GLYMA_12G093300 | 100811827 | LOC100811827 | uncharacterized LOC100811827 |
| GLYMA_12G181600 | 100813077 | LOC100813077 | protein TIC 100 |
| GLYMA_12G229700 | 100807032 | LOC100807032 | uroporphyrinogen decarboxylase 1, chloroplastic |
| GLYMA_13G063400 | 100804040 | LOC100804040 | protease Do-like 8, chloroplastic |
| GLYMA_13G097800 | 100819685 | VTE2-1 | homogentisate phytyltransferase 1, chloroplastic-like |
| GLYMA_13G143600 | 100799580 | LOC100799580 | cell division cycle protein 48 homolog |
| GLYMA_13G221300 | 100798194 | LOC100798194 | protein PLASTID TRANSCRIPTIONALLY ACTIVE 12, chloroplastic |
| GLYMA_13G246900 | 100784070 | LOC100784070 | uncharacterized LOC100784070 |
| GLYMA_13G269900 | 100803496 | LOC100803496 | uroporphyrinogen decarboxylase 1, chloroplastic |
| GLYMA_13G285600 | 100306643 | LOC100306643 | uncharacterized LOC100306643 |
| GLYMA_13G287300 | 100804209 | LOC100804209 | uncharacterized LOC100804209 |
| GLYMA_13G338700 | 100794483 | LOC100794483 | protein TOC75-3, chloroplastic |
| GLYMA_14G018700 | 100812597 | LOC100812597 | trigger factor-like protein TIG, Chloroplastic |
| GLYMA_14G102000 | 100788693 | LOC100788693 | transmembrane 9 superfamily member 8 |
| GLYMA_15G180900 | 100814751 | LOC100814751 | 20 kDa chaperonin, chloroplastic |
| GLYMA_15G250500 | 100819369 | LOC100819369 | ruBisCO large subunit-binding protein subunit beta, chloroplastic |
| GLYMA_15G271200 | NA | NA | NA |
| GLYMA_15G276100 | 100796094 | LOC100796094 | uncharacterized LOC100796094 |
| GLYMA_16G010200 | 100818487 | LOC100818487 | protein FLUORESCENT IN BLUE LIGHT, chloroplastic-like |
| GLYMA_17G042600 | 100787873 | LOC100787873 | uncharacterized LOC100787873 |
| GLYMA_17G047200 | NA | NA | NA |
| GLYMA_17G080200 | NA | NA | infA |
| GLYMA_17G124500 | NA | NA | NA |
| GLYMA_17G125700 | NA | NA | NA |
| GLYMA_17G148400 | 100781644 | LOC100781644 | (S)-ureidoglycine aminohydrolase |
| GLYMA_17G228400 | 100776476 | SLD1.5 | delta8-sphingolipid desaturase |
| GLYMA_18G067200 | 100807501 | LOC100807501 | alpha-glucan phosphorylase, H isozyme |
| GLYMA_18G102100 | 100499940 | LOC100499940 | uncharacterized LOC100499940 |
| GLYMA_18G196000 | NA | NA | NA |
| GLYMA_18G207900 | 100814996 | LOC100814996 | patellin-6 |
| GLYMA_18G213900 | NA | NA | NA |
| GLYMA_19G079600 | 100811815 | LOC100811815 | uncharacterized LOC100811815 |
| GLYMA_19G127700 | 100798849 | LOC100798849 | tubulin beta-4 chain |
| GLYMA_19G226900 | 100797617 | LOC100797617 | GDSL esterase/lipase At1g09390 |
| GLYMA_20G077200 | 100807722 | LOC100807722 | probable voltage-gated potassium channel subunit beta |
| GLYMA_20G089600 | 100815040 | LOC100815040 | DEAD-box ATP-dependent RNA helicase 3, chloroplastic |
| GLYMA_20G103800 | 100786343 | LOC100786343 | pentatricopeptide repeat-containing protein At3g59040 |
| GLYMA_20G161700 | 100799045 | LOC100799045 | sec-independent protein translocase protein TATB, chloroplastic |
| GLYMA_20G204700 | 100527862 | LOC100527862 | uncharacterized LOC100527862 |
| GLYMA_20G205000 | 100796394 | LOC100796394 | CRM-domain containing factor CFM3A, chloroplastic/mitochondrial |
| GLYMA_20G249300 | NA | NA | NA |

**Leaf Pattern-12**

| GeneID | EntrezID | GeneSymbol | Description |
|---|---|---|---|
| GLYMA_01G008400 | 100794532 | LOC100794532 | probable iron/ascorbate oxidoreductase DDB_G0283291-like |
| GLYMA_02G213500 | 100811053 | LOC100811053 | AT-hook motif nuclear-localized protein 28 |
| GLYMA_02G247600 | 100800735 | LOC100800735 | receptor-like cytoplasmic kinase 176 |
| GLYMA_02G263800 | 100791927 | LOC100791927 | uncharacterized protein At5g39865 |
| GLYMA_03G132700 | 547822 | LOC547822 | glucan endo-1,3-beta-glucosidase |
| GLYMA_03G234400 | 100804496 | LOC100804496 | bifunctional epoxide hydrolase 2-like |
| GLYMA_04G057000 | 100815028 | LOC100815028 | copper transporter 5.1 |
| GLYMA_05G138700 | 100819680 | LOC100819680 | probable xyloglucan endotransglucosylase/hydrolase protein 28 |
| GLYMA_06G126600 | 100815774 | LOC100815774 | cyclic nucleotide-gated ion channel 1 |
| GLYMA_06G262700 | 106798964 | LOC106798964 | G-type lectin S-receptor-like serine/threonine-protein kinase At4g27290 |
| GLYMA_07G091800 | 100527326 | LOC100527326 | uncharacterized LOC100527326 |
| GLYMA_07G132900 | 100794335 | LOC100794335 | oxysterol-binding protein-related protein 3A |
| GLYMA_07G155500 | 100792229 | LOC100792229 | abscisic acid receptor PYL4 |
| GLYMA_07G239200 | NA | NA | NA |
| GLYMA_08G008800 | 100806925 | LOC100806925 | acyl carrier protein 1, chloroplastic |
| GLYMA_09G062000 | 100527492 | LOC100527492 | uncharacterized LOC100527492 |
| GLYMA_10G142500 | 100790707 | LOC100790707 | titin |
| GLYMA_12G001800 | 100794623 | LOC100794623 | patellin-3 |
| GLYMA_12G191200 | 100786344 | LOC100786344 | uncharacterized membrane protein YuiD |
| GLYMA_12G222400 | 547601 | LOC547601 | glutamine amidotransferases class-II superfamily protein |
| GLYMA_13G279200 | 100783727 | LOC100783727 | stem-specific protein TSJT1 |
| GLYMA_13G284900 | 100797665 | LOC100797665 | organic cation/carnitine transporter 4 |
| GLYMA_13G351200 | 100788152 | LOC100788152 | PRA1 family protein F3 |
| GLYMA_15G015600 | 100816528 | LOC100816528 | sulfite exporter TauE/SafE family protein 3 |
| GLYMA_16G212900 | 100814953 | LOC100814953 | probable metal-nicotianamine transporter YSL7 |
| GLYMA_17G071300 | 100791924 | LOC100791924 | uncharacterized LOC100791924 |
| GLYMA_17G092800 | 100793143 | LOC100793143 | snakin-2 |
| GLYMA_18G042900 | 100799172 | LOC100799172 | uncharacterized LOC100799172 |
| GLYMA_18G263600 | 100797766 | LOC100797766 | UPF0301 protein Plut_0637 |
| GLYMA_18G267100 | 100810004 | LOC100810004 | probable endo-1,3(4)-beta-glucanase ARB_01444 |
| GLYMA_18G267200 | 100810536 | LOC100810536 | ninja-family protein AFP3 |
| GLYMA_18G287800 | 100798636 | LOC100798636 | heat shock cognate 70 kDa protein 2 |
| GLYMA_18G287900 | 100783822 | LOC100783822 | heat shock cognate 70 kDa protein 2 |
| GLYMA_19G213300 | 100802569 | LOC100802569 | ethylene response sensor 1 |

**Root Pattern-02**

| GeneID | EntrezID | GeneSymbol | Description |
|---|---|---|---|
| GLYMA_05G210900 | 100785841 | LOC100785841 | probable polygalacturonase |
| GLYMA_06G018700 | 100788155 | LOC100788155 | uncharacterized LOC100788155 |
| GLYMA_07G010800 | 100801943 | LOC100801943 | calcineurin B-like protein 10 |
| GLYMA_08G014200 | NA | NA | NA |
| GLYMA_08G076700 | NA | NA | NA |
| GLYMA_08G122200 | 100802321 | LOC100802321 | sulfite exporter TauE/SafE family protein 3 |
| GLYMA_11G034000 | NA | NA | NA |
| GLYMA_11G158200 | NA | NA | NA |
| GLYMA_13G136300 | 100797466 | LOC100797466 | protein REVEILLE 8 |
| GLYMA_14G122000 | NA | NA | NA |
| GLYMA_18G020300 | NA | NA | NA |
| GLYMA_18G084000 | 106797177 | LOC106797177 | disease resistance protein PIK6-NP |

**Root Pattern-03**

| GeneID | EntrezID | GeneSymbol | Description |
|---|---|---|---|
| GLYMA_01G005000 | 100809761 | LOC100809761 | zinc finger CCCH domain-containing protein 20 |
| GLYMA_01G108200 | 100814779 | LOC100814779 | laccase-7 |
| GLYMA_01G115900 | 100785180 | LOC100785180 | chlorophyll a-b binding protein CP29.2, chloroplastic |
| GLYMA_01G134600 | 100777517 | PT01 | glycinol 2-dimethylallyl transferase |
| GLYMA_01G167700 | NA | NA | NA |
| GLYMA_01G185800 | 100791543 | HSF-01 | heat stress transcription factor 1 |
| GLYMA_01G204800 | 100802855 | LOC100802855 | expansin-like B1 |
| GLYMA_02G064700 | NA | NA | NA |
| GLYMA_02G123500 | NA | NA | NA |
| GLYMA_02G135100 | 100805171 | LOC100805171 | peroxidase 10 |
| GLYMA_02G276100 | NA | NA | NA |
| GLYMA_02G292800 | 100800738 | LOC100800738 | GDP-L-galactose phosphorylase 1 |
| GLYMA_03G060300 | NA | NA | NA |
| GLYMA_03G232700 | 100798656 | DNAJ | chaperone protein DnaJ-like |
| GLYMA_04G023500 | 100817868 | LOC100817868 | uncharacterized LOC100817868 |
| GLYMA_04G086700 | 100782076 | LOC100782076 | receptor-like protein kinase HSL1 |
| GLYMA_04G167900 | 100802922 | LOC100802922 | chlorophyll a-b binding protein P4, chloroplastic |
| GLYMA_04G195700 | 100781713 | LOC100781713 | F-box protein At1g67340 |
| GLYMA_04G228300 | 100784383 | LOC100784383 | two-component response regulator-like APRR9 |
| GLYMA_05G055700 | NA | NA | NA |
| GLYMA_06G010200 | NA | NA | NA |
| GLYMA_06G023400 | 100802281 | LOC100802281 | uncharacterized LOC100802281 |
| GLYMA_06G102300 | 100795180 | LOC100795180 | 7-deoxyloganetin glucosyltransferase |
| GLYMA_06G103200 | 100797839 | LOC100797839 | cryptochrome-1-like |
| GLYMA_06G106700 | NA | NA | NA |
| GLYMA_06G123800 | 100805630 | LOC100805630 | pathogen-related protein-like |
| GLYMA_06G136600 | 100777119 | LOC100777119 | two-component response regulator-like PRR95 |
| GLYMA_07G039600 | 100789399 | LOC100789399 | probable GTP diphosphokinase RSH2, chloroplastic |
| GLYMA_07G083000 | 100779789 | LOC100779789 | geraniol 8-hydroxylase |
| GLYMA_08G045300 | NA | NA | NA |
| GLYMA_08G173700 | 100499745 | LOC100499745 | uncharacterized LOC100499745 |
| GLYMA_08G182100 | NA | NA | NA |
| GLYMA_08G182200 | 100797704 | LOC100797704 | actin-101 |
| GLYMA_10G042000 | NA | NA | NA |
| GLYMA_10G042100 | 100499814 | LOC100499814 | uncharacterized LOC100499814 |
| GLYMA_10G070200 | NA | NA | NA |
| GLYMA_10G221500 | 100800578 | E2 | protein GIGANTEA |
| GLYMA_11G075500 | 100797229 | LOC100797229 | histone H1 |
| GLYMA_11G093100 | 100794610 | LOC100794610 | isoflavone 3'-hydroxylase |
| GLYMA_11G163300 | NA | NA | NA |
| GLYMA_12G224300 | 100791650 | LOC100791650 | uncharacterized LOC100791650 |
| GLYMA_13G035100 | 100779760 | LOC100779760 | probable calcium-binding protein CML44 |
| GLYMA_13G072800 | 100819514 | LOC100819514 | E3 ubiquitin-protein ligase RZF1 |
| GLYMA_13G319400 | 100786612 | LOC100786612 | uncharacterized LOC100786612 |
| GLYMA_15G100300 | NA | NA | NA |
| GLYMA_15G172500 | 100812974 | LOC100812974 | protein phosphatase 2C 50 |
| GLYMA_15G205900 | 100305903 | TIL' | temperature-induced lipocalin' |
| GLYMA_15G267900 | 100527328 | LOC100527328 | uncharacterized LOC100527328 |
| GLYMA_16G005200 | 100789089 | LOC100789089 | solute carrier family 25 member 44 |
| GLYMA_16G032400 | 100819572 | LOC100819572 | uncharacterized LOC100819572 |
| GLYMA_16G049000 | NA | NA | NA |
| GLYMA_16G145800 | 100800350 | LOC100800350 | chlorophyll a-b binding protein 6A, chloroplastic |
| GLYMA_16G209800 | NA | NA | NA |
| GLYMA_16G212400 | 100306134 | LOC100306134 | kunitz family trypsin and protease inhibitor |
| GLYMA_16G222200 | 100795046 | LOC100795046 | protein LURP-one-related 8-like |
| GLYMA_17G009800 | 100789627 | LOC100789627 | 1-aminocyclopropane-1-carboxylate oxidase |
| GLYMA_17G027400 | 547885 | LEA5 | desiccation protective protein LEA5 |
| GLYMA_17G134500 | 100789105 | LOC100789105 | uncharacterized LOC100789105 |
| GLYMA_17G137800 | NA | NA | NA |
| GLYMA_17G174500 | NA | NA | NA |
| GLYMA_17G202700 | 100803569 | LOC100803569 | probable E3 ubiquitin-protein ligase RZFP34 |
| GLYMA_17G250100 | 100790867 | LOC100790867 | phosphatidylinositol transfer protein 3 |
| GLYMA_17G250800 | 100815313 | LOC100815313 | probable LRR receptor-like serine/threonine-protein kinase At2g16250 |
| GLYMA_17G251800 | 100798264 | LOC100798264 | uncharacterized LOC100798264 |
| GLYMA_17G257200 | 100814612 | LOC100814612 | cytokinin riboside 5'-monophosphate phosphoribohydrolase LOG1-like |
| GLYMA_18G046300 | 112997608 | LOC112997608 | cysteine-rich receptor-like protein kinase 2 |

| | | | |
|---|---|---|---|
| GLYMA_18G054100 | 100778102 | LOC100778102 | probable serine/threonine-protein kinase WNK4 |
| GLYMA_18G290000 | 100526889 | LOC100526889 | Acyl-CoA N-acyltransferases (NAT) superfamily protein |
| GLYMA_19G011600 | 100791805 | LOC100791805 | WAT1-related protein At3g28050 |
| GLYMA_19G087200 | 106794112 | LOC106794112 | mitochondrial pyruvate carrier 1 |
| GLYMA_19G102400 | NA | NA | NA |
| GLYMA_19G161400 | 100306223 | LOC100306223 | uncharacterized LOC100306223 |
| GLYMA_19G169800 | NA | NA | NA |
| GLYMA_19G182300 | NA | NA | NA |
| GLYMA_20G157000 | 100787230 | LOC100787230 | uncharacterized LOC100787230 |
| GLYMA_U035000 | NA | NA | NA |

**Root Pattern-04**

| GeneID | EntrezID | GeneSymbol | Description |
|---|---|---|---|
| GLYMA_01G153300 | NA | NA | NA |
| GLYMA_06G036800 | 778051 | MYB124 | MYB transcription factor MYB124 |
| GLYMA_10G179200 | 100782566 | LOC100782566 | protein EXORDIUM-like 5 |
| GLYMA_13G095200 | 100782104 | LOC100782104 | probable xyloglucan endotransglucosylase/hydrolase protein 23 |
| GLYMA_16G178500 | 100783411 | LOC100783411 | heavy metal-associated isoprenylated plant protein 39 |

**Root Pattern-05**

| GeneID | EntrezID | GeneSymbol | Description |
|---|---|---|---|
| GLYMA_02G278300 | 100801819 | LOC100801819 | F-box protein At2g26850 |
| GLYMA_09G145700 | 100787859 | LOC100787859 | F-box protein PP2-A13 |
| GLYMA_09G167900 | 100101861 | LOC100101861 | protein REVEILLE 7-like |
| GLYMA_11G074900 | 100795117 | LOC100795117 | probable salt tolerance-like protein At1g78600-like |
| GLYMA_12G172500 | NA | NA | NA |
| GLYMA_13G142400 | 547884 | LOC547884 | heavy metal-associated isoprenylated plant protein 5 |
| GLYMA_13G152300 | 778058 | MYB173 | MYB transcription factor MYB173 |
| GLYMA_14G210600 | 778060 | MYB177 | MYB transcription factor MYB177 |
| GLYMA_16G013400 | NA | NA | NA |
| GLYMA_16G017400 | 100271888 | LCL1 | late elongated hypocotyl and circadian clock associated-1-like protein 1 |
| GLYMA_18G044200 | 780539 | LOC780539 | protein REVEILLE 1 |

**Root Pattern-06**

| GeneID | EntrezID | GeneSymbol | Description |
|---|---|---|---|
| GLYMA_01G105000 | 100806045 | LOC100806045 | protein NRT1/ PTR FAMILY 5.6 |
| GLYMA_03G260600 | 100791439 | LOC100791439 | probable L-gulonolactone oxidase 6 |
| GLYMA_04G022300 | 100820194 | LOC100820194 | CO(2)-response secreted protease |
| GLYMA_04G090000 | 100783854 | LOC100783854 | uncharacterized protein At5g65660 |
| GLYMA_04G141400 | 100812174 | LOC100812174 | protein LNK2 |
| GLYMA_04G250900 | NA | NA | NA |
| GLYMA_04G251000 | 100812537 | LOC100812537 | uncharacterized LOC100812537 |
| GLYMA_05G025900 | 100817354 | LOC100817354 | cyclic dof factor 3 |
| GLYMA_05G128000 | 547862 | CAB3 | chlorophyll a/b-binding protein |
| GLYMA_06G210200 | 102669584 | LOC102669584 | uncharacterized LOC102669584 |
| GLYMA_07G049400 | 100791698 | LOC100791698 | two-component response regulator-like PRR95 |
| GLYMA_07G244700 | 100817397 | LOC100817397 | putative protein phosphatase 2C |
| GLYMA_08G082900 | 100805310 | LOC100805310 | chlorophyll a-b binding protein 3, chloroplastic-like |
| GLYMA_08G179400 | 100781982 | LOC100781982 | protein LNK1 |
| GLYMA_08G321100 | 100796818 | LOC100796818 | aspartyl protease family protein At5g10770 |
| GLYMA_09G223700 | 100795072 | LOC100795072 | putative glycerol-3-phosphate transporter 1-like |
| GLYMA_09G263500 | 100778431 | LOC100778431 | uncharacterized LOC100778431 |
| GLYMA_11G154700 | 100800977 | LOC100800977 | protein LNK2 |
| GLYMA_12G051700 | NA | NA | NA |
| GLYMA_13G090600 | 100306116 | LOC100306116 | uncharacterized LOC100306116 |
| GLYMA_13G090700 | 100778882 | LOC100778882 | glycine-rich cell wall structural protein 2 |
| GLYMA_13G093800 | 100792197 | COL10 | zinc finger protein CONSTANS-LIKE 5-like |
| GLYMA_13G129400 | 100526993 | LOC100526993 | uncharacterized LOC100526993 |
| GLYMA_13G140000 | NA | NA | NA |
| GLYMA_13G199300 | 100781565 | LOC100781565 | protein LNK2 |
| GLYMA_13G303700 | 100801024 | LOC100801024 | zinc finger protein CONSTANS-LIKE 1 |
| GLYMA_13G333100 | NA | NA | NA |
| GLYMA_14G008000 | 100815789 | LOC100815789 | chlorophyll a-b binding protein |
| GLYMA_14G160100 | 100785324 | SWEET38 | sugar efflux transporter SWEET38 |
| GLYMA_16G016100 | 100802138 | LOC100802138 | chlorophyll a-b binding protein 7, chloroplastic |
| GLYMA_16G155000 | 100782155 | PIP2-10 | aquaporin PIP2-10 |
| GLYMA_16G205200 | 100789786 | LOC100789786 | chlorophyll a-b binding protein CP26, chloroplastic |
| GLYMA_17G066600 | 100805693 | COL8 | zinc finger protein CONSTANS-LIKE 5-like |
| GLYMA_17G069500 | NA | NA | NA |
| GLYMA_18G096400 | 100791423 | LOC100791423 | protein translation factor SUI1 homolog 2 |
| GLYMA_18G213300 | 100806984 | LOC100806984 | uncharacterized LOC100806984 |
| GLYMA_19G262400 | 100305853 | LOC100305853 | uncharacterized LOC100305853 |
| GLYMA_20G177300 | 100804182 | LOC100804182 | glycine-rich cell wall structural protein 2 |
| GLYMA_20G214200 | NA | NA | NA |

**Root Pattern-07**

| GeneID | EntrezID | GeneSymbol | Description |
|---|---|---|---|
| GLYMA_01G069000 | NA | NA | NA |
| GLYMA_02G125900 | 100777541 | LOC100777541 | uncharacterized LOC100777541 |
| GLYMA_03G132900 | 100795474 | LOC100795474 | glucan endo-1,3-beta-glucosidase |
| GLYMA_04G012700 | 100806292 | LOC100806292 | glycine-rich RNA-binding protein GRP1A |
| GLYMA_05G098200 | 100806145 | LOC100806145 | transcription factor MYB44 |
| GLYMA_06G012600 | 100777821 | LOC100777821 | glycine-rich RNA-binding protein 2 |
| GLYMA_06G123700 | 100805093 | LOC100805093 | pathogen-related protein |
| GLYMA_06G194600 | NA | NA | NA |
| GLYMA_07G132000 | NA | NA | NA |
| GLYMA_09G005200 | 100775933 | LOC100775933 | peroxisomal membrane protein PMP22 |
| GLYMA_09G236800 | 100791756 | LOC100791756 | probable calcium-binding protein CML45 |
| GLYMA_10G276700 | NA | NA | NA |
| GLYMA_11G073000 | 100499701 | LOC100499701 | uncharacterized LOC100499701 |
| GLYMA_11G117600 | NA | NA | NA |
| GLYMA_11G165600 | 100805754 | LOC100805754 | gamma-interferon-responsive lysosomal thiol protein |
| GLYMA_12G073900 | NA | NA | NA |
| GLYMA_12G077700 | NA | NA | NA |
| GLYMA_13G135900 | 100796933 | LOC100796933 | two-component response regulator-like APRR7 |
| GLYMA_16G215100 | NA | NA | NA |
| GLYMA_17G242100 | 100814610 | LOC100814610 | uncharacterized LOC100814610 |
| GLYMA_18G067500 | 100811254 | LOC100811254 | uncharacterized LOC100811254 |
| GLYMA_19G229700 | NA | NA | NA |

**Root Pattern-08**

| GeneID | EntrezID | GeneSymbol | Description |
|---|---|---|---|
| GLYMA_02G223700 | 100787695 | LOC100787695 | zinc finger protein CONSTANS-LIKE 9 |
| GLYMA_07G186600 | NA | NA | NA |
| GLYMA_10G222500 | 100804126 | LOC100804126 | peroxidase 12 |
| GLYMA_13G244800 | 100499725 | LOC100499725 | uncharacterized LOC100499725 |
| GLYMA_14G190400 | 100783016 | LOC100783016 | zinc finger protein CONSTANS-LIKE 9 |

**Root Pattern-11**

| GeneID | EntrezID | GeneSymbol | Description |
|---|---|---|---|
| GLYMA_07G209900 | 100812961 | LOC100812961 | peroxidase 10 |
| GLYMA_20G220800 | 100796041 | GER11 | germin-like protein |

**Root Pattern-12**

| GeneID | EntrezID | GeneSymbol | Description |
|---|---|---|---|
| GLYMA_01G015400 | 100808005 | LOC100808005 | uncharacterized LOC100808005 |
| GLYMA_01G171100 | 100788562 | LOC100788562 | peroxidase 72 |
| GLYMA_05G157300 | NA | NA | NA |
| GLYMA_05G187300 | 100819146 | LOC100819146 | cellulose synthase A catalytic subunit 6 [UDP-forming]-like |
| GLYMA_05G247300 | 100818955 | LOC100818955 | probable inactive receptor kinase At1g48480 |
| GLYMA_06G160200 | 100786707 | LOC100786707 | patellin-3 |
| GLYMA_07G035800 | 100779783 | LOC100779783 | uncharacterized LOC100779783 |
| GLYMA_08G107300 | 547802 | LOC547802 | xyloglucan endotransglucosylase/hydrolase 2 |
| GLYMA_08G146600 | NA | NA | NA |
| GLYMA_09G207000 | NA | NA | NA |
| GLYMA_11G023200 | 100812711 | PIP1-4 | aquaporin PIP1-4 |
| GLYMA_11G072000 | 102667724 | LOC102667724 | peroxidase 72 |
| GLYMA_11G256500 | NA | NA | NA |
| GLYMA_12G195600 | 100803114 | LOC100803114 | peroxidase 3 |
| GLYMA_12G222500 | 100787229 | LOC100787229 | CBS domain-containing protein CBSX5 |
| GLYMA_13G077800 | 100782997 | LOC100782997 | WAT1-related protein At4g08290 |
| GLYMA_13G307000 | 547872 | LOC547872 | peroxidase |
| GLYMA_14G191700 | NA | NA | NA |
| GLYMA_15G024400 | 100819194 | LOC100819194 | NDR1/HIN1-like protein 13 |
| GLYMA_15G249900 | 100818291 | LOC100818291 | polyadenylate-binding protein-interacting protein 9 |
| GLYMA_17G138300 | 100798971 | LOC100798971 | monocopper oxidase-like protein SKU5 |
| GLYMA_18G190000 | 100305635 | LOC100305635 | uncharacterized LOC100305635 |
| GLYMA_19G047800 | NA | NA | NA |
| GLYMA_20G153200 | 100777262 | LOC100777262 | probable carboxylesterase 120 |
| GLYMA_20G247800 | NA | NA | NA |

Table S3 Co-expressed biosynthetic and transporter genes in soybean roots (Spearman correlation coefficient > 0.7).

| Cluster | GeneID | Category | Symbol | Description |
|---|---|---|---|---|
| **Isoflavone** | Glyma.01G008200 | ABC Transporter | ABCB | ABC transporter B family member 8 |
| | Glyma.03G101000 | ABC Transporter | ABCC | ABC transporter C family member 15 |
| | Glyma.07G233900 | ABC Transporter | ABCG | Pleiotropic drug resistance 3-like isoform X2 |
| | Glyma.10G019000 | ABC Transporter | ABCC | ABC transporter C family member 4-like |
| | Glyma.13G043800 | ABC Transporter | ABCG | ABC transporter G family member 11-like |
| | Glyma.13G361900 | ABC Transporter | ABCG | Pleiotropic drug resistance 1 |
| | Glyma.15G011900 | ABC Transporter | ABCG | Pleiotropic drug resistance 1 |
| | Glyma.20G242000 | ABC Transporter | ABCG | ABC transporter G family member 6-like |
| | Glyma.01G026200 | MATE Transporter | GmMATE3 | NA |
| | Glyma.01G228700 | Isoflavone Biosynthesis | GmCHS7 | chalcone synthase 7 |
| | Glyma.01G239600 | Isoflavone Biosynthesis | HIDH | 2-hydroxyisoflavanone dehydratase |
| | Glyma.02G130400 | Isoflavone Biosynthesis | GmCHS10 | chalcone synthase 10 |
| | Glyma.02G307300 | Isoflavone Biosynthesis | GmCHR6 | chalcone reductase CHR6 |
| | Glyma.06G143000 | Isoflavone Biosynthesis | GmCHI4A, CHI4A | chalcone isomerase 4A |
| | Glyma.07G202300 | Isoflavone Biosynthesis | IFS1 | isoflavone synthase 1 |
| | Glyma.08G109400 | Isoflavone Biosynthesis | GmCHS1 | chalcone synthase 1 |
| | Glyma.08G109500 | Isoflavone Biosynthesis | CHS9 | chalcone synthase 9 |
| | Glyma.11G011500 | Isoflavone Biosynthesis | CHS8 | chalcone synthase 8 |
| | Glyma.11G070500 | Isoflavone Biosynthesis | IFR3 | isoflavone reductase |
| | Glyma.11G070600 | Isoflavone Biosynthesis | IFR4 | NmrA-like family domain-containing protein |
| | Glyma.13G173500 | Isoflavone Biosynthesis | IFS2 | 2-hydroxyisoflavanone synthase |
| | Glyma.14G005700 | Isoflavone Biosynthesis | GmCHR1 | chalcone reductase CHR1 |
| | Glyma.16G175400 | Isoflavone Biosynthesis | UGT4, | isoflavone 7-O-glucosyltransferase UGT4 |
| | Glyma.18G268200 | Isoflavone Biosynthesis | IMAT1 | isoflavone malonyltransferase IMaT1 |
| | Glyma.18G285800 | Isoflavone Biosynthesis | GmCHR5 | chalcone reductase CHR5 |
| | Glyma.20G241500 | Isoflavone Biosynthesis | CHI1A, CHI1 | chalcone--flavonone isomerase 1A |
| **soyasaponin** | Glyma.04G069800 | ABC Transporter | ABCG | Pleiotropic drug resistance 1 |
| | Glyma.08G101500 | ABC Transporter | ABCC | ABC transporter C family member 15 |
| | Glyma.15G148500 | ABC Transporter | ABCC | ABC transporter C family member 14-like |
| | Glyma.19G021500 | ABC Transporter | ABCB | ABC transporter B family member 19 |
| | Glyma.19G184300 | ABC Transporter | ABCB | ABC transporter B family member 1 |
| | Glyma.12G053800 | Isoflavone Secretion | GmICHG | isoflavone conjugate-specific beta-glucosidase |
| | Glyma.07G001300 | Soyasaponin Biosynthesis | GmBAS1 | beta-amyrin synthase |
| | Glyma.07G254600 | Soyasaponin Biosynthesis | UGT73F2, UGT73F4 | glucosyltransferase |
| | Glyma.08G181000 | Soyasaponin Biosynthesis | UGT91H4 | soyasaponin III rhamnosyltransferase |
| | Glyma.08G238100 | Soyasaponin Biosynthesis | CYP72A61 | Hydroxylates the C-22 of β-amyrin or other intermediates |
| | Glyma.08G350800 | Soyasaponin Biosynthesis | CYP93E1 | Hydroxylates the C-24 of β-amyrin or other intermediates |
| | Glyma.10G104700 | Soyasaponin Biosynthesis | UGT91H9 | putative glycosyltransferase UGT91H9 |
| | Glyma.11G053400 | Soyasaponin Biosynthesis | UGT73P2 | soyasapogenol B glucuronide galactosyltransferase |
| | Glyma.15G243300 | Soyasaponin Biosynthesis | CYP72A69 | Hydroxylates the C-21 of β-amyrin or other intermediates |
| | Glyma.16G033700 | Soyasaponin Biosynthesis | UGT73K | UDP-glycosyltransferase UGT73K |
| **Other** | Glyma.13G275400 | MATE Transporter | GmMATE76 | NA |
| | Glyma.16G219400 | Isoflavone Biosynthesis | GmCHR3 | chalcone reductase CHR3 |
| | Glyma.16G175300 | Isoflavone Biosynthesis | UGT2, GmIF7GT2 | isoflavone 7-O-glucosyltransferase |
| | Glyma.20G241600 | Isoflavone Biosynthesis | CHI1B1, CHI2, CHI2-B | chalcone--flavonone isomerase 1B-1 |
